# The genetics of the mood disorder spectrum: genome-wide association analyses of over 185,000 cases and 439,000 controls

**DOI:** 10.1101/383331

**Authors:** Jonathan R. I. Coleman, Héléna A. Gaspar, Julien Bryois, Bipolar Disorder Working Group of the Psychiatric Genomics Consortium, Major Depressive Disorder Working Group of the Psychiatric Genomics Consortium, Gerome Breen

## Abstract

**Background:** Mood disorders (including major depressive disorder and bipolar disorder) affect 10-20% of the population. They range from brief, mild episodes to severe, incapacitating conditions that markedly impact lives. Despite their diagnostic distinction, multiple approaches have shown considerable sharing of risk factors across the mood disorders.

**Methods:** To clarify their shared molecular genetic basis, and to highlight disorder-specific associations, we meta-analysed data from the latest Psychiatric Genomics Consortium (PGC) genome-wide association studies of major depression (including data from 23andMe) and bipolar disorder, and an additional major depressive disorder cohort from UK Biobank (total: 185,285 cases, 439,741 controls; non-overlapping N = 609,424).

**Results:** Seventy-three loci reached genome-wide significance in the meta-analysis, including 15 that are novel for mood disorders. More genome-wide significant loci from the PGC analysis of major depression than bipolar disorder reached genome-wide significance. Genetic correlations revealed that type 2 bipolar disorder correlates strongly with recurrent and single episode major depressive disorder. Systems biology analyses highlight both similarities and differences between the mood disorders, particularly in the mouse brain cell types implicated by the expression patterns of associated genes. The mood disorders also differ in their genetic correlation with educational attainment – positive in bipolar disorder but negative in major depressive disorder.

**Conclusions:** The mood disorders share several genetic associations, and can be combined effectively to increase variant discovery. However, we demonstrate several differences between these disorders. Analysing subtypes of major depressive disorder and bipolar disorder provides evidence for a genetic mood disorders spectrum.

## Introduction

Mood disorders affect 10-20% of the global population across their lifetime, ranging from brief, mild episodes to severe, incapacitating conditions that markedly impact lives (1–4). Major depressive disorder and bipolar disorder are the most common forms and have been grouped together since the third edition of the Diagnostic and Statistical Manual of Mental Disorders (DSM-III) (5). Although given dedicated chapters in DSM5, they remain grouped as mood disorders in the International Classification of Disorders (ICD11) (6, 7).

Depressive episodes are common to major depressive disorder and type 2 bipolar disorder, and are usually present in type 1 bipolar disorder (7). The bipolar disorders are distinguished from major depressive disorder by the presence of mania in type 1 and hypomania in type 2 (7). However, these distinctions are not absolute – some individuals with major depressive disorder may later develop bipolar disorder, and some endorse (hypo)manic symptoms (8–10). Following their first depressive episode, a non-remitting individual might develop recurrent major depressive disorder or bipolar disorder. Treatment guidelines for these disorders differ (11, 12). Identifying shared and distinct genetic associations for major depressive disorder and bipolar disorder could aid our understanding of these diagnostic trajectories.

Twin studies suggest that 35-45% of variance in risk for major depressive disorder and 65-70% of the variance in bipolar disorder risk is accounted for by additive genetic factors (13). These genetic components are partially shared, with a twin genetic correlation (r_g_) of ∼65%, and common variant based r_g_ derived from the results of genome-wide association studies (GWAS) of 30-35% (14–17). Considerable progress has been made in identifying specific genetic variants that underlie genetic risk. Recently, the Psychiatric Genomics Consortium (PGC) published a GWAS of bipolar disorder, including over 20,000 cases, with 30 genomic loci reaching genome-wide significance (16). They also performed a GWAS of major depression, including over 135,000 individuals with major depressive disorder and other definitions of depression, with 44 loci reaching genome-wide significance (15). The PGC GWAS of major depression has since been combined with a broad depression GWAS (Supplementary Note).

GWAS have identified statistical associations with major depressive disorder and with bipolar disorder individually, but have not explored the genetic aspects of the relationship between these disorders. In addition, both major depressive disorder and bipolar disorder exhibit considerable clinical heterogeneity and can be separated into subtypes. For example, the DSM5 includes categories for bipolar disorder type 1 and type 2, and for single episode and recurrent major depressive disorder (7). We use the PGC analyses of major depression and bipolar disorder, along with analyses of formally-defined major depressive disorder from UK Biobank, to explore two aims (18, 19). Firstly, we seek to identify shared and distinct mood disorder genetics by combining studies of major depressive disorder and bipolar disorder. We then explore the genetic relationship of mood disorders to traits from the wider GWAS literature. Secondly, we assess the overall genetic similarities and differences of bipolar disorder subtypes (from the PGC) and major depressive disorder subtypes (from UK Biobank), through comparing their genetic correlations and polygenic risk scores from GWAS.

## Methods and Methods

### Participants

Our primary aim was to combine analyses of bipolar disorder and major depression to examine the shared and distinct genetics of these disorders. Summary statistics were derived from participants of Western European ancestries. Full descriptions of each study and their composite cohorts are provided in each paper (15, 16, 19). Brief descriptions are provided in the Supplementary Methods. Except where otherwise specified, summary statistics are available (or will be made available) at https://www.med.unc.edu/pgc/results-and-downloads.

Major depression data were drawn from the full cohort (PGC MDD: 135,458 cases, 344,901 controls) from (15). This included data from 23andMe (20), access to which requires a Data Transfer Agreement; consequently, the data analysed here differ from the summary statistics available at the link above. Data for bipolar disorder were drawn from the discovery analysis previously reported (PGC BD: 20,352 cases, 31,358 controls), not including replication results (16).

Secondly, we wished to examine genetic correlations between mood disorder subtypes. Summary statistics were available for the primary bipolar disorder subtypes, type 1 bipolar disorder (BD1: 14,879 cases, 30,992 controls) and type 2 bipolar disorder (BD2: 3,421 cases, 22,155 controls), and for schizoaffective bipolar disorder (SAB: 977 cases, 8,690 controls), a mood disorder including psychotic symptoms. Controls are shared across these subtype analyses.

Subtype GWAS are not yet available from PGC MDD. As such, a major depressive disorder cohort was derived from the online mental health questionnaire in the UK Biobank (UKB MDD: 29,475 cases, 63,482 controls; Resource 22 on http://biobank.ctsu.ox.ac.uk) (18). The definition of major depressive disorder in this cohort is based on DSM-5, as described in full elsewhere (18), and in Supplementary Table 1 (7). We defined three major depressive disorder subtypes for analysis. Individuals meeting criteria for major depressive disorder were classed as recurrent cases if they reported multiple depressed periods across their lifetime (rMDD, N = 17,451), and single-episode cases otherwise (sMDD, N = 12,024, Supplementary Table 1). Individuals reporting depressive symptoms, but not meeting case criteria, were excluded from the main analysis but used as a “sub-threshold depression” subtype to examine the continuity of genetic associations with major depressive disorder below clinical thresholds (subMDD, N = 21,596). All subtypes were analysed with the full set of controls. Details on the quality control and analysis of the UK Biobank phenotypes is provided in the Supplementary Methods.

### Meta-analysis of GWAS data

We meta-analysed PGC MDD and UKB MDD to obtain a single major depressive disorder GWAS (combined MDD). We meta-analysed combined MDD with PGC BD, comparing mood disorder cases to controls (MOOD). Further meta-analyses were performed between PGC MDD and each bipolar disorder subtype and major depressive disorder subtype to assess the relative increase in variant discovery when adding different mood disorder definitions to PGC MDD (Supplementary Results).

Summary statistics were limited to common variants (MAF > 0.05; Supplementary Methods) either genotyped or imputed with high confidence (INFO score > 0.6) in all studies. Controls were shared between PGC MDD and PGC BD, and (due to the inclusion of summary data in PGC MDD) the extent of this overlap was unknown. Meta-analyses were therefore performed in METACARPA, which controls for sample overlap of unknown extent between studies using the variance-covariance matrix of the observed effect sizes at each variant, weighted by the sample sizes (21, 22). METACARPA adjusted adequately for known overlap between cohorts (Supplementary Methods). For later analyses (particularly linkage disequilibrium score regression) we used as the sample size a “non-overlapping N” estimated for each meta-analysis (Supplementary Methods). The definition, annotation and visualisation of each meta-analysis is described in the Supplementary Materials.

### Sensitivity analysis using down-sampled PGC MDD

Results from MOOD showed greater similarity to PGC MDD than to PGC BD. Cross-trait meta-analyses may be biased if the power of the composite analyses differs substantially (23, 24). The mean chi-square of combined MDD [1.7] exceeded that of PGC BD [1.39], suggesting this bias may affect our results (Supplementary Table 2). We therefore repeated our analyses, meta-analysing UKB MDD with summary statistics for PGC MDD that did not include participants from 23andMe nor the UK Biobank (mean chi-square = 1.35). All analyses were performed on the full and the down-sampled analyses, with the exception of GSMR analyses. Full results of the down-sampled analyses are described in the Supplementary Materials.

### Estimation of SNP-based heritability captured by common variants and genetic correlations with published GWAS

The SNP-based heritability captured by common variants was assessed using linkage disequilibrium score regression (LDSC) for each meta-analysed set of data (25). SNP-based heritability estimates were transformed to the liability scale, assuming population prevalences of 15% for combined MDD, 1% for PGC BD, and 16% for MOOD, and lower and upper bounds of these prevalences for comparison (Supplementary Methods). LDSC separates genome-wide inflation into components due to polygenicity and confounding (25). Inflation not due to polygenicity was quantified as (intercept-1)/(mean observed chi-square-1) (26). Genetic correlations were calculated in LDSC between each analysis and 414 traits curated from published GWAS. Local estimates of SNP-based heritability and genetic covariance were obtained using HESS v0.5.3b (Supplementary Methods and Results) (27, 28).

### Genetic correlations between subtype analyses

To assess the structure of genetic correlations within the mood disorders, SNP-based heritabilities and genetic correlations were calculated in LDSC between bipolar disorder subtypes (BD1, BD2, SAB), and major depressive disorder subtypes (rMDD, sMDD, subMDD). Putative differences between genetic correlations were identified using a z-test (p < 0.05), and formally tested by applying a block-jackknife, with Bonferroni correction for significance (p < 8.3×10^−4^; Supplementary Methods). Differences between the genetic correlations of PGC MDD and each bipolar disorder subtype, and between PGC BD and each major depressive disorder subtype were also tested (Bonferroni correction for significance, p < 0.0083). Genetic correlations were hierarchically clustered using the gplots package in R v1.4.1 (29, 30). Hierarchical clustering was performed using just the subtypes, and including results from six external GWAS relevant to mood disorders (Supplementary Methods). To validate our conclusion of a genetic mood disorder spectrum, we performed principal component analysis of the genetic correlation matrix including the six external GWAS (Supplementary Methods and Results).

### Association of PGC BD polygenic risk scores with major depressive disorder subtypes

Polygenic risk score analyses were performed using PRSice2 to assess whether rMDD was genetically more similar to PGC BD than were sMDD or subMDD (Supplementary Methods) (36).

### Gene-wise, gene-set, and tissue and single-cell enrichment analyses

For all analyses, gene-wise p-values were calculated as the aggregate of the mean and smallest p-value of SNPs annotated to Ensembl gene locations using MAGMA v1.06 (Supplementary Methods and Results) (37). Gene set analysis was performed in MAGMA (Supplementary Methods and Results). Further analyses were performed to assess the enrichment of associated genes with expression-specificity profiles from tissues (Genotype-Tissue Expression project, version 7) and broadly-defined (“level 1”) and narrowly-defined (“level 2”) mouse brain cell-types (38, 39). Analyses were performed in MAGMA following previously described methods with minor modifications, with Bonferroni-correction for significance (Supplementary Methods) (38). Similar analyses can be performed in LDSC-SEG – we report MAGMA results, which reflect specificity of expression across the range, whereas LDSC-SEG compares the top 10% of the range with the remainder (40). Results using LDSC are included in the Supplementary Tables.

### Mendelian randomisation (GSMR)

Bidirectional Mendelian randomisation analyses were performed using the GSMR option in GCTA to allow exploratory inference of the causal direction of known relationships between mood disorder traits and other traits (41, 42). Specifically, the relationship between the mood disorder analyses (MOOD, combined MDD, PGC BD) and schizophrenia, intelligence, educational attainment, body mass index, and coronary artery disease were explored (Supplementary Methods) (32, 43–46). These traits were previously examined in the PGC major depression GWAS – we additionally tested intelligence following the results of our genetic correlation analyses (15).

### Conditional and reversed-effect analyses

Additional analyses were performed to identify shared and distinct mood disorder loci, using mtCOJO, an extension of GSMR (Supplementary Methods) (41, 42). Analyses were performed on combined MDD conditional on PGC BD, and on PGC BD conditional on combined MDD (Supplementary Results). To identify loci with opposite directions of effect between combined MDD and PGC BD, the MOOD meta-analysis was repeated with reversed direction of effects for PGC BD (Supplementary Methods and Results).

## Results

### Evidence for confounding in meta-analyses

Meta-analysis results were assessed for genome-wide inflation of test statistics using LDSC (25). The LDSC intercept was significantly >1 in most cases (1.00-1.06), which has previously been interpreted as confounding (Supplementary Table 2). However, such inflation can occur in large cohorts without confounding (47). Estimates of inflation not due to polygenicity were small in all meta-analyses (4-7%, Supplementary Table 2).

### Combined MOOD meta-analysis

We meta-analysed the PGC MDD, PGC BD and the UKB MDD cohorts (MOOD, cases = 185,285, controls = 439,741, non-overlapping N = 609,424). 73 loci reached genome-wide significance, of which 55 were also seen in the meta-analysis of PGC MDD and UKB MDD (combined MDD, Supplementary Table 3, Supplementary Figures 1 and 2). Results are summarised in Table 1: 39 of the 44 PGC MDD loci reached genome-wide significance in MOOD (Supplementary Table 3, Supplementary Figures 1-8). In comparison, only four of the 19 PGC BD loci reached genome-wide significance in MOOD. MOOD loci overlapped considerably with previous studies of depression and depressive symptoms (51/73) (20, 23, 48– 52), bipolar disorder (3/73) (53–56), neuroticism (32/73) (23, 57–59), and schizophrenia (15/73) (32, 60), although participants overlap between MOOD and many of these studies. Locus 52 (chromosome 12) passed genome-wide significance in a previous meta-analysis of broad depression and bipolar disorder, although the two other loci from this study did not replicate (51). Six of the 73 associations are entirely novel (p > 5×10^−8^ in previous studies of all phenotypes; Table 1, Supplementary Table 4).

**Table 1:** Loci genome-wide significant (p < 5×10^−8^) in the MOOD meta-analysis. Locus – shared locus number for annotation (Supplementary Table 3), Chr – chromosome, BP – base position, A1 – effect allele, A2 – non-effect allele, Previous report – locus previously implicated in PGC MDD (D), PGC BD (B), previous combined studies of bipolar disorder and major depressive disorder (BD), other studies of major depressive disorder or depressive symptoms (DO), other studies of bipolar disorder (BO), previous combined studies of bipolar disorder and schizophrenia (BS), previous combined studies of major depressive disorder and schizophrenia (DS), neuroticism (N), schizophrenia (S), or other studies (O – see Supplementary Table 4).

| Locus | Chr | BP | Index SNP | A1 | A2 | OR | SE | p | Previous report |
| --- | --- | --- | --- | --- | --- | --- | --- | --- | --- |
| 1 | 1 | 37192741 | rs1002656 | T | C | 0.97 | 0.005 | $2.71 \times 10^{-11}$ | DO, N |
| 2 | 1 | 72837239 | rs7531118 | T | C | 0.96 | 0.004 | $1.05 \times 10^{-16}$ | D, DO, S, O |
| 4 | 1 | 80795989 | rs6667297 | A | G | 0.97 | 0.005 | $5.86 \times 10^{-11}$ | D, DO |
| 5 | 1 | 90796053 | rs4261101 | A | G | 0.97 | 0.005 | $1.78 \times 10^{-8}$ | D |
| 6 | 1 | 175913828 | rs10913112 | T | C | 0.97 | 0.005 | $1.46 \times 10^{-10}$ | DO, O |
| 7 | 1 | 177370033 | rs16851203 | T | C | 0.96 | 0.007 | $2.38 \times 10^{-9}$ | DO, S, O |
| 9 | 2 | 22582968 | rs61533748 | T | C | 0.97 | 0.004 | $3.84 \times 10^{-11}$ | DO, N |
| 10 | 2 | 57987593 | rs11682175 | T | C | 0.97 | 0.004 | $2.18 \times 10^{-11}$ | D, DO, BS, N, S, O |
| 11 | 2 | 157111313 | rs1226412 | T | C | 1.03 | 0.005 | $1.27 \times 10^{-8}$ | D, DO, N, O |
| 12 | 2 | 198807015 | rs1518367 | A | T | 0.97 | 0.005 | $1.18 \times 10^{-8}$ | BS, S, O |
| 13 | 3 | 108148557 | rs1531188 | T | C | 0.96 | 0.006 | $1.61 \times 10^{-9}$ | O |
| 14 | 3 | 158107180 | rs7430565 | A | G | 0.97 | 0.004 | $2.30 \times 10^{-11}$ | D, DO, N, O |
| 16 | 4 | 42047778 | rs34215985 | C | G | 0.97 | 0.006 | $1.72 \times 10^{-10}$ | D, DO, N |
| 17 | 5 | 77709430 | rs4529173 | T | C | 0.97 | 0.005 | $4.29 \times 10^{-9}$ | O |
| 18 | 5 | 88002653 | rs447801 | T | C | 1.03 | 0.004 | $2.29 \times 10^{-10}$ | D, DO, N, O |
| 19 | 5 | 92995013 | rs71639293 | A | G | 1.03 | 0.005 | $5.85 \times 10^{-9}$ | DO, N |
| 20 | 5 | 103904226 | rs12658032 | A | G | 1.04 | 0.005 | $2.19 \times 10^{-16}$ | D, DO, N, O |
| 21 | 5 | 106603482 | rs55993664 | A | C | 0.97 | 0.006 | $1.87 \times 10^{-8}$ | NOVEL LOCUS |
| 22 | 5 | 124251883 | rs116755193 | T | C | 0.97 | 0.005 | $1.47 \times 10^{-10}$ | D, O |
| 23 | 5 | 164523472 | rs11135349 | A | C | 0.97 | 0.004 | $2.96 \times 10^{-11}$ | D, DO, N |
| 24 | 5 | 166992078 | rs4869056 | A | G | 0.97 | 0.005 | $5.21 \times 10^{-9}$ | D |
| 25 | 6 | 28673998 | rs145410455 | A | G | 0.94 | 0.007 | $7.17 \times 10^{-18}$ | D, DO, BO, BS, DS, N, S, O |
| 26 | 6 | 101339400 | rs7771570 | T | C | 0.97 | 0.004 | $9.68 \times 10^{-10}$ | DO, N, O |
| 27 | 6 | 105365891 | rs1933802 | C | G | 0.98 | 0.004 | $1.05 \times 10^{-8}$ | DO, S, O |
| 28 | 7 | 12267221 | rs4721057 | A | G | 0.97 | 0.004 | $7.31 \times 10^{-11}$ | D, DO, N, O |
| 29 | 7 | 24826589 | rs79879286 | C | G | 1.04 | 0.006 | $1.97 \times 10^{-11}$ | B, BS, DO, S |
| 30 | 7 | 82514089 | rs34866621 | T | C | 1.03 | 0.005 | $2.21 \times 10^{-8}$ | DO, O |
| 31 | 7 | 109099919 | rs58104186 | A | G | 1.03 | 0.004 | $7.12 \times 10^{-9}$ | D, DO |
| 34 | 9 | 11379630 | rs10959753 | T | C | 0.96 | 0.005 | $1.45 \times 10^{-13}$ | D, DO, N, O |
| 35 | 9 | 37207269 | rs4526442 | T | C | 0.96 | 0.006 | $7.97 \times 10^{-11}$ | DO, O |
| 36 | 9 | 81413414 | rs11137850 | A | G | 1.03 | 0.005 | $1.25 \times 10^{-8}$ | NOVEL LOCUS |
| 38 | 9 | 119733380 | rs10759881 | A | C | 1.03 | 0.005 | $8.56 \times 10^{-9}$ | D, DO |
| 40 | 9 | 122664468 | rs10818400 | T | G | 0.98 | 0.004 | $1.29 \times 10^{-8}$ | N |
| 41 | 9 | 126682068 | rs7029033 | T | C | 1.04 | 0.008 | $2.61 \times 10^{-8}$ | D, DO, O |
| 42 | 10 | 104684544 | rs78821730 | A | G | 0.96 | 0.007 | $2.95 \times 10^{-8}$ | N, BS, S, O |
| 43 | 10 | 106563924 | rs61867293 | T | C | 0.96 | 0.005 | $5.64 \times 10^{-12}$ | D, DO, N, O |
| 44 | 11 | 16293680 | rs977509 | T | C | 0.97 | 0.005 | $1.19 \times 10^{-8}$ | DO, N, O |
| 45 | 11 | 31850105 | rs1806153 | T | G | 1.03 | 0.005 | $2.81 \times 10^{-9}$ | D, DO, N, O |
| 46 | 11 | 32765866 | rs143864773 | T | C | 1.04 | 0.008 | $1.70 \times 10^{-8}$ | NOVEL LOCUS |
| 47 | 11 | 61557803 | rs102275 | T | C | 0.97 | 0.005 | $5.04 \times 10^{-11}$ | B, DO, BO, O |
| 48 | 11 | 63632673 | rs10792422 | T | G | 0.98 | 0.004 | $2.18 \times 10^{-8}$ | O |
| 49 | 11 | 88743208 | rs4753209 | A | T | 0.97 | 0.004 | $4.15 \times 10^{-9}$ | DO, N, O |
| 50 | 11 | 99268617 | rs1504721 | A | C | 0.98 | 0.004 | $2.24 \times 10^{-8}$ | O |
| 51 | 11 | 113392994 | rs2514218 | T | C | 0.97 | 0.005 | $3.22 \times 10^{-10}$ | DO, BS, N, S, O |
| 52 | 12 | 2344644 | rs769087 | A | G | 1.03 | 0.005 | $3.27 \times 10^{-8}$ | B, BD, BO, DS,<br>BS, S, O |
| 53 | 12 | 23947737 | rs4074723 | A | C | 0.97 | 0.004 | $3.18 \times 10^{-9}$ | D, DO, N, O |
| 54 | 12 | 121186246 | rs58235352 | A | G | 0.95 | 0.009 | $1.64 \times 10^{-10}$ | DO, O |
| 55 | 12 | 121907336 | rs7962128 | A | G | 1.02 | 0.004 | $3.63 \times 10^{-8}$ | NOVEL LOCUS |
| 56 | 13 | 44327799 | rs4143229 | A | C | 0.95 | 0.008 | $2.73 \times 10^{-10}$ | D |
| 57 | 13 | 53625781 | rs12552 | A | G | 1.04 | 0.004 | $1.25 \times 10^{-23}$ | D, DO, O |
| 58 | 14 | 42074726 | rs61990288 | A | G | 0.97 | 0.004 | $2.29 \times 10^{-10}$ | D, DO, O |
| 60 | 14 | 64686207 | rs915057 | A | G | 0.98 | 0.004 | $1.92 \times 10^{-8}$ | D, DO, O |
| 61 | 14 | 75130235 | rs1045430 | T | G | 0.97 | 0.004 | $9.83 \times 10^{-11}$ | D, DO, N, O |
| 62 | 14 | 104017953 | rs10149470 | A | G | 0.97 | 0.004 | $1.15 \times 10^{-10}$ | D, DS, DO, BS, S,<br>O |
| 63 | 15 | 36355868 | rs1828385 | A | C | 0.97 | 0.004 | $1.15 \times 10^{-8}$ | NOVEL LOCUS |
| 64 | 15 | 37643831 | rs8037355 | T | C | 0.97 | 0.004 | $4.09 \times 10^{-15}$ | D, DO, O |
| 65 | 16 | 6310645 | rs8063603 | A | G | 0.97 | 0.005 | $5.36 \times 10^{-11}$ | D, DO |
| 66 | 16 | 7667332 | rs11077206 | C | G | 1.03 | 0.004 | $5.49 \times 10^{-10}$ | D, DO, N, O |
| 67 | 16 | 13038723 | rs12935276 | T | G | 0.97 | 0.005 | $4.75 \times 10^{-10}$ | D, DO, N, O |
| 68 | 16 | 13750257 | rs7403810 | T | G | 1.03 | 0.005 | $7.52 \times 10^{-11}$ | DO, BS, S, O |
| 69 | 16 | 72214276 | rs11643192 | A | C | 1.03 | 0.004 | $1.46 \times 10^{-11}$ | D, O |
| 70 | 17 | 27363750 | rs75581564 | A | G | 1.04 | 0.006 | $2.47 \times 10^{-10}$ | D, DO, O |
| 71 | 18 | 31349072 | rs4534926 | C | G | 1.03 | 0.004 | $9.14 \times 10^{-9}$ | DO, N |
| 72 | 18 | 36883737 | rs62099069 | A | T | 0.97 | 0.004 | $9.52 \times 10^{-10}$ | D, O |
| 73 | 18 | 42260348 | rs117763335 | T | C | 0.97 | 0.005 | $1.33 \times 10^{-8}$ | O |
| 74 | 18 | 50614732 | rs11663393 | A | G | 1.03 | 0.004 | $1.56 \times 10^{-10}$ | D, DO, N, O |
| 75 | 18 | 52517906 | rs1833288 | A | G | 1.03 | 0.005 | $4.54 \times 10^{-8}$ | D, DS, DO, N, S,<br>O |
| 76 | 18 | 53101598 | rs12958048 | A | G | 1.04 | 0.005 | $4.86 \times 10^{-14}$ | D, DO, BS, N, S, O |
| 77 | 19 | 30939989 | rs33431 | T | C | 1.02 | 0.004 | $4.04 \times 10^{-8}$ | DO, O |
| 78 | 20 | 45841052 | rs910187 | A | G | 0.97 | 0.005 | $3.09 \times 10^{-9}$ | DO, O |
| 79 | 22 | 41621714 | rs2179744 | A | G | 1.03 | 0.005 | $3.83 \times 10^{-12}$ | D, B, DO, BS, N,<br>S, O |
| 80 | 22 | 42815358 | rs7288411 | A | G | 1.03 | 0.005 | $3.86 \times 10^{-8}$ | NOVEL LOCUS |
| 81 | 22 | 50679436 | rs113872034 | A | G | 0.96 | 0.006 | $1.10 \times 10^{-9}$ | O |

**Figure 1a:**
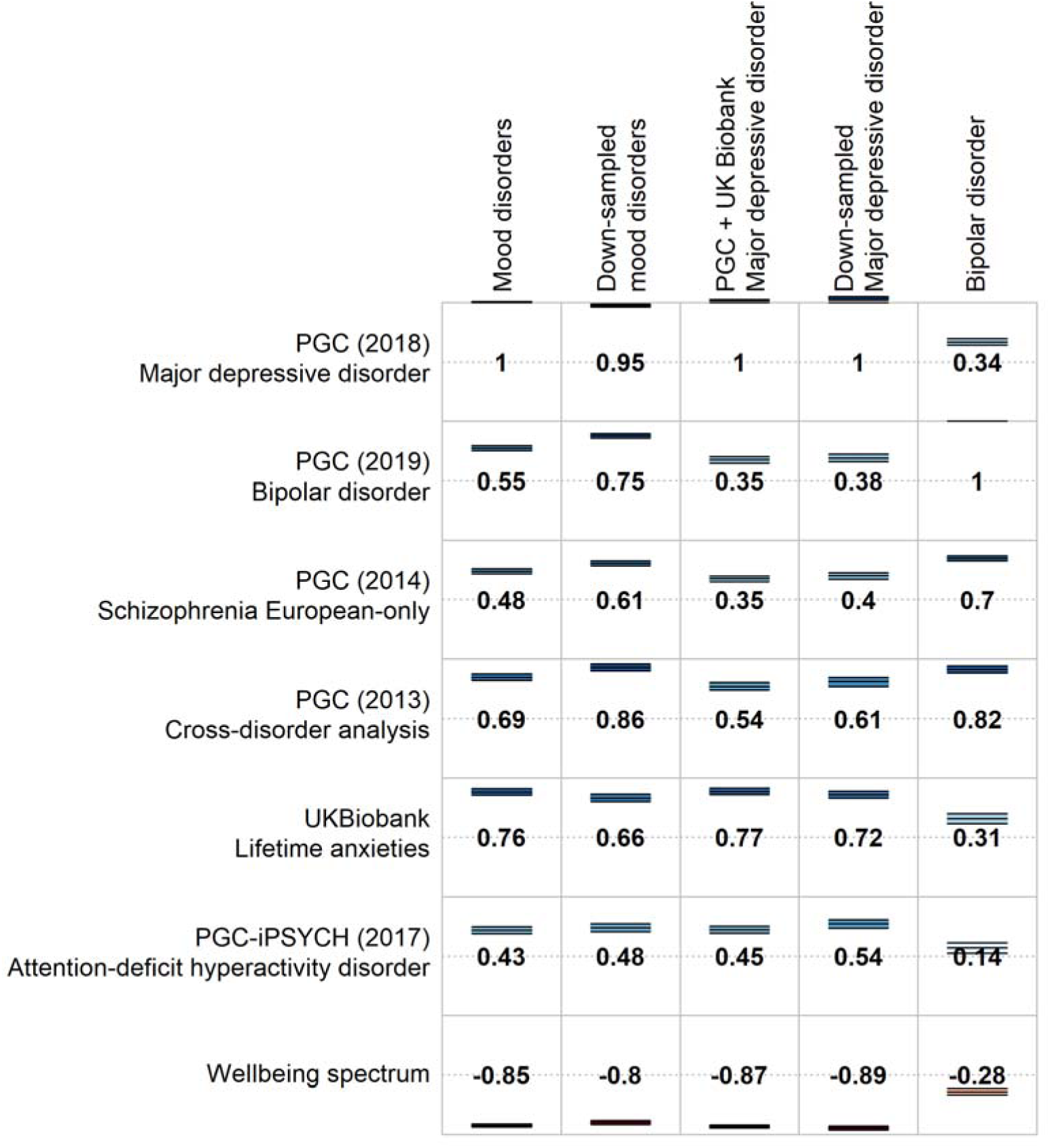
Selected genetic correlations of psychiatric traits with the main meta-analysis (MOOD), the separate mood disorder analyses (combined MDD and PGC BD), and the down-sampled analyses (down-sampled MOOD, down-sampled MDD). Full genetic correlation results are provided in Supplementary Table 5.

**Figure 1b:**
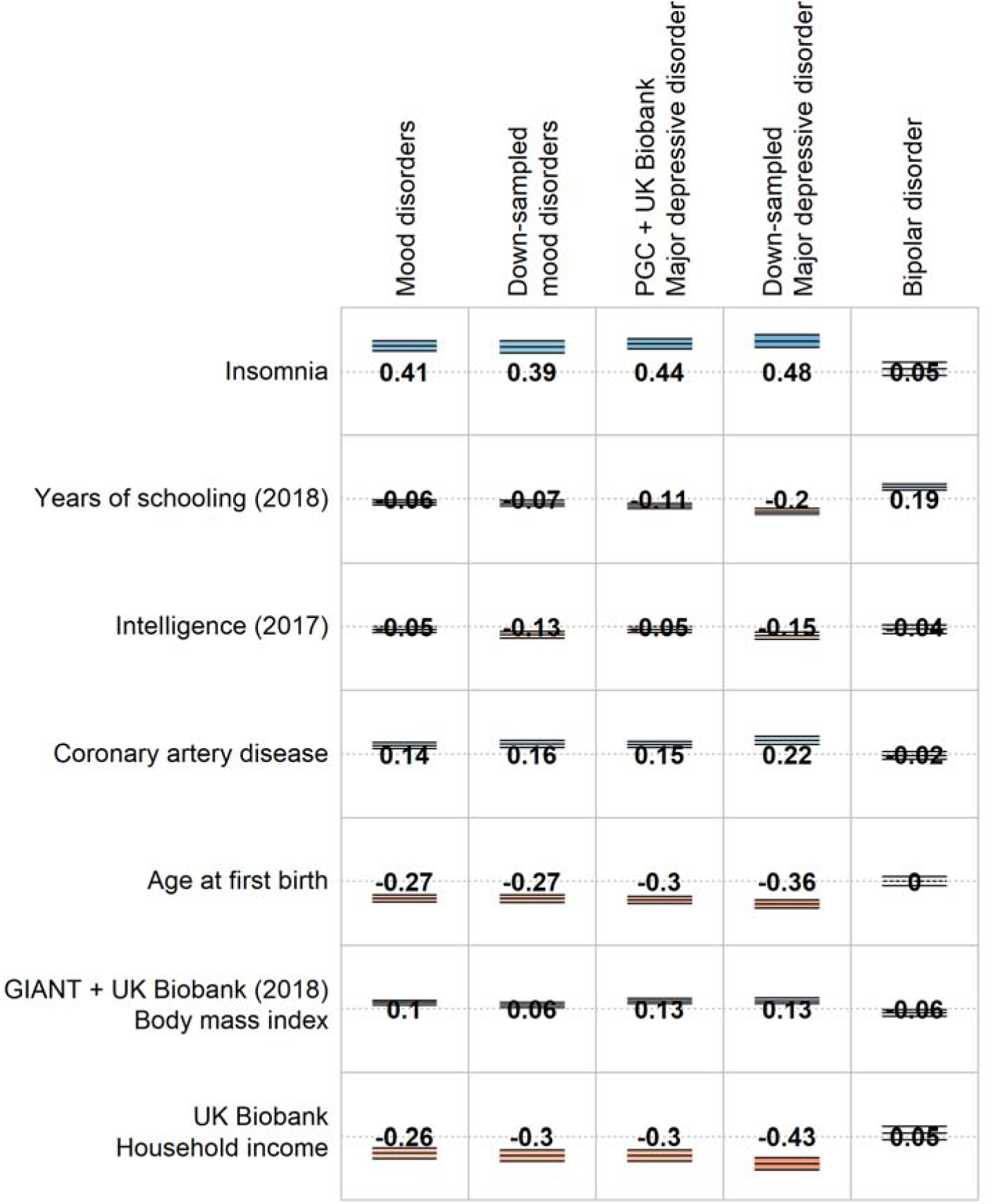
Selected genetic correlations of other traits with the main meta-analysis (MOOD), the separate mood disorder analyses (combined MDD and PGC BD), and the down-sampled analyses (down-sampled MOOD, down-sampled MDD). Full genetic correlation results are provided in Supplementary Table 5.

**Figure 2:**
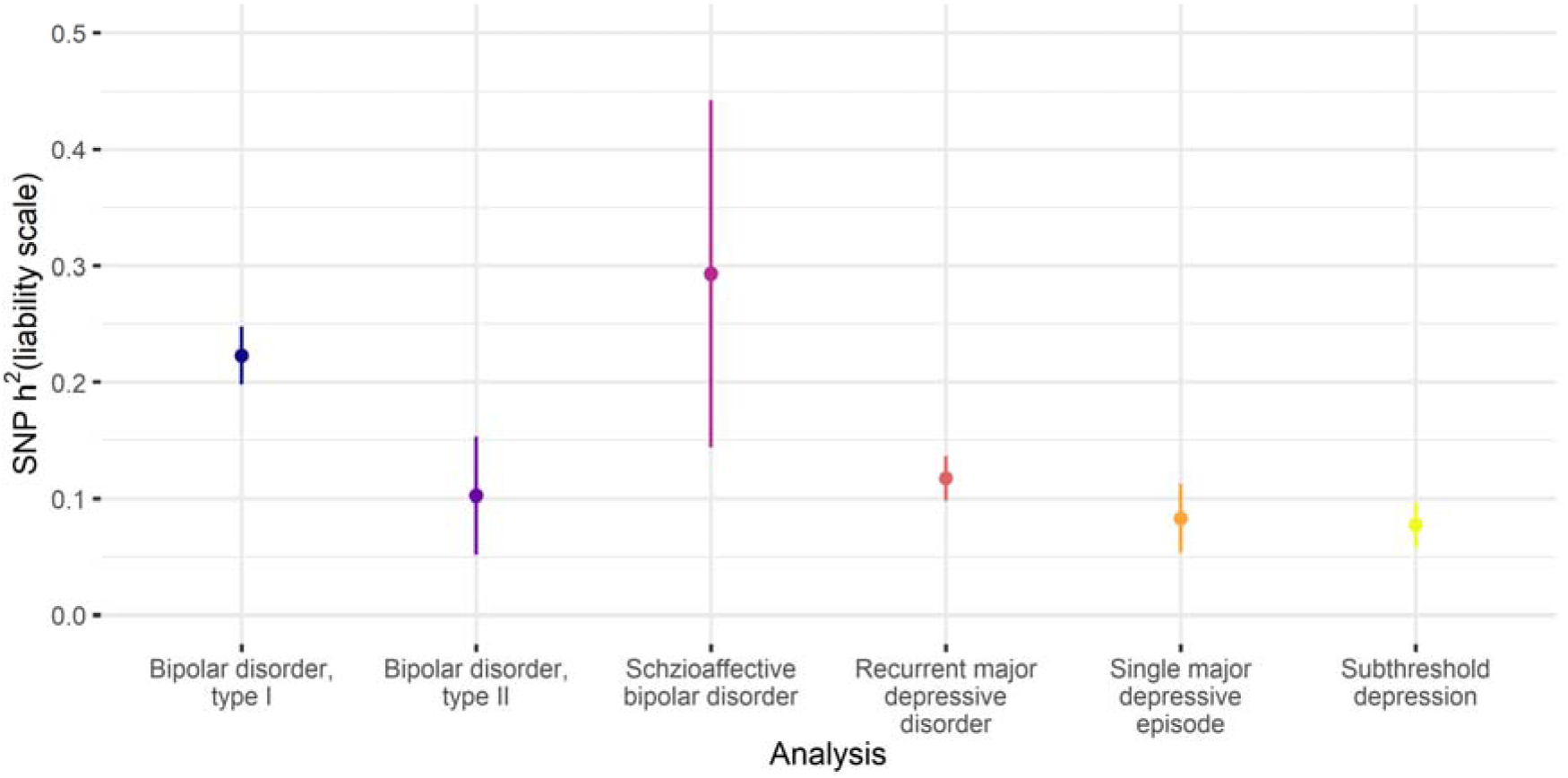
SNP-based heritability estimates for the subtypes of bipolar disorder and subtypes of major depressive disorder. Points = SNP-based heritability estimates. Lines = 95% confidence intervals. Full SNP-based heritability results are provided in Supplementary Table 2.

The down-sampled MOOD (cases = 95,481, controls = 287,932, non-overlapping N = 280,214) showed increased similarity to PGC BD compared to MOOD, but remained more similar to PGC MDD. Nineteen loci reached genome-wide significance in down-sampled MOOD, including nine (20%) from PGC MDD, compared with two (11%) reported in PGC BD (Supplementary Table 3). 17/19 loci were also observed in MOOD. Of the two loci not observed in MOOD, one passed genome-wide significance in PGC BD.

### SNP-based heritability and genetic correlations

The estimate of SNP-based heritability for MOOD (8.8%) was closer to PGC MDD (9%) than to PGC BD (17-23%) (15, 16). Significant genetic correlations between MOOD and other traits included psychiatric and behavioural, reproductive, cardiometabolic, and sociodemographic traits (Figure 1, Supplementary Table 5). Genetic correlations with psychiatric and behavioural traits are consistently observed across psychiatric traits (17, 61). The genetic correlation with educational attainment differs, being negative in combined MDD, but positive in PGC BD (Supplementary Table 6). The genetic correlation (r_g_) between MOOD and educational attainment was −0.058 (p=0.004), intermediate between the results of combined MDD and of PGC BD. Notably, the genetic correlation with intelligence (IQ) was not significant in combined MDD, PGC BD, nor MOOD (p>1.27×10^−4^). However, sensitivity analyses (see below), indicated that including 23andMe in the PGC MDD sample obscured a negative genetic correlation of MDD with IQ.

The SNP-based heritability of down-sampled MOOD from LDSC was 11%, closer to PGC MDD than to PGC BD (Supplementary Table 2). Genetic correlations varied (Supplementary Tables 5 and 7) with some more similar to PGC BD (schizophrenia: down-sampled rg = 0.61, combined MDD rg = 0.35, PGC BD rg = 0.7), and others more similar to combined MDD (ADHD: down-sampled rg = 0.48, combined MDD rg = 0.45, PGC BD rg = 0.14). The genetic correlation with IQ was significant (rg = −0.13, p = 5×10^−7^), because the excluded 23andMe depression cohort has a positive genetic correlation with IQ (rg = 0.06, p = 0.01). The greater genetic correlation of MOOD with combined MDD (0.98) compared to PGC BD (0.55) persisted when comparing down-sampled MOOD to combined MDD (0.85) and PGC BD (0.75; Supplementary Table 6).

### Relationship between mood disorder subtypes

Analyses were performed using GWAS data from subtypes of bipolar disorder (BD1, BD2, SAB) and major depressive disorder (rMDD, sMDD, subMDD). SNP-based heritability for the subtypes ranged from subMDD and sMDD (8%), through BD2 and rMDD (10% and 12%, respectively) to BD1 and SAB (22% and 29% respectively, Figure 2, Supplementary Table 2).

The major depressive disorder subtypes were strongly and significantly genetically correlated (r_g_ = 0.9-0.94, p_rg = 0_ < 8.3×10^−4^). These correlations did not differ significantly from 1 (all p_rg = 1_ > 0.3), nor from each other (all pΔrg = 0 > 0.5, Figure 2, Supplementary Table 8). BD1 and SAB were strongly correlated (r_g_ = 0.77, p_rg = 0_ = 6×10^−13^, p_rg = 1_ = 0.03), as were BD1 and BD2 (r_g_ = 0.86, p_rg = 0_ = 3×10^−16^, p_rg = 1_ = 0.2). However, BD2 was not significantly correlated with SAB (r_g_ = 0.22, p_rg = 0_ = 0.02).

In hierarchical clustering, BD2 clustered with the major depressive disorder subtypes rather than the bipolar disorder subtypes. The strength of correlation between BD2 and BD1 did not differ from that between BD2 and rMDD (r_g_ = 0.68, p_rg = 0_ = 3×10^−8^, p_rg = 1_ = 0.01), following multiple testing correction (Δr_g_ = 0.18, p = 0.02). Overall, these results suggest a spectrum of genetic relationships between major depressive disorder and bipolar disorder, with BD2 bridging the two disorders (Figure 3; Supplementary Figure 9). This spectrum remained when six external phenotypes were added, and was supported by results from principal component analysis (Supplementary Results, Supplementary Figure 10).

**Figure 3:**
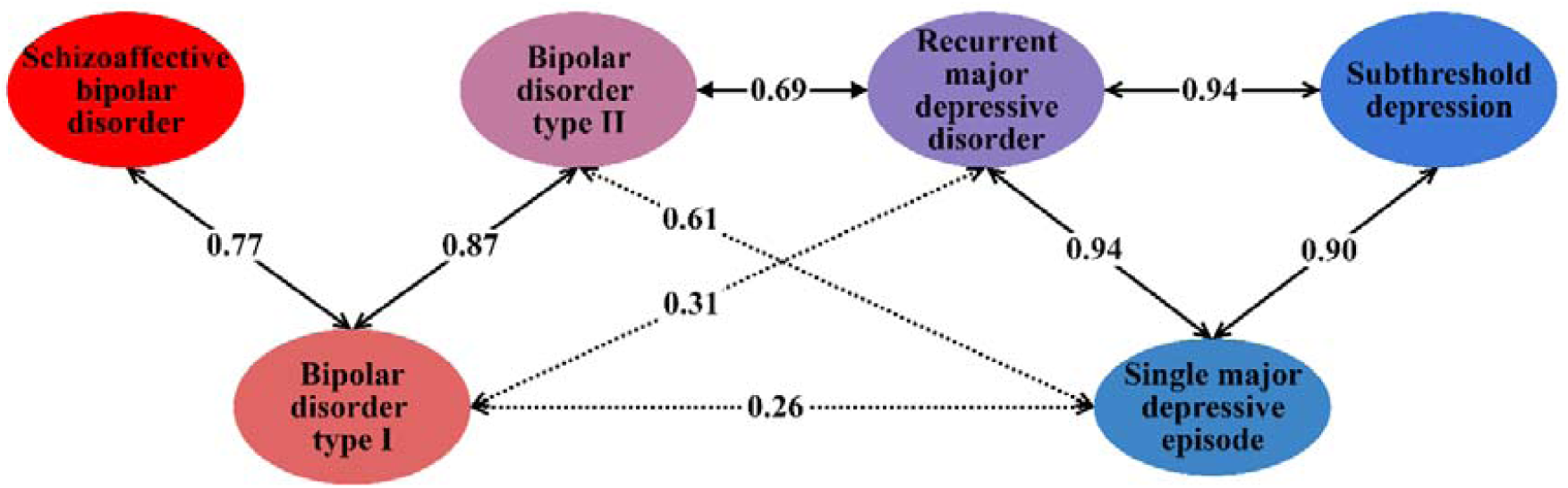
Genetic correlations across the mood disorder spectrum. Labelled arrows show genetic correlations significantly different from 0. Solid arrows represent genetic correlations not significantly different from 1 (p < 0.00333, Bonferroni correction for 15 tests). Full results are provided in Supplementary Table 8.

Polygenic risk score analyses showed that individuals with high polygenic risk scores for PGC BD were more likely to report rMDD than sMDD, and more likely to report sMDD than subMDD (Supplementary Results).

### Tissue and cell-type specificity analyses

The results of gene-wise and gene set analyses are described in the Supplementary Results. The tissue-specificity of associated genes differed minimally between the analyses (Supplementary Table 9). All brain regions were significantly enriched in all analyses, and the pituitary was also enriched in combined MDD and PGC BD (p < 9.43×10^−4^, Bonferroni correction for 53 regions, Supplementary Table 9). Results from down-sampled MOOD and down-sampled MDD were generally consistent with the main analyses, except spinal cord was not enriched in either, nor was the cordate in the down-sampled MDD analysis.

In contrast, cell-type enrichments differed between combined MDD and PGC BD (Figure 4, Supplementary Tables 10 and 11). Genes associated with PGC BD were enriched for expression in pyramidal cells from the CA1 region of the hippocampus and the somatosensory cortex, and in striatal interneurons. None of these enrichments were significant in combined MDD. Genes only associated with combined MDD were significantly enriched for expression in neuroblasts and dopaminergic neurons from adult mice. Further cell-types (dopaminergic neuroblasts; dopaminergic, GABAergic and midbrain nucleus neurons from embryonic mice; interneurons; and medium spiny neurons) were enriched for both combined MDD and PGC BD, but the rank and strength of enrichment differed, most notably for medium spiny neurons. The general pattern of differences persisted when comparing PGC BD with down-sampled MDD, although genes associated with down-sampled MDD were not enriched for expression in adult dopaminergic neurons, embryonic midbrain nucleus neurons, interneurons, nor medium spiny neurons (Supplementary Figure 11).

**Figure 4:**
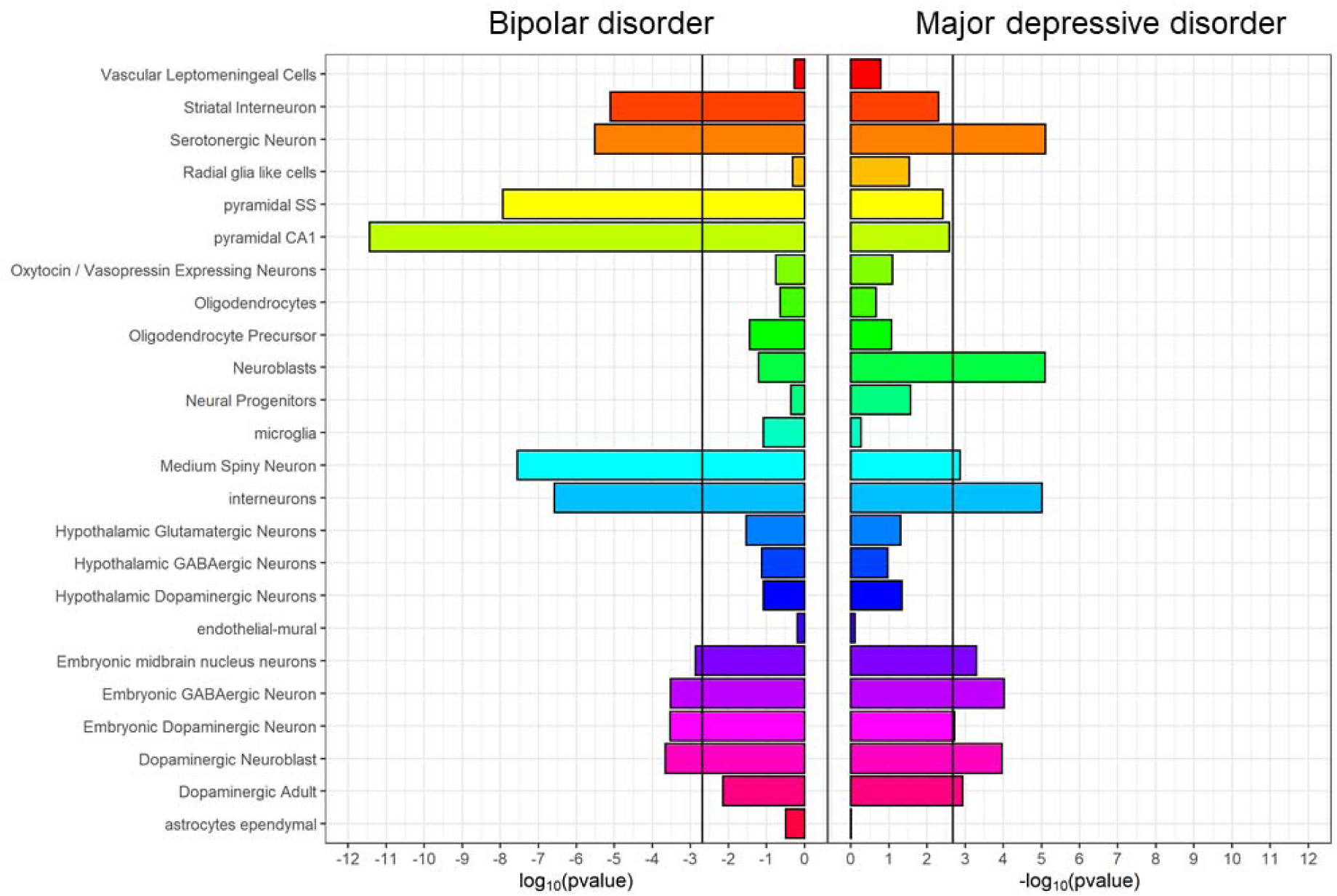
Cell-type expression specificity of genes associated with bipolar disorder (PGC BIP, left) and major depressive disorder (combined MDD, right). Black vertical lines = significant enrichment (p < 2×10^−3^, Bonferroni correction for 24 cell types). See Supplementary Table 10 for full results.

### Shared and distinct relationships with mood disorders and inferred causality

Bidirectional Mendelian randomisation was used to investigate previously-described relationships between mood disorder phenotypes (combined MDD, PGC BD) and external traits: schizophrenia, educational attainment, IQ, body mass index (BMI) and coronary artery disease (CAD; Figure 5, Supplementary Table 12). Associations with PGC BD should be interpreted cautiously, as only 19 loci reached genome-wide significance, several of which were removed as potentially pleiotropic in the analyses below.

**Figure 5:**
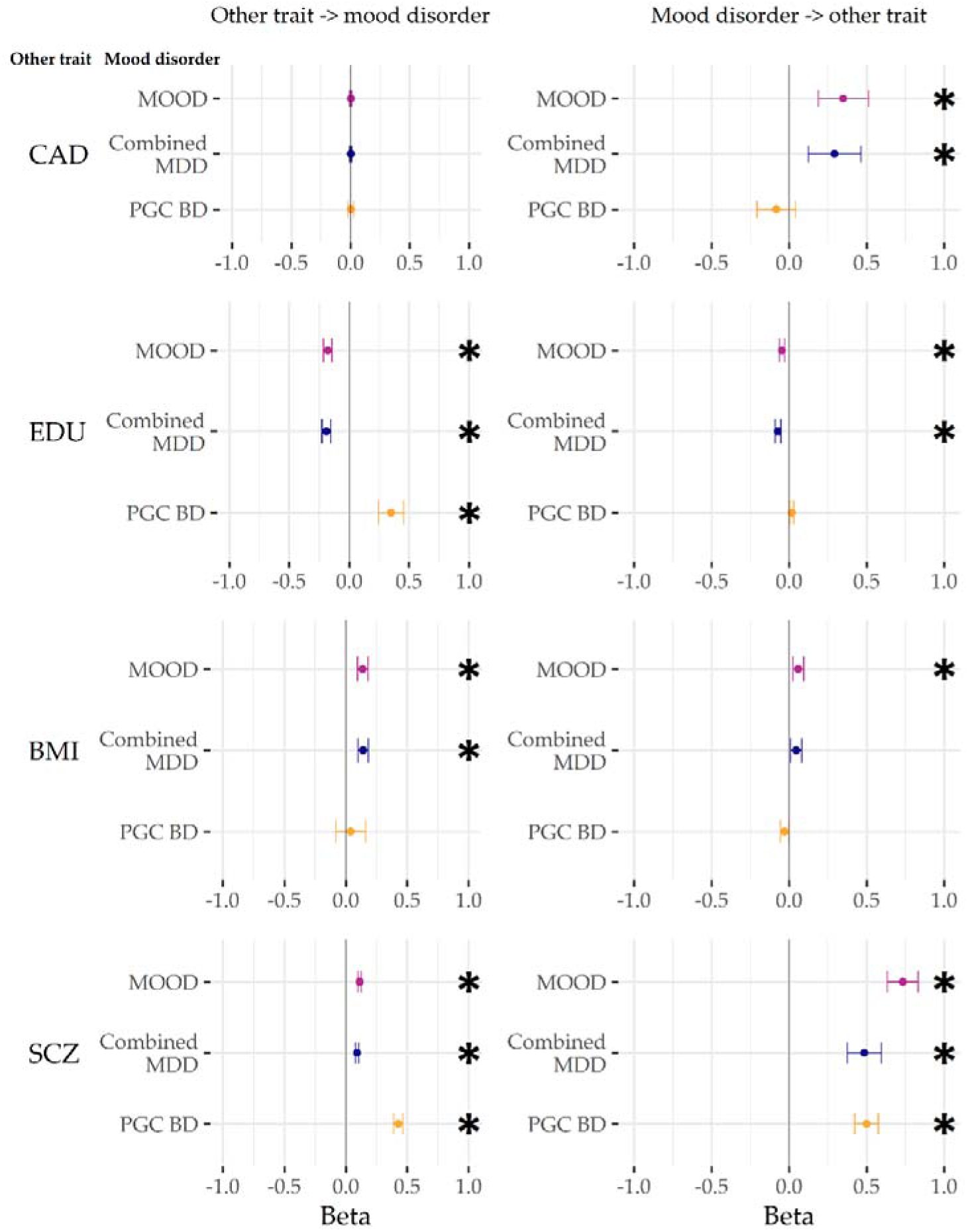
GSMR results from analyses with the main meta-analysis (MOOD), and the major depression and bipolar disorder analyses (combined MDD, PGC BD). External traits are coronary artery disease (CAD), educational attainment (EDU), body mass index (BMI), and schizophrenia (SCZ). Betas are on the scale of the outcome GWAS (logit for binary traits, phenotype scale for continuous). * p < 0.004 (Bonferroni correction for two-way comparisons with six external traits). For figure data, including the number of non-pleiotropic SNPs included in each instrument, see Supplementary Table 12.

A positive bidirectional relationship was observed between combined MDD and PGC BD, and between schizophrenia and both combined MDD and PGC BD. This is consistent with psychiatric disorders acting as causal risk factors for the development of further psychiatric disorders, or being correlated with other causal risk factors, including (but not limited to) the observed shared genetic basis.

The relationship with educational years differed between the mood disorders – there was a negative bidirectional relationship between educational years and combined MDD, but a positive bidirectional relationship with PGC BD (albeit with only nominal significance from PGC BD to educational years). In contrast, no significant relationship was observed between mood phenotypes and IQ. This is consistent with differing causal roles of education (or correlates of education) on the mood disorders, with a weaker reciprocal effect of the mood disorders altering the length of education.

A positive association was seen from BMI to combined MDD, but not from combined MDD to BMI. In contrast, only a nominally significant negative relationship was seen from PGC BD to BMI. A positive association was observed from combined MDD to CAD; no relationship was observed between CAD and PGC BD.

## Discussion

We identified 73 genetic loci by meta-analysing cohorts of major depression and bipolar disorder, including 15 loci novel to mood disorders. Our overall mood disorders meta-analysis results (MOOD) have more in common with our major depressive disorder analysis (combined MDD) than our bipolar disorder analysis (PGC BD). Partly, this results from the greater power of the major depressive disorder analysis compared to the bipolar disorder analysis. Nevertheless, genetic associations from our sensitivity analysis with equivalently powered cohorts (using down-sampled MDD in place of combined MDD) still showed a greater overall similarity to those from major depressive disorder rather than bipolar disorder.

This may reflect a complex genetic architecture in bipolar disorder, wherein one set of variants may be associated more with manic symptoms and another set with depressive symptoms. Variants associated more with mania (or psychosis) may have higher effect sizes, detectable at current bipolar disorder GWAS sample sizes, and may not be strongly associated with major depressive disorder. This could contribute to the observed higher heritability of bipolar disorder compared to major depressive disorder, and agrees with reports that most of the genetic variance for mania is not shared with depression (13, 14). In this case, meta-analysis of bipolar disorder and major depressive disorder cohorts would support variants associated more with depression, but not those associated more with mania. This is consistent with our findings, and with depressive symptoms being both the unifying feature of the mood disorders and the core feature of major depressive disorder.

We assessed genetic correlations between mood disorder subtypes. We observed high, consistent correlations between major depressive disorder subtypes, including sub-threshold depression. Bipolar disorder type 2 showed greater genetic similarity to major depressive disorder compared to type 1. In this, we build on similar findings from polygenic risk scores analyses (16, 56). Individuals with high polygenic risk scores for PGC BD were more likely to report recurrent than single-episode major depressive disorder. However, the genetic correlation of PGC BD with recurrent major depressive disorder was not significantly greater than that with single-episode major depressive disorder. This might reflect the difference in power between these methods. We also examined the genetic correlations between mood disorder subtypes in the context of relevant external traits (Supplementary Results). Our subtype analyses support a genetic mood spectrum consisting of the schizophrenia-like bipolar disorder type 1 and schizoaffective disorder at one pole, and the depressive disorders at the other, with bipolar disorder type 2 occupying an intermediate position.

Conditional and reversed-effect analyses (Supplementary Results) suggest that few of the loci we identified are disorder-specific. However, our results highlight some differences between the genetics of the mood disorders. The expression specificity of associated genes in mouse brain cell types differed between bipolar disorder and major depressive disorder analyses. Cell-types more associated with bipolar disorder (pyramidal neurons and striatal interneurons) were also enriched in analyses of schizophrenia (38). Cell-types more associated in major depressive disorder (neuroblasts, adult dopaminergic neurons, embryonic GABAergic neurons) had weaker enrichments in schizophrenia, but were enriched in analyses of neuroticism (57). The higher rank of the enrichment of serotonergic neurons with major depressive disorder compared to bipolar disorder is striking given the use of drugs targeting the serotonergic system in the treatment of depression (63). Nevertheless, cell-type enrichment analyses are still novel, and require cautious interpretation, especially given the use of non-human reference data (38, 64).

We explored potential causal relationships between the mood disorders and other traits using Mendelian randomisation. The interpretation of these analyses is challenging, especially for complex traits, when the ascertainment of cases varies, and when there are relatively few (< 20) variants used as instruments (for example, in the PGC BD and down-sampled analyses presented) (41, 67, 68). Major depressive disorder and bipolar disorder demonstrate considerable heterogeneity (as our subtype analyses show for bipolar disorder types 1 and 2), potentially confounding the results of Mendelian randomisation. That said, our analyses are consistent with a bidirectional influence of educational attainment on risk for mood disorders (and vice versa), with different directions of effect in the two mood disorders. We found no significant relationship between IQ and either mood disorder. We also find results consistent with major depressive disorder increasing the risk for coronary artery disease in a relatively well powered analysis. This mirrors epidemiological findings, although the mechanism remains unclear (69).

Despite the presence of depressive episodes, the mood disorders are diagnostically distinct. This is reflected in their differing epidemiology – for example, more women than men suffer from major depressive disorder, whereas diagnoses of bipolar disorder are roughly equal between the sexes (3). Differences in our genetic results between major depressive disorder and bipolar disorder may result from epidemiological heterogeneity, rather than distinct biological mechanisms (70). Deeper phenotyping of GWAS datasets is ongoing, and will enable the effect of confounding factors such as sex to be estimated in future studies (71).

We extend previous findings showing genetic continuity across the mood disorders (15–17, 56). Combined analyses of major depressive disorder and bipolar disorder may increase variant discovery, as well as the discovery of shared and distinct neurobiological gene sets and cell types. Our results also indicate some genetic differences between major depressive disorder and bipolar disorder, including opposite bidirectional relationships of each mood disorder with educational attainment, a possible influence of major depressive disorder on coronary artery disease risk and differing mouse brain cell types implicated by the enrichment patterns of associated genes in each disorder. Finally, our data are consistent with the existence of a genetic mood disorder spectrum with separate clusters for bipolar disorder type 1 and depressive disorders, linked by bipolar disorder type 2, and with depression as the common symptom. The mood disorders have a partially genetic aetiology that is partly shared. The identification of specific sets of genetic variants differentially associated with depression and with mania remains an aim for future research.

## Acknowledgements

This paper has previously been made available as a preprint on bioRxiv at https://www.biorxiv.org/content/10.1101/383331v1.

We are deeply indebted to the investigators who comprise the PGC, and to the hundreds of thousands of subjects who have shared their life experiences with PGC investigators. This study represents independent research partly funded by the National Institute for Health Research (NIHR) Biomedical Research Centre (BRC) at South London and Maudsley NHS Foundation Trust and King’s College London. The views expressed are those of the author(s) and not necessarily those of the NHS, the NIHR or the Department of Health and Social Care. High performance computing facilities were funded with capital equipment grants from the GSTT Charity (TR130505) and Maudsley Charity (980). The PGC has received major funding from the US National Institute of Mental Health (NIMH) and the US National Institute of Drug Abuse (NIDA) of the US National Institutes of Health (NIH; U01 MH109528 to PFS, U01MH109514 to MCO, and U01 MH1095320 to A Agrawal). We acknowledge the continued support of the NL Genetic Cluster Computer (http://www.geneticcluster.org/) hosted by SURFsara in the management and curation of PGC data, with funding from Scientific Organization Netherlands (480-05-003 to DP). Central analysis of PGC data was funded by UK Medical Research Council (MRC) Centre and Program Grants (G0801418, G0800509 to PAH, MCO, MJO) and grants from the Australian National Health and Medical Research Council (NHMRC; 1078901,108788 to NRW). GB, JRIC, HG, CL were supported in part by the NIHR as part of the Maudsley BRC. DP is funded by the Dutch Brain Foundation and the VU University Amsterdam Netherlands. PFS gratefully acknowledges support from the Swedish Research Council (Vetenskapsrådet, award D0886501). Acknowledgements and funding for individual cohorts follows. BD_TRS: This work was funded by the Deutsche Forschungsgemeinschaft (DFG, grants FOR2107 DA1151/5-1, SFB-TRR58, and Project C09 to UD) and the Interdisciplinary Center for Clinical Research (IZKF) of the medical faculty of Münster (grant Dan3/012/17 to UD). BiGS: Research was funded by the NIMH (Chicago: R01 MH103368 to ESG, NIMH: R01 MH061613 and ZIA MH002843 to FJM, Pittsburgh: MH63480 to VN, UCSD: MH078151, MH081804, MH59567 to JK). FJM was supported by the NIMH Intramural Research Program, NIH, DHHS. BOMA-Australia: Funding was supplied by the Australian NHMRC (1037196, 1066177, and 1063960 to JMF, 1103623 to SEM, 1037196 to PBM, 1078399 to GWM, 1037196 to PRS). JMF would like to thank Janette M O’Neil and Betty C Lynch for their support. BOMA-Germany I, BOMA-Germany II, BOMA-Germany III, PsyCourse, and Münster MDD Cohort: This work was supported by the German Ministry for Education and Research (BMBF) through the Integrated Network IntegraMent (Integrated Understanding of Causes and Mechanisms in Mental Disorders), under the auspices of the e:Med program (01ZX1314A/01ZX1614A to MMN and SC, 01ZX1314G/01ZX1614G to MR, 01ZX1314K to TGS) and through grants NGFNplus MooDS (Systematic Investigation of the Molecular Causes of Major Mood Disorders and Schizophrenia; grant 01GS08144, 01GS08147 to MMN, MR and SC). This work was also supported by the DFG (NO246/10-1 to MMN [FOR 2107], RI 908/11-1 to MR [FOR 2107], WI 3429/3-1 to SHW, SCHU 1603/4-1, SCHU 1603/5-1 [KFO 241] and SCHU 1603/7-1 [PsyCourse] to TGS), the Swiss National Science Foundation (SNSF, grant 156791 to SC) and the European Union (N Health-F2-2008-222963 to BTB and VA). MMN is supported through the Excellence Cluster ImmunoSensation. TGS is supported by an unrestricted grant from the Dr. Lisa-Oehler Foundation. AJF received support from the BONFOR Programme of the University of Bonn, Germany. MH was supported by the DFG. BOMA-Romania: The work was supported by Unitatea Executiva Pentru Finantarea Invatamantului Superior a Cercetarii (89/2012 to MG-S). Bulgarian Trios: Recruitment was funded by the Janssen Research Foundation, and genotyping was funded by multiple grants to the Stanley Center for Psychiatric Research at the Broad Institute from the Stanley Medical Research Institute, The Merck Genome Research Foundation, and the Herman Foundation to GK. CoFaMS – Adelaide: Research was funded by the Australian NHMRC (APP1060524 to BTB). CONVERGE: Research was funded by the Wellcome Trust (WT090532/Z/09/Z, WT083573/Z/07/Z and WT089269/Z/09/Z to J Flint) and the NIMH (MH100549 to KSK). Danish RADIANT: Research was funded by Højteknologifonden (0001-2009-2 to TW) and the Lundbeck Foundation, (R24-A3242 to TW). deCODE: Research was funded by FP7-People-2011-IAPP grant agreement PsychDPC, (286213 to KS), and NIDA (R01 DA017932 to KS, R01 DA034076 to TT). The authors are thankful to the participants and staff at the Patient Recruitment Center. Edinburgh: Genotyping was conducted at the Genetics Core Laboratory at the Clinical Research Facility (University of Edinburgh). Research was funded by the Wellcome Trust (104036/Z/14/Z to AMM, T-KC, and DJP). DJM is supported by an NRS Clinical Fellowship funded by the CSO. EGCUT: Research was funded by European Union Project, (EstRC-IUT20-60, No. 2014-2020.4.01.15-0012, 692145 to AM). Fran: This research was supported by Foundation FondaMental, Créteil, France and by the Investissements d’Avenir Programs managed by the ANR (ANR-11-IDEX-0004-02 and ANR-10-COHO-10-01 to ML). GenPOD/Newmeds: Research was funded by MRC (G0200243 to GL and MCO), EU 6th Framework, (LSHB-CT-2003-503428 to RH), IMI-JU, (15008 to GL). GenScot: Research was funded by the UK Chief Scientist Office (CZD/16/6 to DJP) and the Scottish Funding Council (HR03006 to DJP). We are grateful to all the families who took part, the general practitioners and the Scottish School of Primary Care for their help in recruiting them, and the whole Generation Scotland team, which includes interviewers, computer and laboratory technicians, clerical workers, research scientists, volunteers, managers, receptionists, healthcare assistants and nurses. Genotyping was conducted at the Genetics Core Laboratory at the Clinical Research Facility (University of Edinburgh). GERA: Participants in the Genetic Epidemiology Research on Adult Health and Aging Study are part of the Kaiser Permanente Research Program on Genes, Environment, and Health, supported by the NIA, NIMH, OD, (RC2 AG036607 to CS, NRisch) and the Wayne and Gladys Valley Foundation, The Ellison Medical Foundation, the Robert Wood Johnson Foundation, and the Kaiser Permanente Regional and National Community Benefit Programs. GSK_Munich: We thank all participants in the GSK-Munich study. We thank numerous people at GSK and Max-Planck Institute, BKH Augsburg and Klinikum Ingolstadt in Germany who contributed to this project. Halifax: Halifax data were obtained with support from the Canadian Institutes of Health Research to MA. Harvard i2b2: Research funded by NIMH (R01 MH085542 to JWS, R01 MH086026 to RHP). iPSYCH: The iPSYCH (The Lundbeck Foundation Initiative for Integrative Psychiatric Research) team acknowledges funding from The Lundbeck Foundation (grant no R102-A9118 and R155-2014-1724, R129-A3973 and R24-A3243), the Stanley Medical Research Institute, the European Research Council (294838), the Novo Nordisk Foundation for supporting the Danish National Biobank resource, the Capital Region of Denmark, (R144-A5327), and grants from Aarhus and Copenhagen Universities and University Hospitals, including support to the iSEQ Center, the GenomeDK HPC facility, and the CIRRAU Center. All funding was to the iPSYCH PIs: TW, ADB, OM, MN, DH, and PBM. Janssen: Funded by Janssen Research & Development, LLC. We are grateful to the study volunteers for participating in the research studies and to the clinicians and support staff for enabling patient recruitment and blood sample collection. We thank the staff in the former Neuroscience Biomarkers of Janssen Research & Development for laboratory and operational support (e.g., biobanking, processing, plating, and sample de-identification), and to the staff at Illumina for genotyping Janssen DNA samples. MARS/BiDirect: This work was funded by the Max Planck Society, by the Max Planck Excellence Foundation, and by a grant from the German Federal Ministry for Education and Research (BMBF) in the National Genome Research Network framework (NGFN2 and NGFN-Plus, FKZ 01GS0481), and by the BMBF Program FKZ01ES0811. We acknowledge all study participants. We thank numerous people at Max-Planck Institute, and all study sites in Germany and Switzerland who contributed to this project. Controls were from the Dortmund Health Study which was supported by the German Migraine & Headache Society, and by unrestricted grants to the University of Münster from Almirall, Astra Zeneca, Berlin Chemie, Boehringer, Boots Health Care, Glaxo-Smith-Kline, Janssen Cilag, McNeil Pharma, MSD Sharp & Dohme, and Pfizer. Blood collection was funded by the Institute of Epidemiology and Social Medicine, University of Münster. Genotyping was supported by the German Ministry of Research and Education (BMBF grant 01ER0816, 01ER1506 to KB). Mayo Bipolar Disorder Biobank: Research was funded by grants from the Marriot Foundation and the Mayo Clinic Center for Individualized Medicine to JMB and MF. Michigan (NIMH/Pritzker Neuropsychiatric Disorders Research Consortium): Research was funded by NIMH (R01 MH09414501A1, MH105653 to MB). We thank the participants who donated their time and DNA to make this study possible. We thank members of the NIMH Human Genetics Initiative and the University of Michigan Prechter Bipolar DNA Repository for generously providing phenotype data and DNA samples. Many of the authors are members of the Pritzker Neuropsychiatric Disorders Research Consortium which is supported by the Pritzker Neuropsychiatric Disorders Research Fund L.L.C. A shared intellectual property agreement exists between this philanthropic fund and the University of Michigan, Stanford University, the Weill Medical College of Cornell University, HudsonAlpha Institute of Biotechnology, the Universities of California at Davis, and at Irvine, to encourage the development of appropriate findings for research and clinical applications. Mount Sinai: This work was funded in part by a NARSAD Young Investigator award to EAS, and by NIH (R01MH106531, R01MH109536 to PS and EAS). NeuRA-CASSI-Australia: This work was funded by the NSW Ministry of Health, Office of Health and Medical Research, and by the NHRMC (568807 to CSW and TWW). CSW was a recipient of NHMRC Fellowships (#1117079, #1021970). NeuRA-IGP-Australia: Research was funded by the NHMRC (630471, 1061875, 1081603 to MJG. NESDA: Research was funded by Nederlandse Organisatie voor Wetenschappelijk (NOW; ZonMW Geestkracht grant to PWJHP). Norway: Research was funded by the Vetenskapsrådet to IA, the Western Norway Regional Health Authority to KJO, the Research Council of Norway (#421716 to IM, #249711, #248778, #223273, and #217776 to OAA), the South-East Norway Regional Health Authority (#2012-132 and #2012-131 to OAA, #2016-064 to OBS, #2017-004 to OAA and OBS, #2013-088, #2014-102, and #2011-085 to IM), and the KG Jebsen Stiftelsen to OAA. TE was funded by The South-East Norway Regional Health Authority (#2015-078) and a research grant from Mrs. Throne-Holst. NTR: Research was funded by NWO (480-15-001/674 to DIB). Pfizer: Research was funded by the EU Innovative Medicine Initiative Joint Undertaking (115008.5). PsyColaus: PsyCoLaus/CoLaus received additional support from research grants from GlaxoSmithKline and the Faculty of Biology and Medicine of Lausanne, and the SNSF (3200B0–105993, 3200B0-118308, 33CSCO-122661, 33CS30-139468, 33CS30-148401 to MP). QIMR: We thank the twins and their families for their willing participation in our studies. Research was funded by NHMRC (941177, 971232, 3399450 and 443011 to NGM) and NIAAA (AA07535, AA07728, and AA10249 to ACH). RADIANT: Research was funded by MRC (G0701420 to GB and CML, G0901245 to GB) and NIMH (U01 MH109528 to GB). Rotterdam Study: The Rotterdam Study is funded by Erasmus Medical Center and Erasmus University, and NWO (175.010.2005.011, 911-03-012 to AGU). SHIP-LEGEND/TREND: SHIP is part of the Community Medicine Research net of the University of Greifswald which is funded by the DFG (GR 1912/5-1 to HJG), Federal Ministry of Education and Research (grants 01ZZ9603, 01ZZ0103, and 01ZZ0403), the Ministry of Cultural Affairs, and the Social Ministry of the Federal State of Mecklenburg-West Pomerania. Genotyping in SHIP was funded by Siemens Healthineers and the Federal State of Mecklenburg-West Pomerania. Genotyping in SHIP-TREND-0 was supported by the Federal Ministry of Education and Research (grant 03ZIK012). Span2: Research was funded by Instituto de Salud Carlos III (PI12/01139, PI14/01700, PI15/01789, PI16/01505), and cofinanced by the European Regional Development Fund (ERDF), Agència de Gestió d’Ajuts Universitaris i de Recerca-AGAUR, Generalitat de Catalunya (2014SGR1357), Departament de Salut, Generalitat de Catalunya, Spain, and a NARSAD Young Investigator Grant from the Brain & Behavior Research Foundation. This project has also received funding from the European Union’s Horizon 2020 Research and Innovation Programme under the grant agreements No 667302 and 643051. CSM is a recipient of a Sara Borrell contract (CD15/00199) and a mobility grant (MV16/00039) from the Instituto de Salud Carlos III, Ministerio de Economía, Industria y Competitividad, Spain. MR is a recipient of a Miguel de Servet contract (CP09/00119 and CPII15/00023) from the Instituto de Salud Carlos III, Ministerio de Economía, Industria y Competitividad, Spain. STAR*D: Research was funded by NIMH (R01 MH-072802 to SPH). The authors appreciate the efforts of the STAR*D investigator team for acquiring, compiling, and sharing the STAR*D clinical data set. SUNY DMC: Research was funded by NIMH (R01MH085542 to CP, MTP, JAK, and HM). SWEBIC: Research was funded by NIMH (MH077139), the Vetenskapsrådet (K2014-62X-14647-12-51 and K2010-61P-21568-01-4), the Swedish foundation for Strategic Research (KF10-0039) and the Stanley Center for Psychiatric Research, Broad Institute from a grant from Stanley Medical Research Institute, all to ML. We are deeply grateful for the participation of all subjects contributing to this research, and to the collection team that worked to recruit them. We also wish to thank the Swedish National Quality Register for Bipolar Disorders: BipoläR. Sweden: This work was funded by the Vetenskapsrådet (to MS and CL), the Stockholm County Council (to MS, CL, LB, LF, and UÖ) and the Söderström Foundation (to LB). TwinGene: Research was funded by GenomeEUtwin, (EU/QLRT-2001-01254; QLG2-CT-2002-01254 to NLP), Heart and Lung Foundation (20070481 to PKM), SFF and Vetenskapsrådet, (M-2005-1112 to U de Faire).We thank the Karolinska Institutet for infrastructural support of the Swedish Twin Registry. UCL: Research was funded by the MRC (G1000708 to AM). UCLA-Utrecht (Los Angeles): Research was funded by NIMH (R01MH090553, U01MH105578 to NBF, RAO, LMOL, and APSO). UK -BDRN: Research was funded by MRC Centre and Program Grants (G0801418, G0800509 to MCO and MJO), the Wellcome Trust (078901 to NC, IJ, LAJ), the Stanley Medical Research Institute (5710002223-01 to NC, IJ, LAJ), and a European Commission Marie Curie Fellowship (623932 to ADF). BDRN would like to acknowledge the research participants who continue to give their time to participate in our research. UK Biobank: This research has been carried out under application numbers 4844, 6818, and 16577, funded by the National Institute for Health Research under its Biomedical Research Centres funding initiative (to GB) and the Wellcome Trust (04036/Z/14/Z to AMM). UNIBO / University of Barcelona, Hospital Clinic, IDIBAPS, CIBERSAM: EV thanks the support of the Spanish Ministry of Economy and Competitiveness (PI15/00283 to EV) integrated into the Plan Nacional de I+D+I y cofinanciado por el ISCIII-Subdirección General de Evaluación y el Fondo Europeo de Desarrollo Regional (FEDER); CIBERSAM; and the Comissionat per a Universitats i Recerca del DIUE de la Generalitat de Catalunya to the Bipolar Disorders Group (2014 SGR 398). USC: Research funded by NIH (R01MH085542 to JLS). WTCCC: The principal funder of this project was the Wellcome Trust to NC and AHY. For the 1958 Birth Cohort, venous blood collection was funded by the UK MRC. AHY is funded by the National Institute for Health Research (NIHR) Biomedical Research Centre at South London and Maudsley NHS Foundation Trust and King’s College London. 23andMe: We thank the 23andMe research participants included in the analysis, all of whom provided informed consent and participated in the research online according to a human subjects protocol approved by an external AAHRPP-accredited institutional review board (Ethical & Independent Review Services), and the employees of 23andMe for making this work possible. 23andMe acknowledges the-invaluable contributions of Michelle Agee, Babak Alipanahi, Adam Auton, Robert K. Bell, Katarzyna Bryc, Sarah L. Elson, Pierre Fontanillas, Nicholas A. Furlotte, David A. Hinds, Bethann S. Hromatka, Karen E. Huber, Aaron Kleinman, Nadia K. Litterman, Matthew H. McIntyre, Joanna L. Mountain, Carrie A.M. Northover, Steven J. Pitts, J. Fah Sathirapongsasuti, Olga V. Sazonova, Janie F. Shelton, Suyash Shringarpure, Chao Tian, Joyce Y. Tung, Vladimir Vacic, and Catherine H. Wilson.

## Article Information

The Bipolar Disorder and Major Depressive Disorder Working Groups of the Psychiatric Genomics Consortium are collaborative co-authors for this article. The individual authors are (numbers refer to affiliations listed in the Supplement): Enda M. Byrne^4^, Andreas J. Forstner^5;6;7;8;9^, Peter A. Holmans^10^, Christiaan A. de Leeuw^11^, Manuel Mattheisen^12;13;14;15;16^, Andrew McQuillin^17^, Jennifer M. Whitehead Pavlides^18^, Tune H. Pers^19;20^, Stephan Ripke^21;22;23^, Eli A. Stahl^19;24;25^, Stacy Steinberg^26^, Vassily Trubetskoy^22^, Maciej Trzaskowski^4^, Yunpeng Wang^27;28^, Liam Abbott^21^, Abdel Abdellaoui^29^, Mark J. Adams^30^, Annelie Nordin Adolfsson^31^, Esben Agerbo^16;32;33^, Huda Akil^34^, Diego Albani^35^, Ney Alliey-Rodriguez^36^, Thomas D. Als^12;13;16^, Till F. M. Andlauer^37;38^, Adebayo Anjorin^39^, Verneri Antilla^23^, Sandra Van der Auwera^40^, Swapnil Awasthi^22^, Silviu-Alin Bacanu^41^, Judith A Badner^42^, Marie Bækvad-Hansen^16;43^, Jack D. Barchas^44^, Nicholas Bass^17^, Michael Bauer^45^, Aartjan T. F. Beekman^46^, Richard Belliveau^21^, Sarah E. Bergen^3^, Tim B. Bigdeli^41;47^, Elisabeth B. Binder^37;48^, Erlend Bøen^49^, Marco Boks^50^, James Boocock^51^, Monika Budde^52^, William Bunney^53^, Margit Burmeister^54^, Henriette N. Buttenschøn^3;12;55^, Jonas Bybjerg-Grauholm^16;43^, William Byerley^56^, Na Cai^57;58^, Miquel Casas^59;60;61;62^, Enrique Castelao^63^, Felecia Cerrato^21^, Pablo Cervantes^64^, Kimberly Chambert^21^, Alexander W. Charney^25^, Danfeng Chen^21^, Jane Hvarregaard Christensen^12;13;55^, Claire Churchhouse^21;23^, David St Clair^65^, Toni-Kim Clarke^30^, Lucía Colodro-Conde^66^, William Coryell^67^, Baptiste Couvy-Duchesne^18;68^, David W. Craig^69^, Gregory E. Crawford^70;71^, Cristiana Cruceanu^37;64^, Piotr M. Czerski^72^, Anders M. Dale^73;74;75;76^, Gail Davies^77^, Ian J. Deary^77^, Franziska Degenhardt^7;8^, Jurgen Del-Favero^78^, J Raymond DePaulo^79^, Eske M. Derks^66^, Nese Direk^80;81^, Srdjan Djurovic^82;83^, Amanda L. Dobbyn^24;25^, Conor V. Dolan^29^, Ashley Dumont^21^, Erin C. Dunn^21;84;85^, Thalia C. Eley^1^, Torbjørn Elvsåshagen^86;87^, Valentina Escott-Price^10^, Chun Chieh Fan^76^, Hilary K. Finucane^88;89^, Sascha B. Fischer^5;9^, Matthew Flickinger^90^, Jerome C. Foo^91^, Tatiana M. Foroud^92^, Liz Forty^10^, Josef Frank^91^, Christine Fraser^10^, Nelson B. Freimer^93^, Louise Frisén^94;95;96^, Katrin Gade^52;97^, Diane Gage^21^, Julie Garnham^98^, Claudia Giambartolomei^51^, Fernando S. Goes^99^, Jaqueline Goldstein^21^, Scott D. Gordon^66^, Katherine Gordon-Smith^100^, Elaine K. Green^101^, Melissa J. Green^102^, Tiffany A. Greenwood^75^, Jakob Grove^12;13;16;103^, Weihua Guan^104^, Lynsey S. Hall^30;105^, Marian L. Hamshere^10^, Christine Søholm Hansen^16;43^, Thomas F. Hansen^16;106;107^, Martin Hautzinger^108^, Urs Heilbronner^52^, Albert M. van Hemert^109^, Stefan Herms^5;7;8;9^, Ian B. Hickie^110^, Maria Hipolito^111^, Per Hoffmann^5;7;8;9^, Dominic Holland^73;112^, Georg Homuth^113^, Carsten Horn^114^, Jouke-Jan Hottenga^29^, Laura Huckins^24;25^, Marcus Ising^15^, Stéphane Jamain^116;117^, Rick Jansen^46^, Jessica S. Johnson^24;25^, Simone de Jong^1;2^, Eric Jorgenson^118^, Anders Juréus^3^, Radhika Kandaswamy^1^, Robert Karlsson^3^, James L. Kennedy^119;120;121;122^, Farnush Farhadi Hassan Kiadeh^123^, Sarah Kittel-Schneider^124^, James A. Knowles^125;126^, Manolis Kogevinas^127^, Isaac S. Kohane^128;129;130^, Anna C. Koller^7;8^, Julia Kraft^22^, Warren W. Kretzschmar^131^, Jesper Krogh^132^, Ralph Kupka^46;133^, Zoltán Kutalik^134;135^, Catharina Lavebratt^94^, Jacob Lawrence^136^, William B. Lawson^111^, Markus Leber^137^, Phil H. Lee^21;23;138^, Shawn E. Levy^139^, Jun Z. Li^140^, Yihan Li^131^, Penelope A. Lind^66^, Chunyu Liu^141^, Loes M. Olde Loohuis^93^, Anna Maaser^7;8^, Donald J. MacIntyre^142;143^, Dean F. MacKinnon^99^, Pamela B. Mahon^79;144^, Wolfgang Maier^145^, Robert M. Maier^18^, Jonathan Marchini^146^, Lina Martinsson^95^, Hamdi Mbarek^29^, Steve McCarroll^21;147^, Patrick McGrath^148^, Peter McGuffin^1^, Melvin G. McInnis^149^, James D. McKay^150^, Helena Medeiros^126^, Sarah E. Medland^66^, Divya Mehta^18;151^, Fan Meng^34;149^, Christel M. Middeldorp^29;152;153^, Evelin Mihailov^154^, Yuri Milaneschi^46^, Lili Milani^154^, Saira Saeed Mirza^80^, Francis M. Mondimore^99^, Grant W. Montgomery^4^, Derek W. Morris^155;156^, Sara Mostafavi^157;158^, Thomas W Mühleisen^5;159^, Niamh Mullins^1^, Matthias Nauck^160;161^, Bernard Ng^158^, Hoang Nguyen^24;25^, Caroline M. Nievergelt^75;162^, Michel G. Nivard^29^, Evaristus A. Nwulia^111^, Dale R. Nyholt^163^, Claire O’Donovan^98^, Paul F. O’Reilly^1^, Anil P. S. Ori^93^, Lilijana Oruc^164^, Urban Ösby^165^, Hogni Oskarsson^166^, Jodie N. Painter^66^, José Guzman Parra^167^, Carsten Bøcker Pedersen^16;32;33^, Marianne Giørtz Pedersen^16;32;33^, Amy Perry^100^, Roseann E. Peterson^41;168^, Erik Pettersson^3^, Wouter J. Peyrot^46^, Andrea Pfennig^45^, Giorgio Pistis^63^, Shaun M. Purcell^25;144^, Jorge A. Quiroz^169^, Per Qvist^12;13;55^, Eline J. Regeer^170^, Andreas Reif^124^, Céline S. Reinbold^5;9^, John P. Rice^171^, Brien P. Riley^41^, Fabio Rivas^167^, Margarita Rivera^1;172^, Panos Roussos^24;25;173^, Douglas M. Ruderfer^174^, Euijung Ryu^175^, Cristina Sánchez-Mora^59;60;62^, Alan F. Schatzberg^176^, William A. Scheftner^177^, Robert Schoevers^178^, Nicholas J. Schork^179^, Eva C. Schulte^52;180^, Tatyana Shehktman^75^, Ling Shen^118^, Jianxin Shi^181^, Paul D. Shilling^75^, Stanley I. Shyn^182^, Engilbert Sigurdsson^183^, Claire Slaney^98^, Olav B. Smeland^73;184;185^, Johannes H. Smit^46^, Daniel J. Smith^186^, Janet L. Sobell^187^, Anne T. Spijker^188^, Michael Steffens^189^, John S. Strauss^121;190^, Fabian Streit^91^, Jana Strohmaier^91^, Szabolcs Szelinger^191^, Katherine E. Tansey^192^, Henning Teismann^193^, Alexander Teumer^194^, Robert C Thompson^149^, Wesley Thompson^55;75;87;107^, Pippa A. Thomson^195^, Thorgeir E. Thorgeirsson^26^, Matthew Traylor^196^, Jens Treutlein^91^, André G. Uitterlinden^197^, Daniel Umbricht^198^, Helmut Vedder^199^, Alexander Viktorin^3^, Peter M. Visscher^4;18^, Weiqing Wang^24;25^, Stanley J. Watson^149^, Bradley T. Webb^168^, Cynthia Shannon Weickert^102;200^, Thomas W. Weickert^102;200^, Shantel Marie Weinsheimer^55;107^, Jürgen Wellmann^193^, Gonneke Willemsen^29^, Stephanie H. Witt^91^, Yang Wu^4^, Hualin S. Xi^201^, Wei Xu^202;203^, Jian Yang^4;18^, Allan H. Young^204^, Peter Zandi^205^, Peng Zhang^206^, Futao Zhang^4^, Sebastian Zollner^149^, Rolf Adolfsson^31^, Ingrid Agartz^14;49;207^, Martin Alda^98;208^, Volker Arolt^209^, Lena Backlund^95^, Bernhard T. Baune^210^, Frank Bellivier^211;212;213;214^, Klaus Berger^193^, Wade H. Berrettini^215^, Joanna M. Biernacka^175^, Douglas H. R. Blackwood^30^, Michael Boehnke^90^, Dorret I. Boomsma^29^, Aiden Corvin^156^, Nicholas Craddock^10^, Mark J. Daly^21;23^, Udo Dannlowski^209^, Enrico Domenici^216^, Katharina Domschke^217^, Tõnu Esko^19;147;154;218^, Bruno Etain^211;213;214;219^, Mark Frye^220^, Janice M. Fullerton^200;221^, Elliot S. Gershon^36;222^, EJC de Geus^29;223^, Michael Gill^156^, Fernando Goes^79^, Hans J. Grabe^40^, Maria Grigoroiu-Serbanescu^224^, Steven P. Hamilton^225^, Joanna Hauser^72^, Caroline Hayward^226^, Andrew C. Heath^171^, David M. Hougaard^16;43^, Christina M. Hultman^3^, Ian Jones^10^, Lisa A. Jones^100^, René S. Kahn^25;50^, Kenneth S. Kendler^41^, George Kirov^10^, Stefan Kloiber^115;121;190^, Mikael Landén^3;227^, Marion Leboyer^117;211;228^, Glyn Lewis^17^, Qingqin S. Li^229^, Jolanta Lissowska^230^, Susanne Lucae^115^, Pamela A. F. Madden^119^, Patrik K. Magnusson^3^, Nicholas G. Martin^66;231^, Fermin Mayoral^167^, Susan L. McElroy^232^, Andrew M. McIntosh^30;77^, Francis J. McMahon^233^, Ingrid Melle^234;235^, Andres Metspalu^154;236^, Philip B. Mitchell^102^, Gunnar Morken^237;238^, Ole Mors^16;239^, Preben Bo Mortensen^12;16;32;33^, Bertram Müller-Myhsok^37;240;241^, Richard M. Myers^139^, Benjamin M. Neale^19;21;23^, Vishwajit Nimgaonkar^242^, Merete Nordentoft^16;243^, Markus M. Nöthen^7;8^, Michael C. O’Donovan^10^, Ketil J. Oedegaard^244;245^, Michael J. Owen^10^, Sara A. Paciga^246^, Carlos Pato^126;247^, Michele T. Pato^126^, Nancy L. Pedersen^3^, Brenda W. J. H. Penninx^46^, Roy H. Perlis^248;249^, David J. Porteous^195^, Danielle Posthuma^11;250^, James B. Potash^79^, Martin Preisig^63^, Josep Antoni Ramos-Quiroga^59;60;61;62^, Marta Ribasés^59;60;62^, Marcella Rietschel^91^, Guy A. Rouleau^251;252^, Catherine Schaefer^118^, Martin Schalling^94^, Peter R. Schofield^200;221^, Thomas G. Schulze^52;79;91;97;233^, Alessandro Serretti^253^, Jordan W. Smoller^21;84;85^, Hreinn Stefansson^26^, Kari Stefansson^26;254^, Eystein Stordal^255;256^, Henning Tiemeier^80;257;258^, Gustavo Turecki^259^, Rudolf Uher^98^, Arne E. Vaaler^260^, Eduard Vieta^261^, John B. Vincent^190^, Henry Völzke^194^, Myrna M. Weissman^148;262^, Thomas Werge^16;107;263^, Ole A. Andreassen^184;185^, Anders D. Børglum^12;13;16^, Sven Cichon^5;7;9;159^, Howard J. Edenberg^264^, Arianna Di Florio^10;265^, John Kelsoe^75^, Douglas F. Levinson^176^, Cathryn M. Lewis^1;2;266^, John I. Nurnberger^92;267^, Roel A. Ophoff^50;51;93^, Laura J. Scott^90^, Pamela Sklar^24;25†^, Patrick F. Sullivan^3;265;268^, Naomi R. Wray^4;18^.

## Data availability

GWAS results from analyses including 23andMe are restricted by a data transfer agreement with 23andMe. For these analyses, LD-independent sets of 10,000 SNPs will be made available via the Psychiatric Genetics Consortium (https://www.med.unc.edu/pgc/results-and-downloads). Summary statistics not including 23andMe will be made available via the Psychiatric Genetics Consortium (https://www.med.unc.edu/pgc/results-and-downloads).

