## Supplementary Information for "The genetics of the mood disorder spectrum: genome-wide association analyses of over 185,000 cases and 439,000 controls"

### Consortium Affiliations

1	Social, Genetic and Developmental Psychiatry Centre, Institute of Psychiatry, Psychology and Neuroscience, King's College London, London, United Kingdom

2	NIHR Maudsley Biomedical Research Centre, King's College London, London, United Kingdom

3	Department of Medical Epidemiology and Biostatistics, Karolinska Institutet, Stockholm, Sweden

4	Institute for Molecular Bioscience, The University of Queensland, Brisbane, QLD, Australia

5	Human Genomics Research Group, Department of Biomedicine, University of Basel, Basel, Switzerland

6	Department of Psychiatry (UPK), University of Basel, Basel, Switzerland

7	Institute of Human Genetics, University of Bonn, School of Medicine & University Hospital Bonn, Bonn, Germany

8	Department of Genomics, Life&Brain Center, University of Bonn, Bonn, Germany

9	Institute of Medical Genetics and Pathology, University Hospital Basel, Basel, Switzerland

10	Medical Research Council Centre for Neuropsychiatric Genetics and Genomics, Division of Psychological Medicine and Clinical Neurosciences, Cardiff University, Cardiff, United Kingdom

11	Department of Complex Trait Genetics, Center for Neurogenomics and Cognitive Research, Amsterdam Neuroscience, Vrije Universiteit Amsterdam, Amsterdam, Netherlands

12	iSEQ, Center for Integrative Sequencing, Aarhus University, Aarhus, Denmark

13	Department of Biomedicine - Human Genetics, Aarhus University, Aarhus, Denmark

14	Department of Clinical Neuroscience, Centre for Psychiatry Research, Karolinska Institutet, Stockholm, Sweden

15	Department of Psychiatry, Psychosomatics and Psychotherapy, Center of Mental Health, University Hospital Würzburg, Würzburg, Germany

16	iPSYCH, The Lundbeck Foundation Initiative for Psychiatric Research, Copenhagen, Denmark

17	Division of Psychiatry, University College London, London, United Kingdom

18	Queensland Brain Institute, The University of Queensland, Brisbane, QLD, Australia

19	Medical and Population Genetics, Broad Institute, Cambridge, MA, USA

20	Division of Endocrinology and Center for Basic and Translational Obesity Research, Boston Children’s Hospital, Boston, MA, USA

21	Stanley Center for Psychiatric Research, Broad Institute, Cambridge, MA, USA

22	Department of Psychiatry and Psychotherapy, Charité - Universitätsmedizin, Berlin, Germany

23	Analytic and Translational Genetics Unit, Massachusetts General Hospital, Boston, MA, USA

24	Department of Genetics and Genomic Sciences, Icahn School of Medicine at Mount Sinai, New York, NY, USA

25	Department of Psychiatry, Icahn School of Medicine at Mount Sinai, New York, NY, USA

26	deCODE Genetics / Amgen, Reykjavik, Iceland

27	Institute of Biological Psychiatry, Mental Health Centre Sct. Hans, Copenhagen, Denmark

28	Institute of Clinical Medicine, University of Oslo, Oslo, Norway

29	Dept of Biological Psychology & EMGO+ Institute for Health and Care Research, Vrije Universiteit Amsterdam, Amsterdam, Netherlands

30	Division of Psychiatry, University of Edinburgh, Edinburgh, United Kingdom

31	Department of Clinical Sciences, Psychiatry, Umeå University Medical Faculty, Umeå, Sweden

32	National Centre for Register-Based Research, Aarhus University, Aarhus, Denmark

33	Centre for Integrated Register-based Research, Aarhus University, Aarhus, Denmark

34	Molecular & Behavioral Neuroscience Institute, University of Michigan, Ann Arbor, MI, USA

35	Neuroscience, Istituto Di Ricerche Farmacologiche Mario Negri, Milano, Italy

36	Department of Psychiatry and Behavioral Neuroscience, University of Chicago, Chicago, IL, USA

37	Department of Translational Research in Psychiatry, Max Planck Institute of Psychiatry, Munich, Germany

38	Department of Neurology, Klinikum rechts der Isar, Technical University of Munich, Munich, Germany

39	Psychiatry, Berkshire Healthcare NHS Foundation Trust, Bracknell, United Kingdom

40	Department of Psychiatry and Psychotherapy, University Medicine Greifswald, Greifswald, Mecklenburg-Vorpommern, Germany

41	Department of Psychiatry, Virginia Commonwealth University, Richmond, VA, USA

42	Psychiatry, Rush University Medical Center, Chicago, IL, USA

43	Center for Neonatal Screening, Department for Congenital Disorders, Statens Serum Institut, Copenhagen, Denmark

44	Department of Psychiatry, Weill Cornell Medical College, New York, NY, USA

45	Department of Psychiatry and Psychotherapy, University Hospital Carl Gustav Carus, Technische Universität Dresden, Dresden, Germany

46	Department of Psychiatry, Vrije Universiteit Medical Center and GGZ inGeest, Amsterdam, Netherlands

47	Virginia Institute for Psychiatric and Behavior Genetics, Richmond, VA, USA

48	Department of Psychiatry and Behavioral Sciences, Emory University School of Medicine, Atlanta, GA, USA

49	Department of Psychiatric Research, Diakonhjemmet Hospital, Oslo, Norway

50	Psychiatry, UMC Utrecht Hersencentrum Rudolf Magnus, Utrecht, Netherlands

51	Human Genetics, University of California Los Angeles, Los Angeles, CA, USA

52	Institute of Psychiatric Phenomics and Genomics (IPPG), University Hospital, LMU Munich, Munich, Germany

53	Department of Psychiatry and Human Behavior, University of California, Irvine, Irvine, CA, USA

54	Molecular & Behavioral Neuroscience Institute and Department of Computational Medicine & Bioinformatics, University of Michigan, Ann Arbor, MI, USA

55	iPSYCH, The Lundbeck Foundation Initiative for Integrative Psychiatric Research, Denmark

56	Psychiatry, University of California San Francisco, San Francisco, CA, USA

57	Human Genetics, Wellcome Trust Sanger Institute, Cambridge, United Kingdom

58	Statistical genomics and systems genetics, European Bioinformatics Institute (EMBL-EBI), Cambridge, United Kingdom

59	Instituto de Salud Carlos III, Biomedical Network Research Centre on Mental Health (CIBERSAM), Madrid, Spain

60	Department of Psychiatry, Hospital Universitari Vall d´Hebron, Barcelona, Spain

61	Department of Psychiatry and Forensic Medicine, Universitat Autònoma de Barcelona, Barcelona, Spain

62	Psychiatric Genetics Unit, Group of Psychiatry Mental Health and Addictions, Vall d´Hebron Research Institut (VHIR), Universitat Autònoma de Barcelona, Barcelona, Spain

63	Department of Psychiatry, University Hospital of Lausanne, Prilly, Vaud, Switzerland

64	Department of Psychiatry, Mood Disorders Program, McGill University Health Center, Montreal, QC, Canada

65	Institute for Medical Sciences, University of Aberdeen, Aberdeen, United Kingdom

66	Genetics and Computational Biology, QIMR Berghofer Medical Research Institute, Brisbane, QLD, Australia

67	University of Iowa Hospitals and Clinics, Iowa City, IA, USA

68	Centre for Advanced Imaging, The University of Queensland, Brisbane, QLD, Australia

69	Translational Genomics, USC, Phoenix, AZ, USA

70	Center for Genomic and Computational Biology, Duke University, Durham, NC, USA

71	Department of Pediatrics, Division of Medical Genetics, Duke University, Durham, NC, USA

72	Department of Psychiatry, Laboratory of Psychiatric Genetics, Poznan University of Medical Sciences, Poznan, Poland

73	Department of Neurosciences, University of California San Diego, La Jolla, CA, USA

74	Department of Radiology, University of California San Diego, La Jolla, CA, USA

75	Department of Psychiatry, University of California San Diego, La Jolla, CA, USA

76	Department of Cognitive Science, University of California San Diego, La Jolla, CA, USA

77	Centre for Cognitive Ageing and Cognitive Epidemiology, University of Edinburgh, Edinburgh, United Kingdom

78	Applied Molecular Genomics Unit, VIB Department of Molecular Genetics, University of Antwerp, Antwerp, Belgium

79	Department of Psychiatry and Behavioral Sciences, Johns Hopkins University, Baltimore, MD, USA

80	Epidemiology, Erasmus MC, Rotterdam, Zuid-Holland, Netherlands

81	Psychiatry, Dokuz Eylul University School Of Medicine, Izmir, Turkey

82	Department of Medical Genetics, Oslo University Hospital Ullevål, Oslo, Norway

83	NORMENT, KG Jebsen Centre for Psychosis Research, Department of Clinical Science, University of Bergen, Bergen, Norway

84	Department of Psychiatry, Massachusetts General Hospital, Boston, MA, USA

85	Psychiatric and Neurodevelopmental Genetics Unit (PNGU), Massachusetts General Hospital, Boston, MA, USA

86	Department of Neurology, Oslo University Hospital, Oslo, Norway

87	NORMENT, KG Jebsen Centre for Psychosis Research, Oslo University Hospital, Oslo, Norway

88	Department of Epidemiology, Harvard T.H. Chan School of Public Health, Boston, MA, USA

89	Department of Mathematics, Massachusetts Institute of Technology, Cambridge, MA, USA

90	Center for Statistical Genetics and Department of Biostatistics, University of Michigan, Ann Arbor, MI, USA

91	Department of Genetic Epidemiology in Psychiatry, Central Institute of Mental Health, Medical Faculty Mannheim, Heidelberg University, Mannheim, Germany

92	Department of Medical & Molecular Genetics, Indiana University, Indianapolis, IN, USA

93	Center for Neurobehavioral Genetics, University of California Los Angeles, Los Angeles, CA, USA

94	Department of Molecular Medicine and Surgery, Karolinska Institutet and Center for Molecular Medicine, Karolinska University Hospital, Stockholm, Sweden

95	Department of Clinical Neuroscience, Karolinska Institutet and Center for Molecular Medicine, Karolinska University Hospital, Stockholm, Sweden

96	Child and Adolescent Psychiatry Research Center, Stockholm, Sweden

97	Department of Psychiatry and Psychotherapy, University Medical Center Göttingen, Göttingen, Germany

98	Department of Psychiatry, Dalhousie University, Halifax, NS, Canada

99	Psychiatry & Behavioral Sciences, Johns Hopkins University, Baltimore, MD, USA

100	Department of Psychological Medicine, University of Worcester, Worcester, United Kingdom

101	School of Biomedical and Healthcare Sciences, Plymouth University Peninsula Schools of Medicine and Dentistry, Plymouth, United Kingdom

102	School of Psychiatry, University of New South Wales, Sydney, NSW, Australia

103	Bioinformatics Research Centre, Aarhus University, Aarhus, Denmark

104	Biostatistics, University of Minnesota System, Minneapolis, MN, USA

105	Institute of Genetic Medicine, Newcastle University, Newcastle upon Tyne, United Kingdom

106	Danish Headache Centre, Department of Neurology, Rigshospitalet, Glostrup, Denmark

107	Institute of Biological Psychiatry, MHC Sct. Hans, Mental Health Services Copenhagen, Roskilde, Denmark

108	Department of Psychology, Eberhard Karls Universität Tübingen, Tubingen, Germany

109	Department of Psychiatry, Leiden University Medical Center, Leiden, Netherlands

110	Brain and Mind Centre, University of Sydney, Sydney, NSW, Australia

111	Department of Psychiatry and Behavioral Sciences, Howard University Hospital, Washington, DC, USA

112	Center for Multimodal Imaging and Genetics, University of California San Diego, La Jolla, CA, USA

113	Interfaculty Institute for Genetics and Functional Genomics, Department of Functional Genomics, University Medicine and Ernst Moritz Arndt University Greifswald, Greifswald, Mecklenburg-Vorpommern, Germany

114	Roche Pharmaceutical Research and Early Development, Pharmaceutical Sciences, Roche Innovation Center Basel, F. Hoffmann-La Roche Ltd, Basel, Switzerland

115	Max Planck Institute of Psychiatry, Munich, Germany

116	Psychiatrie Translationnelle, Inserm U955, Créteil, France

117	Faculté de Médecine, Université Paris Est, Créteil, France

118	Division of Research, Kaiser Permanente Northern California, Oakland, CA, USA

119	Campbell Family Mental Health Research Institute, Centre for Addiction and Mental Health, Toronto, ON, Canada

120	Neurogenetics Section, Centre for Addiction and Mental Health, Toronto, ON, Canada

121	Department of Psychiatry, University of Toronto, Toronto, ON, Canada

122	Institute of Medical Sciences, University of Toronto, Toronto, ON, Canada

123	Bioinformatics, University of British Columbia, Vancouver, BC, Canada

124	Department of Psychiatry, Psychosomatic Medicine and Psychotherapy, University Hospital Frankfurt, Frankfurt am Main, Germany

125	Cell Biology, SUNY Downstate Medical Center College of Medicine, Brooklyn, NY, USA

126	Institute for Genomic Health, SUNY Downstate Medical Center College of Medicine, Brooklyn, NY, USA

127	Center for Research in Environmental Epidemiology (CREAL), Barcelona, Spain

128	Department of Biomedical Informatics, Harvard Medical School, Boston, MA, USA

129	Department of Medicine, Brigham and Women's Hospital, Boston, MA, USA

130	Informatics Program, Boston Children's Hospital, Boston, MA, USA

131	Wellcome Trust Centre for Human Genetics, University of Oxford, Oxford, United Kingdom

132	Department of Endocrinology at Herlev University Hospital, University of Copenhagen, Copenhagen, Denmark

133	Psychiatry, Altrecht, Utrecht, Netherlands

134	Institute of Social and Preventive Medicine (IUMSP), University Hospital of Lausanne, Lausanne, VD, Switzerland

135	Swiss Institute of Bioinformatics, Lausanne, VD, Switzerland

136	Psychiatry, North East London NHS Foundation Trust, Ilford, United Kingdom

137	Clinic for Psychiatry and Psychotherapy, University Hospital Cologne, Cologne, Germany

138	Psychiatric and Neurodevelopmental Genetics Unit, Massachusetts General Hospital, Boston, MA, USA

139	HudsonAlpha Institute for Biotechnology, Huntsville, AL, USA

140	Department of Human Genetics, University of Michigan, Ann Arbor, MI, USA

141	Psychiatry, University of Illinois at Chicago College of Medicine, Chicago, IL, USA

142	Mental Health, NHS 24, Glasgow, United Kingdom

143	Division of Psychiatry, Centre for Clinical Brain Sciences, University of Edinburgh, Edinburgh, United Kingdom

144	Psychiatry, Brigham and Women's Hospital, Boston, MA, USA

145	Department of Psychiatry and Psychotherapy, University of Bonn, Bonn, Germany

146	Statistics, University of Oxford, Oxford, United Kingdom

147	Department of Genetics, Harvard Medical School, Boston, MA, USA

148	Psychiatry, Columbia University College of Physicians and Surgeons, New York, NY, USA

149	Department of Psychiatry, University of Michigan, Ann Arbor, MI, USA

150	Genetic Cancer Susceptibility Group, International Agency for Research on Cancer, Lyon, France

151	School of Psychology and Counseling, Queensland University of Technology, Brisbane, QLD, Australia

152	Child and Youth Mental Health Service, Children's Health Queensland Hospital and Health Service, South Brisbane, QLD, Australia

153	Child Health Research Centre, University of Queensland, Brisbane, QLD, Australia

154	Estonian Genome Center, University of Tartu, Tartu, Estonia

155	Discipline of Biochemistry, Neuroimaging and Cognitive Genomics (NICOG) Centre, National University of Ireland, Galway, Galway, Ireland

156	Neuropsychiatric Genetics Research Group, Dept of Psychiatry and Trinity Translational Medicine Institute, Trinity College Dublin, Dublin, Ireland

157	Medical Genetics, University of British Columbia, Vancouver, BC, Canada

158	Statistics, University of British Columbia, Vancouver, BC, Canada

159	Institute of Neuroscience and Medicine (INM-1), Research Centre Jülich, Jülich, Germany

160	DZHK (German Centre for Cardiovascular Research), Partner Site Greifswald, University Medicine, University Medicine Greifswald, Greifswald, Mecklenburg-Vorpommern, Germany

161	Institute of Clinical Chemistry and Laboratory Medicine, University Medicine Greifswald, Greifswald, Mecklenburg-Vorpommern, Germany

162	Research/Psychiatry, Veterans Affairs San Diego Healthcare System, San Diego, CA, USA

163	Institute of Health and Biomedical Innovation, Queensland University of Technology, Brisbane, QLD, Australia

164	Department of Clinical Psychiatry, Psychiatry Clinic, Clinical Center University of Sarajevo, Sarajevo, Bosnia and Herzegovina

165	Department of Neurobiology, Care sciences, and Society, Karolinska Institutet and Center for Molecular Medicine, Karolinska University Hospital, Stockholm, Sweden

166	Humus, Reykjavik, Iceland

167	Mental Health Department, University Regional Hospital, Biomedicine Institute (IBIMA), Málaga, Spain

168	Virginia Institute for Psychiatric & Behavioral Genetics, Virginia Commonwealth University, Richmond, VA, USA

169	Solid Biosciences, Boston, MA, USA

170	Outpatient Clinic for Bipolar Disorder, Altrecht, Utrecht, Netherlands

171	Department of Psychiatry, Washington University in Saint Louis, Saint Louis, MO, USA

172	Department of Biochemistry and Molecular Biology II, Institute of Neurosciences, Center for Biomedical Research, University of Granada, Granada, Spain

173	Department of Neuroscience, Icahn School of Medicine at Mount Sinai, New York, NY, USA

174	Medicine, Psychiatry, Biomedical Informatics, Vanderbilt University Medical Center, Nashville, TN, USA

175	Department of Health Sciences Research, Mayo Clinic, Rochester, MN, USA

176	Psychiatry and Behavioral Sciences, Stanford University School of Medicine, Stanford, CA, USA

177	Rush University Medical Center, Chicago, IL, USA

178	Department of Psychiatry, University of Groningen, University Medical Center Groningen, Groningen, Netherlands

179	Scripps Translational Science Institute, La Jolla, CA, USA

180	Department of Psychiatry and Psychotherapy, Medical Center of the University of Munich, Campus Innenstadt, Munich, Germany

181	Division of Cancer Epidemiology and Genetics, National Cancer Institute, Bethesda, MD, USA

182	Behavioral Health Services, Kaiser Permanente Washington, Seattle, WA, USA

183	Faculty of Medicine, Department of Psychiatry, School of Health Sciences, University of Iceland, Reykjavik, Iceland

184	Div Mental Health and Addiction, Oslo University Hospital, Oslo, Norway

185	NORMENT, University of Oslo, Oslo, Norway

186	Institute of Health and Wellbeing, University of Glasgow, Glasgow, United Kingdom

187	Psychiatry and the Behavioral Sciences, University of Southern California, Los Angeles, CA, USA

188	Mood Disorders, PsyQ, Rotterdam, Netherlands

189	Research Division, Federal Institute for Drugs and Medical Devices (BfArM), Bonn, Germany

190	Centre for Addiction and Mental Health, Toronto, ON, Canada

191	Neurogenomics, TGen, Los Angeles, AZ, USA

192	College of Biomedical and Life Sciences, Cardiff University, Cardiff, United Kingdom

193	Institute of Epidemiology and Social Medicine, University of Münster, Münster, Nordrhein-Westfalen, Germany

194	Institute for Community Medicine, University Medicine Greifswald, Greifswald, Mecklenburg-Vorpommern, Germany

195	Medical Genetics Section, CGEM, IGMM, University of Edinburgh, Edinburgh, United Kingdom

196	Clinical Neurosciences, University of Cambridge, Cambridge, United Kingdom

197	Internal Medicine, Erasmus MC, Rotterdam, Zuid-Holland, Netherlands

198	Roche Pharmaceutical Research and Early Development, Neuroscience, Ophthalmology and Rare Diseases Discovery & Translational Medicine Area, Roche Innovation Center Basel, F. Hoffmann-La Roche Ltd, Basel, Switzerland

199	Psychiatry, Psychiatrisches Zentrum Nordbaden, Wiesloch, Germany

200	Neuroscience Research Australia, Sydney, NSW, Australia

201	Computational Sciences Center of Emphasis, Pfizer Global Research and Development, Cambridge, MA, USA

202	Department of Biostatistics, Princess Margaret Cancer Centre, Toronto, ON, Canada

203	Dalla Lana School of Public Health, University of Toronto, Toronto, ON, Canada

204	Psychological Medicine, Institute of Psychiatry, Psychology & Neuroscience, King's College London, London, United Kingdom

205	Department of Mental Health, Johns Hopkins University Bloomberg School of Public Health, Baltimore, MD, USA

206	Institute of Genetic Medicine, Johns Hopkins University School of Medicine, Baltimore, MD, USA

207	NORMENT, KG Jebsen Centre for Psychosis Research, Division of Mental Health and Addiction, Institute of Clinical Medicine and Diakonhjemmet Hospital, University of Oslo, Oslo, Norway

208	National Institute of Mental Health, Klecany, Czechia

209	Department of Psychiatry, University of Münster, Münster, Germany

210	Department of Psychiatry, University of Melbourne, Melbourne, VIC, Australia

211	Department of Psychiatry and Addiction Medicine, Assistance Publique - Hôpitaux de Paris, Paris, France

212	Paris Bipolar and TRD Expert Centres, FondaMental Foundation, Paris, France

213	UMR-S1144 Team 1: Biomarkers of relapse and therapeutic response in addiction and mood disorders, INSERM, Paris, France

214	Psychiatry, Université Paris Diderot, Paris, France

215	Psychiatry, University of Pennsylvania, Philadelphia, PA, USA

216	Centre for Integrative Biology, Università degli Studi di Trento, Trento, Trentino-Alto Adige, Italy

217	Department of Psychiatry and Psychotherapy, Medical Center, University of Freiburg, Faculty of Medicine, University of Freiburg, Freiburg, Germany

218	Division of Endocrinology, Children's Hospital Boston, Boston, MA, USA

219	Centre for Affective Disorders, Institute of Psychiatry, Psychology and Neuroscience, London, United Kingdom

220	Department of Psychiatry & Psychology, Mayo Clinic, Rochester, MN, USA

221	School of Medical Sciences, University of New South Wales, Sydney, NSW, Australia

222	Department of Human Genetics, University of Chicago, Chicago, IL, USA

223	Amsterdam Public Health Institute, Vrije Universiteit Medical Center, Amsterdam, Netherlands

224	Biometric Psychiatric Genetics Research Unit, Alexandru Obregia Clinical Psychiatric Hospital, Bucharest, Romania

225	Psychiatry, Kaiser Permanente Northern California, San Francisco, CA, USA

226	Medical Research Council Human Genetics Unit, Institute of Genetics and Molecular Medicine, University of Edinburgh, Edinburgh, United Kingdom

227	Institute of Neuroscience and Physiology, University of Gothenburg, Gothenburg, Sweden

228	INSERM, Paris, France

229	Neuroscience Therapeutic Area, Janssen Research and Development, LLC, Titusville, NJ, USA

230	Cancer Epidemiology and Prevention, M. Sklodowska-Curie Cancer Center and Institute of Oncology, Warsaw, Poland

231	School of Psychology, The University of Queensland, Brisbane, QLD, Australia

232	Research Institute, Lindner Center of HOPE, Mason, OH, USA

233	Human Genetics Branch, Intramural Research Program, National Institute of Mental Health, Bethesda, MD, USA

234	Division of Mental Health and Addiction, Oslo University Hospital, Oslo, Norway

235	Division of Mental Health and Addiction, University of Oslo, Institute of Clinical Medicine, Oslo, Norway

236	Institute of Molecular and Cell Biology, University of Tartu, Tartu, Estonia

237	Mental Health, Faculty of Medicine and Health Sciences, Norwegian University of Science and Technology - NTNU, Trondheim, Norway

238	Psychiatry, St Olavs University Hospital, Trondheim, Norway

239	Psychosis Research Unit, Aarhus University Hospital, Risskov, Denmark

240	Munich Cluster for Systems Neurology (SyNergy), Munich, Germany

241	University of Liverpool, Liverpool, United Kingdom

242	Psychiatry and Human Genetics, University of Pittsburgh, Pittsburgh, PA, USA

243	Mental Health Services in the Capital Region of Denmark, Mental Health Center Copenhagen, University of Copenhagen, Copenhagen, Denmark

244	Division of Psychiatry, Haukeland Universitetssjukehus, Bergen, Norway

245	Faculty of Medicine and Dentistry, University of Bergen, Bergen, Norway

246	Human Genetics and Computational Biomedicine, Pfizer Global Research and Development, Groton, CT, USA

247	College of Medicine Institute for Genomic Health, SUNY Downstate Medical Center College of Medicine, Brooklyn, NY, USA

248	Psychiatry, Harvard Medical School, Boston, MA, USA

249	Division of Clinical Research, Massachusetts General Hospital, Boston, MA, USA

250	Department of Clinical Genetics, Amsterdam Neuroscience, Vrije Universiteit Medical Center, Amsterdam, Netherlands

251	Department of Neurology and Neurosurgery, McGill University, Faculty of Medicine, Montreal, QC, Canada

252	Montreal Neurological Institute and Hospital, Montreal, QC, Canada

253	Department of Biomedical and NeuroMotor Sciences, University of Bologna, Bologna, Italy

254	Faculty of Medicine, University of Iceland, Reykjavik, Iceland

255	Department of Psychiatry, Hospital Namsos, Namsos, Norway

256	Department of Neuroscience, Norges Teknisk Naturvitenskapelige Universitet Fakultet for naturvitenskap og teknologi, Trondheim, Norway

257	Child and Adolescent Psychiatry, Erasmus MC, Rotterdam, Zuid-Holland, Netherlands

258	Psychiatry, Erasmus MC, Rotterdam, Zuid-Holland, Netherlands

259	Department of Psychiatry, McGill University, Montreal, QC, Canada

260	Dept of Psychiatry, Sankt Olavs Hospital Universitetssykehuset i Trondheim, Trondheim, Norway

261	Clinical Institute of Neuroscience, Hospital Clinic, University of Barcelona, IDIBAPS, CIBERSAM, Barcelona, Spain

262	Division of Epidemiology, New York State Psychiatric Institute, New York, NY, USA

263	Department of Clinical Medicine, University of Copenhagen, Copenhagen, Denmark

264	Biochemistry and Molecular Biology, Indiana University School of Medicine, Indianapolis, IN, USA

265	Department of Psychiatry, University of North Carolina at Chapel Hill, Chapel Hill, NC, USA

266	Department of Medical & Molecular Genetics, King's College London, London, United Kingdom

267	Department of Psychiatry, Indiana University School of Medicine, Indianapolis, IN, USA

268	Department of Genetics, University of North Carolina at Chapel Hill, Chapel Hill, NC, USA

† Deceased

### Supplementary Note

#### Relationship between these analyses and recent depression analyses from the UK Biobank

The PGC GWAS of major depression included data from the first 150,000 UK Biobank individuals whose genotypic data was released (1). Depression GWAS in the full UK Biobank cohort have since been published, including both broad and narrow definitions (2). The broad depression GWAS was meta-analysed with data from the PGC publication (3). We conducted a further meta-analysis of PGC and UK Biobank major depressive disorder data, using data from the online mental health phenotyping, including questions derived from the Composite International Diagnostic Interview – Short Form (4, 5). This phenotype has good concordance with direct clinical assessments of major depressive disorder and can be considered a major depressive disorder phenotype, compared to the less specific broad depression phenotype used by Howard et al (3, 6, 7). The effects on GWAS of using different depression phenotypes from the UK Biobank is investigated in depth elsewhere (8). We compare our results with those from Howard et al where appropriate (3).

### Supplementary Methods

#### Participants

The PGC MDD cohort consists of an anchor set of 29 cohorts (16,823 cases and 25,632 controls), with case individuals meeting international consensus criteria (DSM-IV, ICD-9, or ICD-10) for a lifetime diagnosis of major depressive disorder using structured diagnostic instruments. Six additional cohorts (118,635 cases and 319,269 controls) were drawn from broader population-based studies, and cases met criteria through self-report or responses to structured diagnostic instruments. Controls in most samples were screened for the absence of lifetime psychiatric disorders. All participants were of Western European ancestries. Individuals from the anchor cohort meet criteria for major depressive disorder. However, the additional cohorts include individuals who self-reported their diagnosis, and might not have met criteria for major depressive disorder in a clinical setting. In particular, these additional cohorts included data from 23andMe, where case participants were defined by a positive endorsement of a single question "Have you ever been diagnosed with clinical depression?" (or a different version of this question with similar phrasing) (9). These participants self-reported a professional diagnosis of depression, rather than being ascertained via a direct examination of all criteria for major depressive disorder. As such, it was considered more appropriate to refer to these individuals as having major depression, rather than major depressive disorder (1).

The PGC BD cohort consists of 32 studies (20,352 cases and 31,358 controls) of Western European ancestries. Case individuals were required to meet international consensus criteria for a lifetime diagnosis of bipolar disorder using structured diagnostic instruments. Controls in most samples were screened for the absence of lifetime psychiatric disorders.

In the UKB MDD cohort, participants were defined as cases if they met criteria based on questions derived from the Composite International Diagnostic Interview (CIDI). Participants were excluded if they self-reported previous diagnoses of schizophrenia (or other psychoses) or bipolar disorder. Controls were excluded if they self-reported any mental illness, reported taking any drug with an antidepressant indication, had previously been hospitalised with a mood disorder or met previously-defined criteria for a mood disorder (Supplementary Table 1) (10).

Quality control and imputation of UK Biobank data was performed centrally and is described elsewhere (11). Additional quality control was performed and is described in full elsewhere (5). In brief, participants were limited to unrelated individuals (KING correlation coefficient < 0.044 with all pairs, equivalent to removing all third degree-or-closer relatives) from probable Western European ancestries with good quality genotype data (passed Affymetrix and central UK Biobank quality assurance processes, genotyping call rate > 98%, concordant genotypic and phenotypic sex). Genome-wide association analyses (GWAS) of each UKB cohort were performed in BGenie v1.2, limited to variants with minor allele frequency (MAF) > 0.01 that were genotyped or imputed with confidence (IMPUTE2 INFO score > 0.4) (11, 12). All GWAS included six genotypic principal components (derived from the Western European ancestries subset of the UKBiobank using flashpca2) (13) and factors of genotyping batch and assessment centre as covariates to control for batch effects and population stratification. BGenie performs linear regressions on phenotypes residualised for covariates - as such, the resulting beta effect sizes (for all UKB analyses) were converted to odds ratios for meta-analysis using LMOR (14). Standard errors for the odds ratios were calculated by transforming the BGenie p-value to a Z score and dividing log(odds ratio) by Z (15).

#### Use of MAF > 0.05 as a cutoff

All summary statistics were limited to variants with MAF > 0.05. This was chosen because previous analyses of the BD2 subtype suggested that including lower MAF variants may bias SNP-based heritability estimates (16). Specifically, the BD2 subtype comprises multiple small cohorts, some with unbalanced case/control numbers. Consequently, potentially spurious effects in a single study can drive results for low-frequency variants. We therefore chose to remove lower frequency variants from all analyses in this paper.

#### Population prevalences

SNP-based heritability estimates were transformed to the liability scale assuming that the combined population prevalence between major depressive disorder and bipolar disorder is the sum of the disorder prevalences. Specifically, we assumed a population prevalence of 15% for combined MDD, 1% for PGC BD, and thus 16% for MOOD. Further estimates were made for the lower bounds of prevalence and upper bounds of prevalence for comparison. Namely, we set lower bounds of population prevalence at 10% for combined MDD, 0.5% for PGC BD and 10.5% for MOOD, and upper bounds at 20%, 2% and 22% for combined MDD, PGC BD and MOOD respectively.

#### Definition of non-overlapping N

As some analyses (particularly LD score regression) use the total number of subjects in the analysis for calculations, a "non-overlapping N" was estimated for each meta-analysis, using the following equation (derived from the equation describing the genetic covariance intercept in LD Score) (17):

Non-overlapping N = N1 + (N2 - (g_cov_int_ * √N1N2))

where N1 is the cohort size of the larger component part of the meta-analysis, N2 is the same for the smaller cohort and g_cov_int_ is the genetic covariance intercept from the calculation of genetic correlation between the component parts in LD Score. This method can be extended to the meta-analysis of three cohorts in a two-step process (calculating a non-overlapping N for cohorts one and two, and then the non-overlapping N for the meta-analysis of the combined one-two cohort with cohort three). In the case of MOOD herein, the non-overlapping N was calculated as if meta-analysing PGC MDD and PGC BD followed by UKB MDD; and as if meta-analysing PGC MDD and UKB MDD followed by PGC BD. The average of the two results was taken as the non-overlapping N. Note that this equation implicitly assumes that the phenotypic correlation between the traits of interest in the overlapping samples is 1 (which is reasonable in this instance). Note also that the resulting non-overlapping N is underestimated in the presence of shared confounding between the cohorts (such as through population stratification) (18).

#### Comparison of METACARPA with meta-analysis of independent cohorts

We contributed individuals from the UK Biobank to PGC MDD, and so were able to define the overlap between PGC MDD and UKB MDD (3,087 cases and 5,128 controls, representing 10% and 8% of UKB MDD respectively). To examine the robustness of meta-analysis using overlapping cohorts in METACARPA, we re-ran UKB MDD excluding these overlapping individuals and then meta-analysed the results with PGC MDD using inverse variance weighted meta-analysis in METAL. We calculated genetic correlations between the results using LDSC, and calculated Pearson's correlations between the betas and p-values of the two analyses. Results of these analyses were highly consistent (betas r = 0.99, p-values r = 0.98, LDSC r_g_ = 1), suggesting METACARPA can adjust adequately for overlap between cohorts.

#### Definition of GWAS loci

GWAS results were clumped using PLINK1.9, assigning nominally-significant (p < 10^-4^) variants to a clump if they were in linkage disequilibrium (r^2^ > 0.1 in unrelated non-Finnish European participants from 1000 Genomes project) with a variant with a lower p-value lying within 3Mb (19, 20). Non-Finnish Europeans were used for the LD reference panel because they were the best match for the participants included in this analysis, who were of predominantly Western European ancestries. Loci were declared genome-wide significant if a variant in the locus reached the conventional threshold for genome-wide significance (p = 5 x 10^-8^). Results were visualised using FUMA, including mapping the loci to potentially affected genes through expression quantitative trait loci (eQTLs) and chromatin contact sites in brain tissues or neural progenitor cells (21).

All genomic loci reaching genome-wide significance in at least one analysis were combined for annotation. Where loci from different GWAS overlapped, they were combined into a single locus ranging from the minimum base position from any of the constituent loci to the maximum. Annotation was performed using RegionAnnotator version 1.63 (<https://github.com/ivankosmos/RegionAnnotator>), which includes data from: the NHGRI-EBI GWAS Catalog; OMIM; GENCODE genes; genes previously implicated in autism and in intellectual disability; copy-number variants previously implicated in psychiatric disorders; and mouse knockout phenotypes. Results from the GWAS Catalog module of RegionAnnotator were filtered to include only variants reaching genome-wide significance. Where multiple variants are listed as significant from a previous GWAS of a specific phenotype, only the variant with the lowest p-value is reported. Results from the GWAS Catalog module were supplemented by direct query of the NHGRI-EBI GWAS Catalog for each region (data from 2018-11-05), lookup of top SNPs in <http://atlas.ctglab.nl>, and manual assessment of psychiatric and behavioural GWAS not yet listed in the NHGRI-EBI GWAS Catalog.

#### Relationship between meta-analyses (hierarchical clustering)

The relationship between the loci identified in the meta-analyses (MOOD, combined MDD) and the constituent analyses (PGC MDD, UKB MDD, PGC BD) was assessed using a hierarchically-clustered heatmap, with the 2014 PGC schizophrenia analysis (SCZ) included for comparison (22). For the purpose of this comparison, an index SNP was selected for each locus (the variant with the lowest p-value that was common to all four analyses) to obtain the direction of effect in each analysis. Index SNP p-values were converted to -log_10_(p-value). If the OR_index_ < 1 in a given analysis, the log_10_(p-value) was used in place of the -log_10_(p-value). A hierarchically-clustered heatmap was then generated using the default options of the heatmap.2 function of the gplots package in R v1.4.1 (complete clustering on the Euclidean distance between vectors) with clustering performed separately for rows and for columns (23, 24).

#### Hierarchical clustering of genetic correlations between subtypes, and extension to examine relationships with related traits

Genetic correlations between the major depressive disorder and bipolar disorder subtypes were hierarchically clustered using the method described above. In addition, we included results from six external GWAS relevant to mood disorders. We examined relationships with anxiety disorders (correlated with depressive phenotypes), schizophrenia (correlated with bipolar disorder), and ADHD (showed differing genetic correlations with PGC MDD and PGC BD) (22, 25, 26). We also examined subjective wellbeing, which may reflect positive mood, and included measures of specific aspects of wellbeing, namely eudaimonic wellbeing (feeling life has meaning) and hedonic wellbeing (feeling happy) (27, 28). For hierarchical clustering only, the sign of genetic correlations with the three wellbeing phenotypes were reversed so that positive effect sizes meant poor outcomes for all phenotypes.

We concluded that the genetic correlations between subtypes of major depressive disorder and bipolar disorder were indicative of a genetic mood disorder spectrum. To validate this conclusion, we performed a principal component analysis of the genetic correlation matrix as used for hierarchical clustering above (that is, including the six external GWAS with correlations with the wellbeing phenotypes reversed). Principal component analysis was performed using the prcomp function from base R, and using plot3D (https://cran.r-project.org/package=plot3D) for visualisation (23).

#### Conditional and reversed-effect analyses

Analyses were performed to understand which genomic loci are shared or distinct between the disorders, using mtCOJO, an extension of the GSMR method implemented in GCTA (29, 30). mtCOJO adjusts the results of a genome-wide association analysis, conditioning on the effects of a set of significantly associated, independent variants from a second set of summary statistics (a putative instrumental variable). This putative instrumental variable is also used as a proxy for the trait of interest to infer causal direction in GSMR analyses. The effect size estimated by this method is robust to confounding caused by genetic or environmental effects shared between the studies analysed, assuming these are uncorrelated with the instrumental variable. mtCOJO adjusts for sample overlap (i.e. the same individuals being present in both datasets) using the genetic covariance intercept from LD score regression (29, 31).

Conditional analyses in mtCOJO were performed on combined MDD conditional on PGC BD (MDDcBD), and on PGC BD conditional on combined MDD (BDcMDD). Variant selection for conditioning was performed with the default settings in mtCOJO, clumping using the UK Biobank dataset, and selecting at least ten variants with p<5x10^-8^, which were not in linkage disequilibrium (r^2^ < 0.05) with a variant with a lower p-value, and which did not show evidence of pleiotropy (passed the HEIDI-outlier analysis, threshold 0.01) (29). As down-sampled MDD had only eight variants with p<5x10-8, conditional analyses and GSMR were not performed using this meta-analysis. Similarly, BDcMDD had only six variants passing genome-wide significance, so GSMR analyses with BDcMDD as the exposure were not possible. Results were clumped in PLINK using the procedure described above. Genetic correlations were compared between combined MDD and MDDcBD, and between PGC BD and BDcMDD, first using a z-test to identify putative differences (p < 0.05), and then formally testing by applying a block-jackknife (described below). A conservative Bonferroni correction was used to determine significance (p < 1.27x10^-4^, approximate correction for 414 tests) (32–34).

A further analysis was performed to identify loci with opposite directions of effect between combined MDD and PGC BD. For this analysis, the direction of effects for the PGC BD analysis was reversed, and the MOOD meta-analysis repeated as described in the main text.

#### GSMR

GSMR uses the HEIDI test to remove pleiotropic variants from the instrument variable set. All analyses were run using the default settings in GSMR, assuming at least ten linkage-independent (r^2^ < 0.05) significant (p<5x10^-8^) variants pass the HEIDI test (threshold p<0.01).

#### Comparing two genetic correlations using jackknife and LDSC

Let there be four phenotypes A, B, C, and D. The goal is to compare the genetic correlation between A and B to the genetic correlation between C and D. Global estimates of these correlations can be computed using the LDSC software and will be noted *r(A,B)* and *r(C,D)*. The same software can output jackknife delete values for genetic covariance: G(A,B), G(C,D), as well as for heritability: H(A,B) and H(C,D). These jackknife delete values are estimated by excluding blocks of values (here, number of blocks *n* = 200). The *n*-dimensional vectors G(A,B), G(C,D), H(A,B) and H(C,D) can be used to generate genetic correlation delete values R(A,B) and R(C,D). The difference between the global estimates *r(A,B)* and *r(C,D)* is *d(AB,CD)*, and the difference between the vectors R(A,B) and R(C,D) is D(AB,CD). The global genetic correlation difference *d(AB,CD)* and the delete values D(AB,CD) are used to compute jackknife pseudovalues. The *i*th pseudovalue is

$$P_{i}(AB,CD)=n\times d(AB,CD)-(n-1)*D_{i}(AB,CD)$$

The mean and variance of the jackknife pseudovalues are

$$m(AB,CD)=\frac{1}{n}\sum_{i=1}^{n} P_{i}(AB,CD)$$

$$v(AB,CD)=\frac{1}{n-1}\sum_{i=1}^{n} {(P_{i}(AB,CD)-m(AB,CD))}^{2}$$

The jackknife estimate of the difference between the two correlations *m(AB,CD)* can then be compared to test H_0_ : *θ* = *θ_0_* (*θ_0_* = 0 for no difference between genetic correlations), and a p-value can be derived from the z statistic:

$$z(AB,CD)=\frac{m(AB,CD)-\theta_{0}}{\sqrt{(1/n)\times v(AB,CD)}}$$

#### Estimating local SNP-based heritability and genetic covariance in HESS

Local estimates of SNP-based heritability and genetic correlation were obtained using HESS v0.5.3b (35, 36). All analyses used the reference panel provided with the software (1000 Genomes Project European individuals) and previously defined blocks of the genome in linkage equilibrium, with 50 eigenvectors for inverting the LD matrix and a minimum eigenvalue cutoff of 1 (37, 38). Overlap between cohorts was calculated consistent with the calculation of non-overlapping N (see above), assuming overlapping individuals had a phenotypic correlation of 1 (that is, overlapping individuals were controls in both studies). Local heritability estimates were calculated for all meta-analyses, component GWAS and subtypes. Local genetic covariance was calculated between combined MDD and PGC BD, between MOOD and down-sampled MOOD, and between all of the major depressive disorder and bipolar disorder subtype pairwise.

#### Gene-wise and gene-set enrichment analyses

For all analyses, gene-wise p-values were calculated as the aggregate of the mean and smallest p-value across all SNPs annotated to Ensembl gene locations using MAGMA v1.06 (using the build 37 reference supplied on the MAGMA website) (39). SNPs were assigned to genes if they lay between 35kb upstream and 10kb downstream of the gene location (40). MAGMA accounts for possible confounders such as gene size, gene density, linkage disequilibrium and minor allele count. The threshold for genome-wide significance was defined at p < 2.6x10^-6^ (Bonferroni correction for the 19,041 genes tested). Genes passing genome-wide significance were defined as coming from the same locus if they lay within 100kb of each other, or if they overlapped a locus from the single variant analysis. Gene set analysis was performed in MAGMA for 13,567 gene sets. Significance was set at a Bonferroni-corrected threshold of p = 5.34x10^-6^ for 9,361 effectively independent tests within each analysis.

Gene set analysis was performed for all analyses. A gene set matrix **P** was generated with elements *P_g,p_* = 1 if gene *g* was in set *p* and *P_g,p_* = 0 otherwise. Association between gene set membership and gene-wise z-scores was computed using MAGMA. 13,567 gene sets were drawn from OpenTargets (downloaded January 2017) (41), GO ontologies, canonical gene sets drawn from MSigSB v5.2 C2 and C5 datasets (42), and biological gene sets related to psychiatric disorders described in various scientific publications. There is considerable overlap of genes between gene sets (within and between sources). Gene sets that overlapped entirely were treated as a single gene set in analyses. The effective number of gene sets tested was defined as the number of principal components accounting for 99.5% of explained variance in the gene set similarity matrix, obtained by computing the Tanimoto similarity between gene sets. This results in a Bonferroni-corrected threshold of p = 5.34x10^-6^ for 9,361 effectively independent tests for each matrix.

#### Tissue and single-cell enrichment analyses

Further analyses were performed to assess the enrichment of associated genes with expression-specificity profiles from tissues (version 7 data from the Genotype-Tissue Expression project) and broadly-defined ("level 1") and narrowly-defined ("level 2") cell-types (Karolinska Institutet mouse brain single-cell RNA sequencing superset) (43, 44). Analyses were performed in MAGMA following previously described methods with minor modifications (44). Briefly, for the single cell data set (44), gene expression for each cell type was scaled to 1,000,000 unique molecular identifiers prior to computing specificity scores. Specificity scores are defined as the proportion of the total expression of a specific gene found in a given cell type. For the GTEx dataset, transcripts per million (TPM) were transformed to log_2_(TPM +1) prior to computing specificity scores. For each tissue or cell-type, the specificity scores were then rank-transformed to a standard normal distribution using the *rntranform* function from the GenABEL R package (45). The standard normalised specificity scores were then regressed on gene-wise association in the meta-analysis, defined as the mean p-value across all SNPs assigned to the gene. Multiple-testing correction was applied using Bonferroni-correction within each analysis.

#### Association with predicted brain tissue gene expression

Variant-level meta-analysis results were used to predict gene expression using S-PrediXcan and genomic and transcriptomic reference data from the thirteen brain regions assayed in the GTEx project (version 7) (43, 46). Associations were calculated between these predicted gene expression levels and each meta-analysed phenotype. Significance was set at 8.5x10^-8^, the Bonferroni correction for 586,469 tests (45,113 genes across 13 tissues) as in the original S-PrediXcan publication (46). Genes were defined as coming from the same locus following the approach described for MAGMA analyses.

#### Polygenic risk score prediction of UKB subtypes using PGC BD summary statistics

In order to determine if the recurrent major depressive disorder subtype (rMDD) was genetically more similar to PGC BD than were other major depressive disorder subtypes (single episode major depressive disorder, sMDD;and subthreshold depression, subMDD), polygenic risk score analyses were performed using PRSice2 (47). PGC BD results were used as the base analysis to produce polygenic risk scores (PRS) in the genotyped data from the UKB sample, and these were then compared across the major depressive disorder subtypes using logistic regression (including the covariates described above for the UKB GWAS). PRS were derived using linkage-independent (r^2^ < 0.1, ± 250kb) variants at seven p-value thresholds from the PGC BD data (pT = 0.001, 0.05, 0.1, 0.2, 0.3, 0.4, 0.5). Correction for multiple-thresholding was performed by using 20000 permutations (using the permutation function in PRSice2) to produce an empirical p-value (minimum possible empirical p = 5x10^-5^). Variance explained was initially calculated as Nagelkerke pseudo-R^2^ (using the fmsb package in R) and was subsequently converted to liability scale using http://cnsgenomics.com/shiny/abc/ (48, 49). Population prevalences for each subtype were set as follows: rMDD = 0.05; sMDD = 0.15; subMDD = 0.2. The population prevalence for each comparison was then calculated as each prevalence divided by the summed prevalence to give the following: rMDD vs sMDD = 0.25 (that is, 0.05 / [0.05+0.15] ); rMDD vs subMDD = 0.2; sMDD vs subMDD = 0.429.

### Supplementary Results

#### Conditional analyses

We performed analyses of combined MDD conditioning on PGC BD (MDDcBD). Diminished effects (shrinkage of the odds ratio shrinkage towards 1) were observed in 51/63 loci reaching genome-wide significance in combined MDD, suggesting most loci significantly associated with combined MDD have the same direction of effect in PGC BD (Supplementary Table 13). Results from the reverse analysis (PGC BD conditioned on combined MDD: BDcMDD) support this conclusion, with 14/19 associated loci from PGC BD showing a diminished effect (Supplementary Table 13).

The SNP-based heritability of the conditional analyses showed reduced estimates compared to the respective main analyses (MDDcBD: 7%, combined MDD: 9%, BDcMDD: 17%, PGC BD: 20%, Supplementary Table 2). Genetic correlations for MDDcBD mirror those for combined MDD, except that genetic correlations with schizophrenia (and related analyses, such as the schizophrenia-bipolar disorder meta-analyses) were significantly smaller (Supplementary Figure 12, Supplementary Tables 5 and 14) (22, 50). The genetic correlations from BDcMDD were similar to PGC BD, with significant reductions only with studies of depression and anxiety (Supplementary Tables 15 and 15) (26, 51, 52).

In addition to conditional analyses, we reversed the observed effects from PGC BD and meta-analysed with combined MDD. 32 loci were significant in the resulting MOOD BD Reversed analysis (Supplementary Table 3). All loci were strongly associated with combined MDD (max p = 2x10^-7^), 27 passing genome-wide significance, compared with only one locus passing significance in PGC BD. 19 loci showed consistent direction of effect between combined MDD and PGC BD, indicating that these loci were driven by strong associations with combined MDD, rather than having a differing effect on major depressive disorder than on bipolar disorder. The smallest p-value observed in PGC BD for any of the 13 loci with differing directions of effect was p=6x10^-6^, for locus Rev2 on chromosome 7. This locus contains the *CTTNBP2* gene, rare variants in which have suggestive evidence for implication in autism spectrum disorder (53).

Down-sampled reversed analyses yielded three loci passing genome-wide significance (Supplementary Table 3). All of these loci passed genome-wide significance in down-sampled MDD and had shared directions of effect between PGC BD and down-sampled MDD, indicating these loci do not have a differing effect on major depressive disorder than on bipolar disorder.

Tissue and cell-type expression specificity analyses showed high consistency between the main and the conditional analyses (Supplementary Tables 9-11). All brain tissues were enriched in both conditional analyses. In cell-type analyses, neuroblasts, adult dopaminergic neurons, embryonic GABAergic and midbrain nucleus neurons were significantly enriched in MDDcBD and not in BDcMDD. Conversely, medium spiny neurons (as well as both sets of pyramidal cells and the striatal interneurons) were significantly enriched in BDcMDD and not in MDDcBD.

GSMR analyses also showed high consistency between the main and the conditional analyses, although these analyses were limited because BDcMDD had too few variants with p<5x10^-8^ and so could only be used as an outcome, not an exposure, in GSMR analyses (Supplementary Figure 13, Supplementary Tables 12). MDDcBD had no significant relationship with PGC BD (nor BDcMDD with combined MDD), suggesting that conditioning was effective at removing the bidirectional relationship seen between combined MDD and PGC BD. Otherwise, results observed for combined MDD were also observed for MDDcBD, except that the positive association of major depressive disorder on CAD was attenuated and did not pass significance in MDDcBD. Results observed for PGC BD (as an outcome) were also observed for BDcMDD.

#### Gene-wise and gene set analyses

Gene-wise association analyses in MAGMA identified 361 genes associated with the MOOD phenotype at p < 2.6x10-6 (Supplementary Table 16). Associated genes were distributed across 120 loci, including 47 of the loci identified in MOOD. However, proximity is only a weak indication of the association of a gene with a trait (54). More evidence is provided by the convergence of brain-derived eQTL and chromatin contact data from a locus onto a single gene (Supplementary Figures 14-33 - note no figures are provided for chromosomes 8 or 21, as there are no significant loci on these chromosomes). In MOOD, such evidence suggested significant loci may act on *NEGR1* (loci 2 and 3), *RSRC1* (locus 14), *TMEM161B* (locus 18), *LHX2* (locus 41), *SOX5* (locus 53), *LACC1* (locus 56), *PCDH8* (locus 57) and *ZC2HC1C* (locus 61). However, diverse eQTL and chromatin contacts were observed at many of these loci, suggesting these associations may act through other genes as well.

Results from gene set analysis were generally similar between combined MDD and PGC BD, and in each of the conditional analyses (Supplementary Table 17). Gene sets significantly enriched across all analyses included genes previously implicated in schizophrenia, targets of the RNA splicing proteins *CELF4* and *RBFOX1/RBFOX3*, loss-of-function intolerant (pLI09) genes, and genes with products potentially involved in synaptic processes (Supplementary Table 17). Certain gene sets were enriched in one disorder only - for example, RBFOX2 targets were significantly enriched in combined MDD, but not in PGC BD. In contrast, gene sets annotated as mutation-intolerant (constrained and genic intolerance RVIS) were significantly enriched in PGC BD but not combined MDD. In the conditional analyses, results for combined MDD and MDDcBD were similar, with significantly associated gene sets falling into broad categories of psychiatrically associated, neurodevelopmental, and anthropometric gene sets (Supplementary Table 17). Fewer significant gene sets were observed in BDcMDD than in PGC BD, but included mutation-intolerant gene sets (Supplementary Table 17).

#### Local SNP-based heritability and genetic covariance

Genome-wide SNP-based heritability estimates on the observed scale were similar between LDSC and HESS for all main meta-analyses (Supplementary Table 18). For both MOOD and combined MDD, local SNP-based heritability was significantly >0 in the region overlapping loci 2 and 3 (near *NEGR1*), and for multiple regions comprising locus 25 (the major histocompatibility locus; p < 2.94x10^-5^, Bonferroni correction for 1703 LD-independent regions). No regions had significant local SNP-based heritability for PGC BD. Combined MDD and PGC BD were significantly genetically correlated (0.29, compare 0.35 from LDSC; Supplementary Table 19), but no regions had local genetic covariance that significantly differed from 0 (p > 2.94x10^-5^).

The observed SNP-based heritability of down-sampled MOOD from HESS was 11% (compare LDSC 8%; Supplementary Tables 2 and 18). Only one region, part of locus 25, had local SNP-based heritability significantly >0. One region on chromosome 10 had local SNP-based heritability significantly >0 in down-sampled MDD. However, this is probably a false positive, there are no variants significantly associated with down-sampled MDD in the region. In addition, this region encompasses the centromere of chromosome 10, which may result in the LD structure of the region being specified incorrectly.

#### Genetic correlations of mood disorder subtypes

Genetic correlations between the bipolar disorder and major depressive disorder subtypes suggest a spectrum of genetic relationships between major depressive disorder and bipolar disorder, with BD2 bridging the two disorders (Supplementary Figure 34). Adding in six external phenotypes resulted in two clusters, with two sets of intermediate phenotypes (Supplementary Table 10, Supplementary Figures 35-40). Major depressive disorder subtypes cluster with anxiety disorders and the wellbeing spectrum, albeit with negative genetic correlations with wellbeing. The relationship of the wellbeing spectrum with depressive disorders was captured more effectively by hedonic rather than eudaimonic wellbeing - however, neither of these wellbeing subtypes clustered with depressive disorders, reinforcing previous conclusions that wellbeing is multidimensional (55). In contrast, schizophrenia clusters with schizoaffective bipolar disorder and bipolar disorder type 1, consistent with the greater genetic similarity of these subtypes to schizophrenia (16, 56). ADHD has a moderate genetic correlation with bipolar disorder type 2, but not with the other bipolar disorder subtypes, arguing that the weaker genetic correlation between ADHD and bipolar disorder (compared to major depressive disorder) is specific to type 1 bipolar disorder.

Principal component analysis identified three principal components accounting for >90% of the variance in the genetic correlation matrix (Supplementary Figure 41). The first principal component accounted for 59% of the variance, and described the spectrum as proposed above, separating the cluster of schizophrenia, schizoaffective bipolar disorder, and bipolar disorder type 1 from the cluster of the depressive disorders, with bipolar disorder type 2 and ADHD intermediate between the two. The second (21% variance explained) principal component separated the eudaimonic and hedonic wellbeing phenotypes from the other phenotypes, and the third principal component (12% variance explained) separated ADHD from the other phenotypes (Supplementary Figure 41).

Estimates of local heritability (Supplementary Table 18) and genetic covariance (Supplementary Table 19) were calculated in HESS to assess whether specific regions of the genome were shared or distinct between subtypes. However, with the exception of BD1 and rMDD, SNP-based heritability estimates on the observed scale from HESS did not differ significantly from 0 (BD1 = 33%; BD2 = 3%; SAB = 0.4%; rMDD = 9%; sMDD = 0%; subMDD = 0.1%). This most likely resulted from the small cohort size of the subtype analyses, which results in a downward bias in SNP-based heritability estimation in HESS (36). No regions had significant local SNP-based heritability for any of the subtypes (all p > 2.94x10-5, Bonferroni correction for 1703 LD-independent regions). BD1 and rMDD were genetically correlated (0.36, compare 0.31 from LDSC; Supplementary Table 10), but no region had a genetic covariance significantly >0 (all p > 2.94x10-5).

#### Genetic correlations of PGC MDD and PGC BD with mood disorder subtypes

The genetic correlation of PGC MDD and BD2 was stronger than those with BD1 (Δr_g_ [difference between r_g_ estimates] = 0.39, p = 1x10^-4^) and with SAB (Δr_g_ = 0.52, p = 2x10^-5^), but the genetic correlations with BD1 and with SAB were not significantly different (Δr_g_ = 0.14, p = 0.05). PGC BD had a stronger genetic correlation with rMDD than with subMDD (Δr_g_ = 0.27, p = 3x10^-5^), but the genetic correlation between PGC BD and sMDD was not significantly different to those with rMDD (Δr_g_ = -0.07, p = 0.5) nor with subMDD (Δr_g_ = 0.20, p = 0.009).

#### Results from PGC MDD + subtype meta-analyses

##### Bipolar disorder subtypes

64 loci reached genome-wide significance across the meta-analyses between PGC MDD and the bipolar disorder subtypes (MDD-BD1, MDD-BD2, and MDD-SAB; Supplementary Table 3). Of these, 54 also reached significance in MOOD. The ten remaining loci were significant in MDD-BD2 alone (four loci), in MDD-BD2 and in MDD-SAB (three), in MDD-BD1 and in MDD-BD2 (one), in MDD-BD1 (one), and in MDD-SAB (one). Gene-wise association analyses in MAGMA identified 272 genes associated at p < 2.6x10^-6^ in at least one of the meta-analyses (Supplementary Table 16). Associated genes were distributed across 86 loci, including 44 of the loci identified by at least one single variant meta-analysis.

Heritability estimates for the meta-analyses between PGC MDD and the bipolar disorder subtypes were all very similar, ranging from 8-10% (assuming a lower bound of population prevalence of 10.5%, and an upper bound of 22%; Supplementary Table 2). Genetic correlations with previously-published traits were broadly similar across the different meta-analyses, and mirrored those from the main MOOD meta-analysis (psychiatric and behavioural, reproductive, and sociodemographic traits; Supplementary Table 5).

##### Major depressive disorder subtypes

65 loci reached genome-wide significance across the meta-analyses between PGC MDD and the major depressive disorder subtypes (MDD-rMDD, MDD-sMDD, and MDD-subMDD; Supplementary Table 3). Of these, 52 also reached significance in the MOOD analysis - the remaining 13 loci were significant in MDD-rMDD alone (four loci), in MDD-subMDD alone (four), in MDD-rMDD and MDD-subMDD (three), in all three analyses (one) and in MDD-sMDD and MDD-subMDD alone (one). Gene-wise association analyses in MAGMA identified 261 genes associated at p < 2.6x10^-6^ in at least one of the meta-analyses (Supplementary Table 16). Associated genes were distributed across 93 loci, including 45 of the loci identified by at least one single variant meta-analysis.

Heritability estimates for the meta-analyses between PGC MDD and the major depressive disorder subtypes were all similar, ranging from 7-10% (assuming a lower bound of population prevalence of 10%, and an upper bound of 20%), although estimates for MDD-rMDD were slightly higher than those for MDD-sMDD and MDD-subMDD (Supplementary Table 2). Genetic correlations with previously-published traits were broadly similar across the different meta-analyses, and mirrored those from the main MOOD meta-analysis (psychiatric and behavioural, reproductive, and sociodemographic traits; Supplementary Table 5).

#### Gains in discovery through adding individuals with different mood disorder diagnoses

The PGC MDD analysis was meta-analysed with PGC BD and UKB MDD cohorts and with the subtypes of both bipolar disorder and major depressive disorder. The relative increase in mood-disorder associated loci obtained by adding 1000 effective cases from different definitions to the PGC MDD GWAS was assessed. Effective cases were defined as half of the effective N (2 / [[1+Cases] + [1+Controls]]) (57). The resultant increase in locus discovery per 1000 effective cases of UKB MDD, PGC BD and each subtype is described in Supplementary Table 20. With the exception of SAB (the power of which is very low), meta-analysis with PGC MDD resulted in an increased number of loci in all cases, with BD2 providing the most additional loci per 1000 effective cases (0.67). BD1 cases provided a similar amount of additional loci to rMDD (0.5 vs 0.51), and both out-performed sMDD (0.2). This suggests BD1 cases may function in a similar manner to rMDD cases in meta-analysis with PGC MDD, while BD2 cases appear to be equivalent to more extreme rMDD cases. As expected, like-for-like, rMDD cases provide more loci than sMDD cases, most likely due to increased heterogeneity of sMDD cases (potentially because single depressive episodes may be more likely to represent a reaction to a specific event). This fits with the higher heritability of recurrent major depressive disorder (45%) versus single episode major depressive disorder (34%) (58). These conclusions mirror the increase in mean-chi-square of each meta-analysis compared to PGC MDD alone (Supplementary Table 20). However, limitations remain, including the unknown effects of error (such that it is difficult to assess the meaning of differences between subtypes robustly) and the fact that the major depressive disorder subtypes are drawn from the same source (UK Biobank), which may differ from clinically-ascertained major depressive disorder cohorts, both in heterogeneity and in severity (4, 59).

#### Associations of polygenic risk scores for PGC BD with UKB MDD subtypes

Polygenic risk scores derived from the PGC BD analysis (Supplementary Table 21, Supplementary Figure 41) were significantly positively associated with rMDD when compared to sMDD (p = 3x10^-16^; empirical p = 5x10^-5^) and when compared to subMDD (p = 3x10^-19^; empirical p = 5x10^-5^). In contrast, the association between the PGC BD risk score and sMDD compared to subMDD was not significant when taking into account the multiple thresholds tested (p = 0.04; empirical p = 0.13). Grouping rMDD and sMDD together as UKB MDD cases, the PGC BD risk score was significantly positively associated with UKB MDD cases compared to controls (p = 6x10^-40^; empirical p = 5x10^-5^). Taken together, these results suggest that rMDD has more in common genetically with PGC BD than does sMDD. This mirrors previous findings that showed BD2 was more similar genetically to PGC MDD than was BD1 (16).

#### Relationship of meta-analysis results (hierarchical clustering)

Hierarchical clustering of the significant loci from MOOD and combined MDD with the same loci from PGC MDD, UKB MDD, and PGC BD (and the PGC2 SCZ for comparison) (22) resulted in MOOD clustering most closely with PGC MDD. This indicates that the primary contribution to significant loci in the meta-analysis came from PGC MDD rather than PGC BD (Supplementary Figure 42). Results from UKB MDD for these loci clustered closer to PGC BD than to PGC MDD, suggesting that the component analyses cluster primarily by their contribution to the meta-analysis, rather than by trait. Despite this similarity between UKB MDD and PGC BD when considering genome-wide significant loci, comparisons of the SNP-heritability of the component analyses (Supplementary Table 2) and of the genetic correlations between them (Supplementary Table 10) confirm that UKB MDD is more similar in general to PGC MDD than to PGC BD.

#### Sensitivity analysis - Equivalently-powered cohorts

Summary statistics were available from the PGC comprising the PGC MDD cohort without the inclusion of the 23andMe and the original UK Biobank cohorts. The mean chi-square of the meta-analysis between the down-sampled PGC MDD and UKB MDD was 1.35 (compared to 1.70 in combined MDD), similar to the mean chi-square of PGC BD and therefore suitable for the purpose of the sensitivity analysis (Supplementary Table 2). We therefore meta-analysed down-sampled PGC MDD, PGC BD and UKB MDD (down-sampled MOOD; cases = 95,418, controls = 192,514, non-overlapping N = 280,214).

19 loci reached genome-wide significance, of which 17 were present in MOOD and two did not reach significance in any of the main analyses (Supplementary Table 3). Hierarchical clustering of the significant loci from down-sampled MOOD with the same loci from down-sampled PGC MDD, UKB MDD, and PGC BD (and the PGC2 SCZ for comparison) (22) resulted in all three component analysis clustering together, with the meta-analysis clustering separately (Supplementary Figure 43). This suggests that the clustering of PGC MDD with MOOD in the main paper resulted from the power difference. UKB MDD again clustered more closely with PGC BD than with down-sampled PGC MDD, although the genetic correlation of UKB MDD with down-sampled PGC MDD (r_g_ = 0.86) was still greater than with PGC BD (r_g_ = 0.34).

Nine of the 44 PGC MDD loci reached genome-wide significance in the down-sampled MOOD meta-analysis (20% of all PGC MDD loci), as did both of the loci that reached genome-wide significance in down-sampled PGC MDD (Supplementary Table 3). In comparison, only two of the 19 PGC BD loci reached genome-wide significance in the meta-analysis (11%), suggesting that the addition of individuals with bipolar disorder to major depressive disorder cohorts still appears to enrich more for associations with major depressive disorder than for bipolar disorder.

Two loci reaching genome-wide significance in down-sampled MOOD did not reach genome-wide significance in MOOD, of which one reached significance in PGC BD. The other is a multi-gene locus on chromosome 3 that has reached genome-wide significance in a wide variety of traits, including depressive symptoms (60). In addition to this locus, a further seven loci reached genome-wide significance in down-sampled MOOD that did not reach significance in PGC MDD or PGC BD including locus 30, near *PCLO* (but not locus 51, near *DRD2*).

The estimate of SNP-heritability for down-sampled MOOD (11% with population prevalence 16%) was greater than that for MOOD (8.8%), but still remains more similar to PGC MDD (9%) than to PGC BD (17-23%; Supplementary Table 2) (1, 16). Similarly, the genetic correlations between down-sampled MOOD and other traits broadly recapitulated those for MOOD (Supplementary Table 5). Significantly greater correlations (compared with MOOD) were seen between down-sampled MOOD and bipolar disorder, schizophrenia, combined analyses of bipolar disorder and schizophrenia, and the cross-disorder analysis, while reduced correlations were seen with PGC MDD and anxiety (Supplementary Table 7). Interestingly, a significant negative correlation with IQ was observed (r_g_ = -0.13, p = 5x10^-7^), which was not observed in MOOD, PGC MDD nor PGC BD. Further investigation of this genetic correlation revealed that the 23andMe depression cohort has a positive genetic correlation with IQ (r_g_ = 0.06, p = 0.01); including this cohort in the PGC MDD sample obscured a negative genetic correlation with IQ.

Overall, the sensitivity analyses suggest that the difference in power between combined MDD and PGC BD does contribute to the greater similarity of MOOD to PGC MDD than to PGC BD. However, the pervasive similarity to PGC MDD seen in the down-sampled analysis suggests the results seen in the main analysis are not just a consequence of the power difference.

The observed SNP-based heritability of down-sampled MOOD from HESS was 11% (compare LDSC 8%; Supplementary Tables 2 and 18). Only one region, part of locus 25, had local SNP-based heritability significantly >0 in down-sampled MOOD. One novel region on chromosome 10 had local SNP-based heritability significantly >0 in down-sampled MDD (and in the down-sampled PGC MDD) – however, no significant variants were observed in this region, so this may be spurious. Down-sampled MDD and PGC BD were significantly genetically correlated (0.33, compare 0.37 from LDSC), but no regions had local genetic covariance that significantly differed from 0 (all p > 2.94x10^-5^; Supplementary Table 19).

### Supplementary Figures

#### Supplementary Figure 1

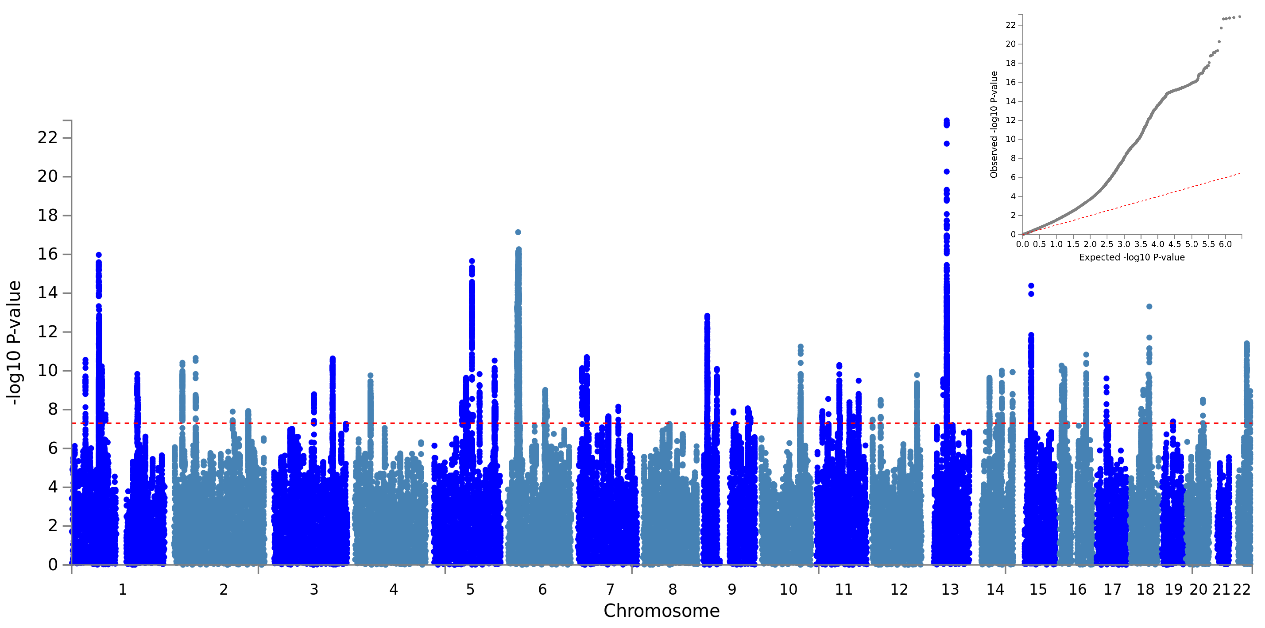

Supplementary Figure 1: Manhattan and QQ plot of results for the mood disorders (MOOD) meta-analysis. Red line is 5x10^-8^

#### Supplementary Figure 2

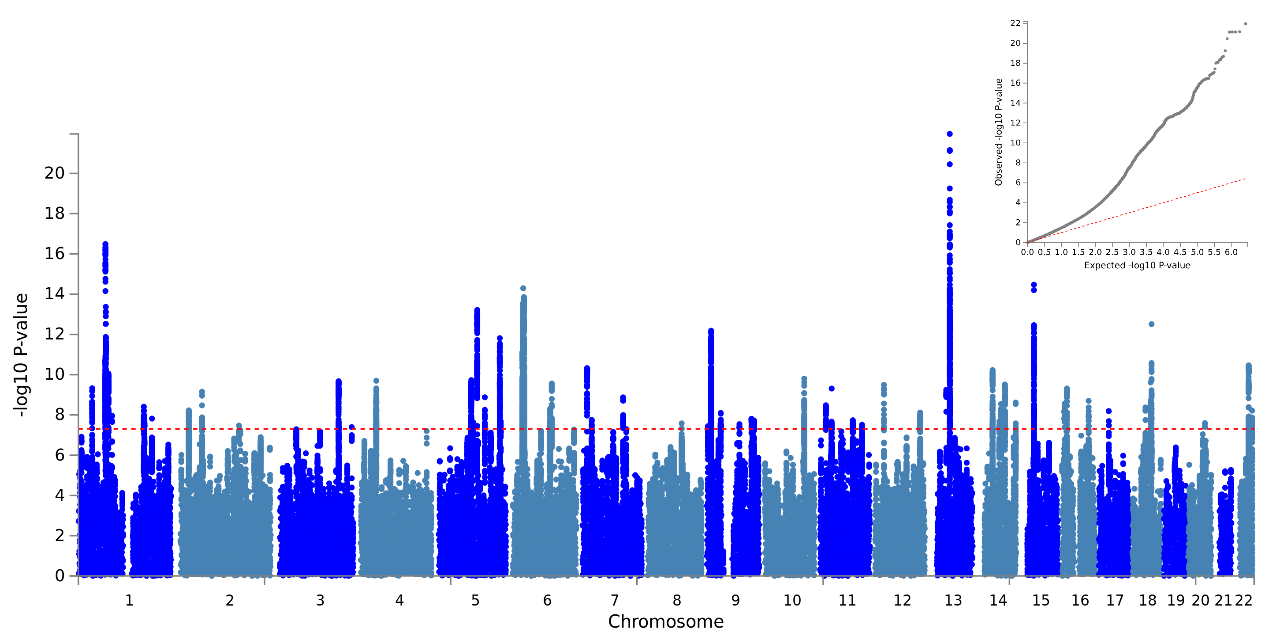

Supplementary Figure 2: Manhattan and QQ plot of results for the combined MDD meta-analysis. Red line is 5x10^-8^

#### Supplementary Figure 3

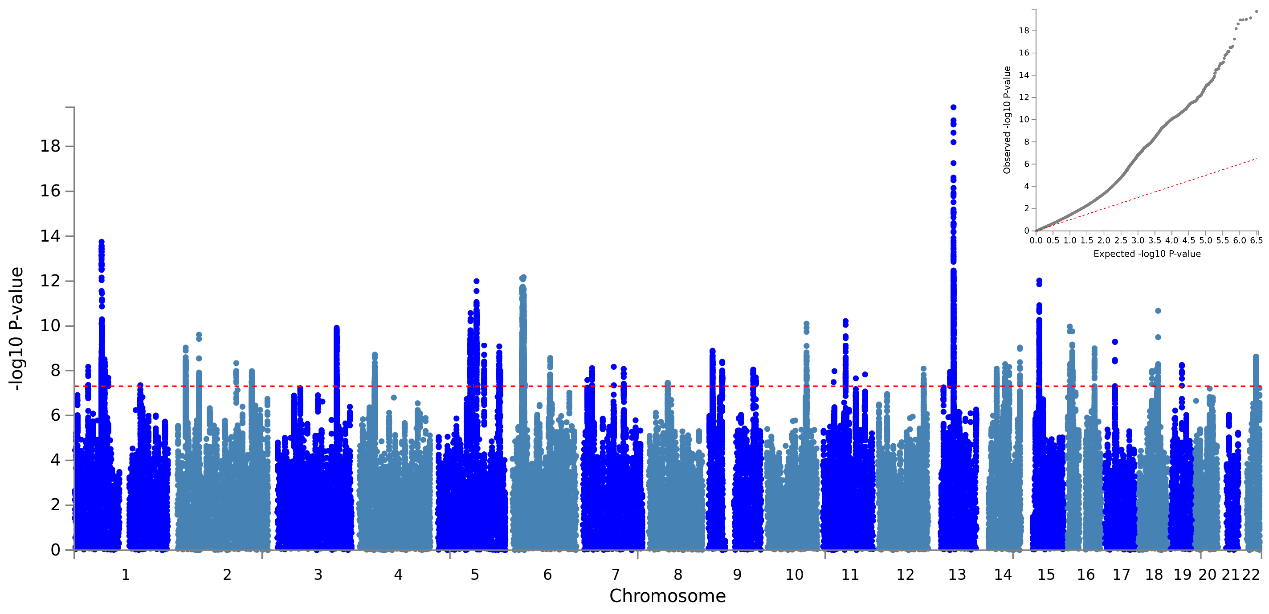

Supplementary Figure 3: Manhattan and QQ plot of results for the MDD-BD1 meta-analysis. Red line is 5x10^-8^

#### Supplementary Figure 4

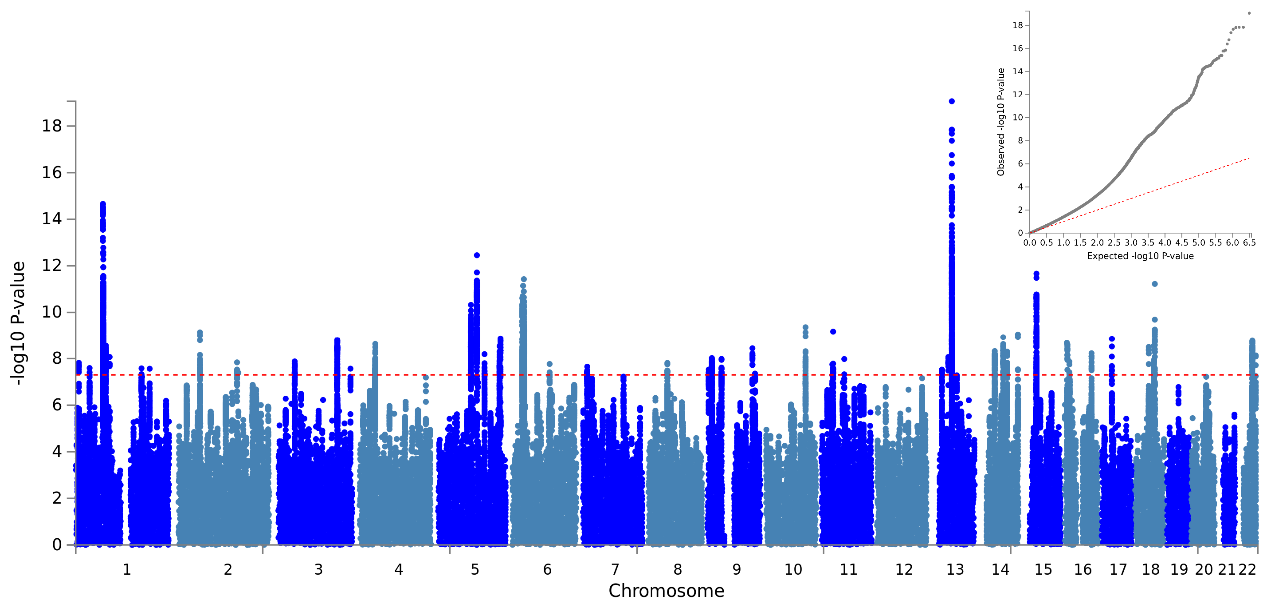

Supplementary Figure 4: Manhattan and QQ plot of results for the MDD-BD2 meta-analysis. Red line is 5x10^-8^

#### Supplementary Figure 5

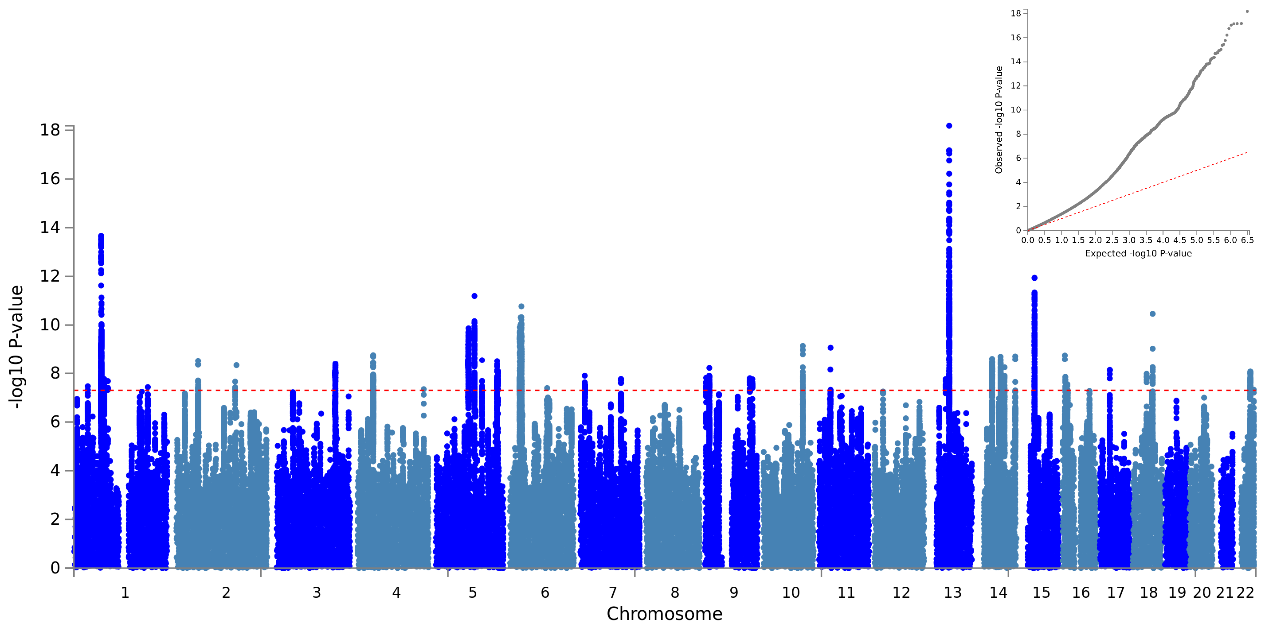

Supplementary Figure 5: Manhattan and QQ plot of results for the MDD-SAB meta-analysis. Red line is 5x10^-8^

#### Supplementary Figure 6

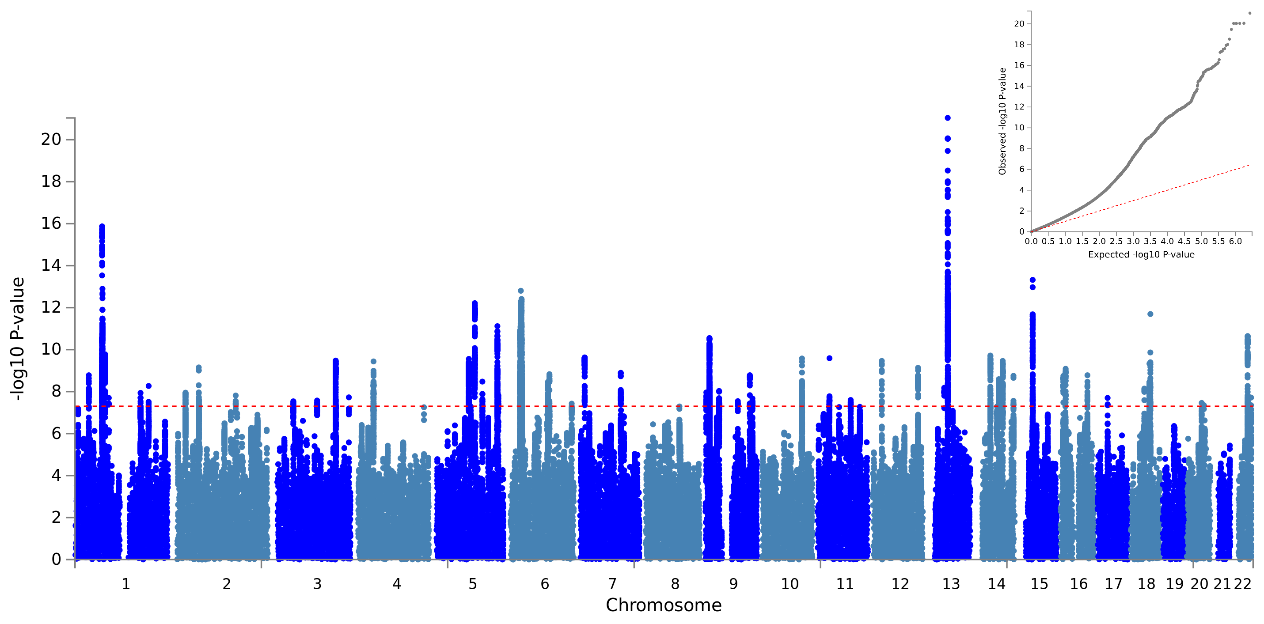

Supplementary Figure 6: Manhattan and QQ plot of results for the MDD-rMDD meta-analysis. Red line is 5x10^-8^

#### Supplementary Figure 7

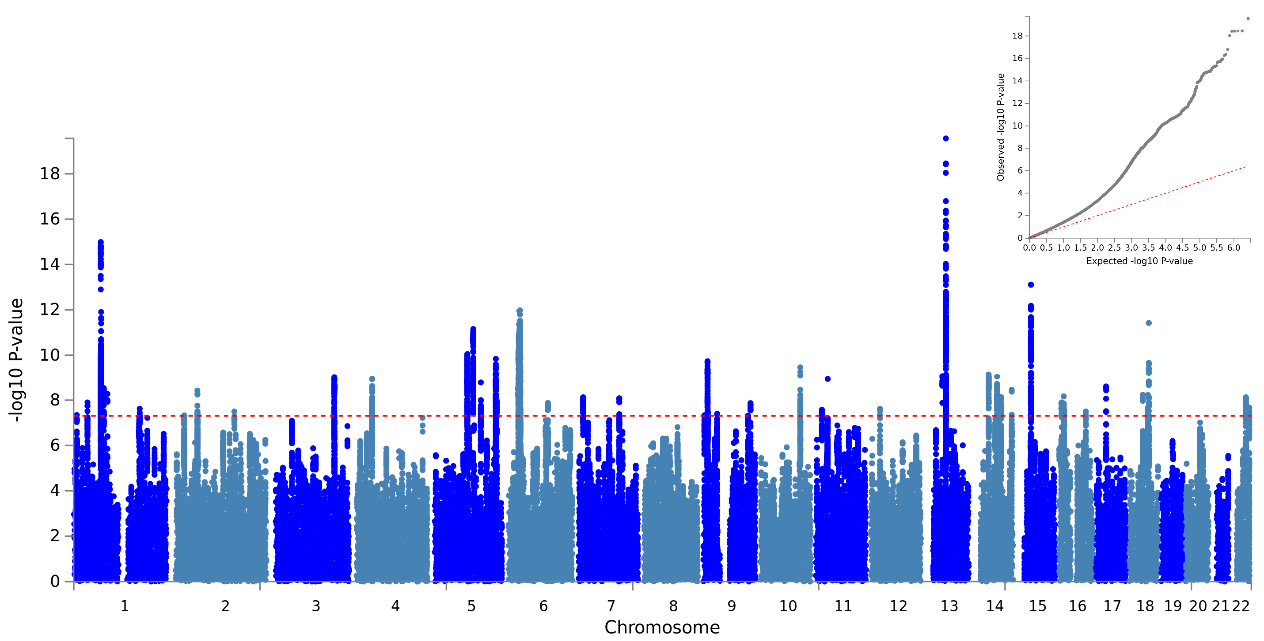

Supplementary Figure 7: Manhattan and QQ plot of results for the MDD-sMDD meta-analysis. Red line is 5x10^-8^

#### Supplementary Figure 8

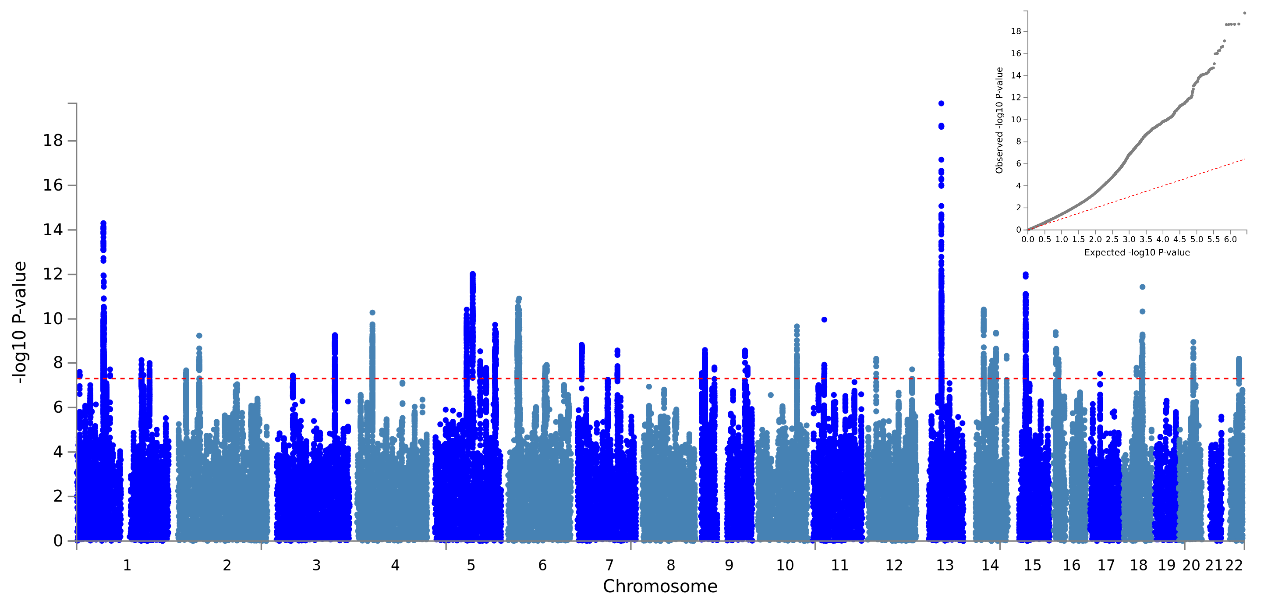

Supplementary Figure 8: Manhattan and QQ plot of results for the MDD-subMDD meta-analysis. Red line is 5x10^-8^

#### Supplementary Figure 9

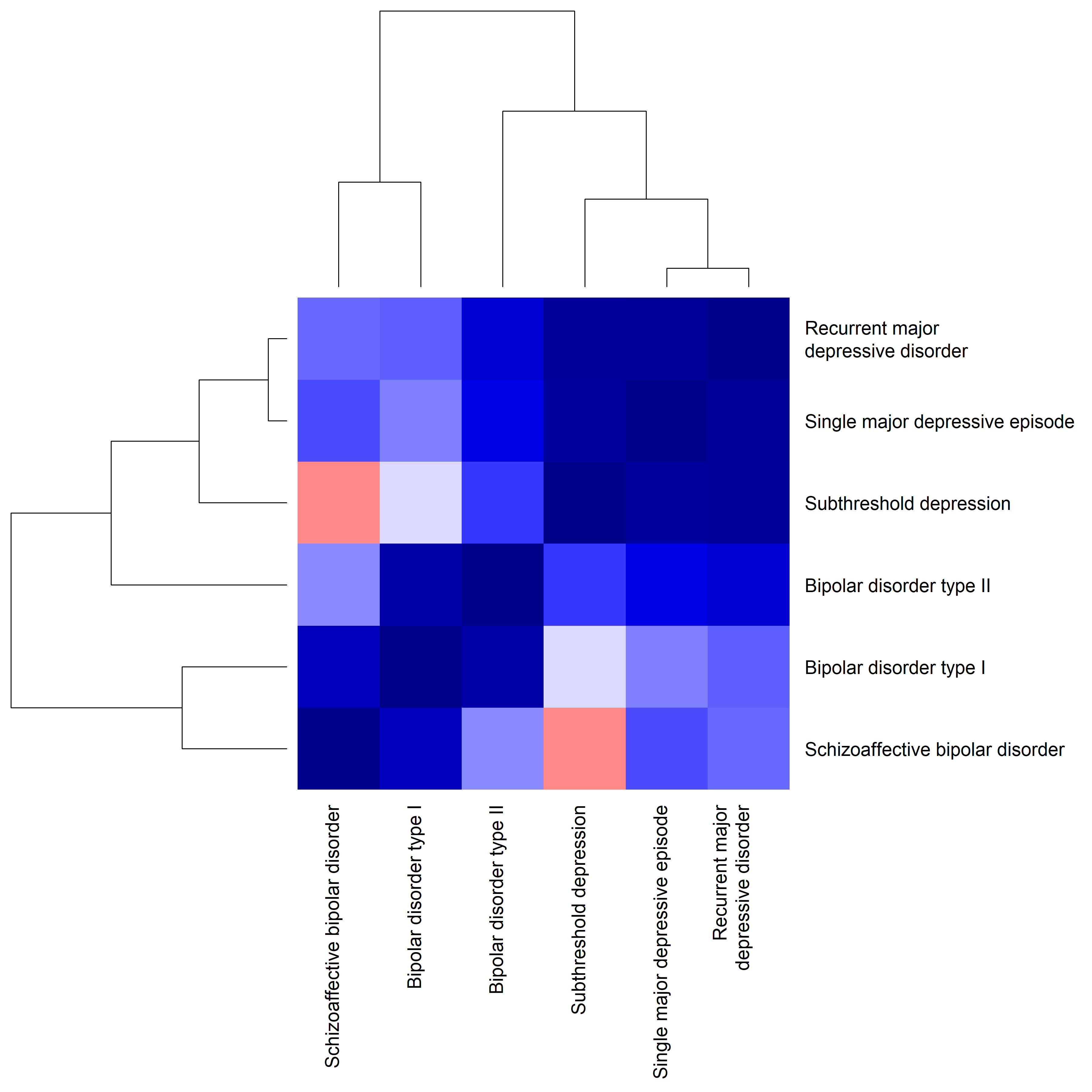

Supplementary Figure 9: Hierarchical clustering of the genetic correlations between major depression subtypes from UK Biobank (rMDD, sMDD, subMDD) and bipolar disorder subtypes (BD1, BD2, SAB). Blue = positive genetic correlation. Red = negative genetic correlation. Full genetic correlation results are provided in Supplementary Table 4.

#### Supplementary Figure 10

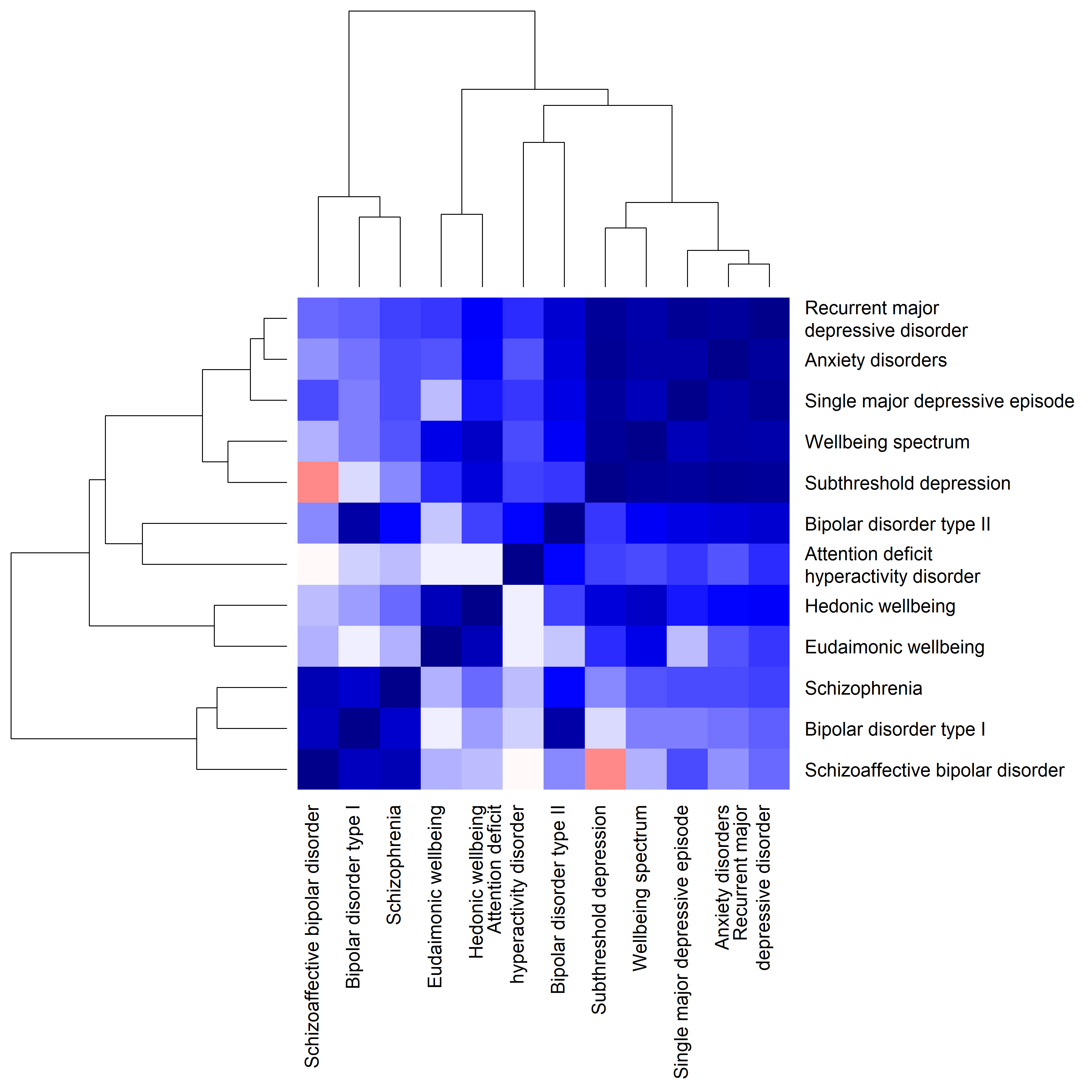

Supplementary Figure 10: Hierarchical clustering of the genetic correlations between major depression subtypes from UK Biobank (rMDD, sMDD, subMDD) and bipolar disorder subtypes (BD1, BD2, SAB), in the context of genetic correlations with external traits (schizophrenia, anxiety disorders, attention deficit hyperactivity disorder, the wellbeing spectrum, hedonic wellbeing and eudaimonic wellbeing). Genetic correlations with the wellbeing spectrum are reversed, such that they are correlations with low wellbeing.
Blue = positive genetic correlation. Red = negative genetic correlation. Full genetic correlation results are provided in Supplementary Table 4 and Supplementary Table 7 (for external traits).

#### Supplementary Figure 11

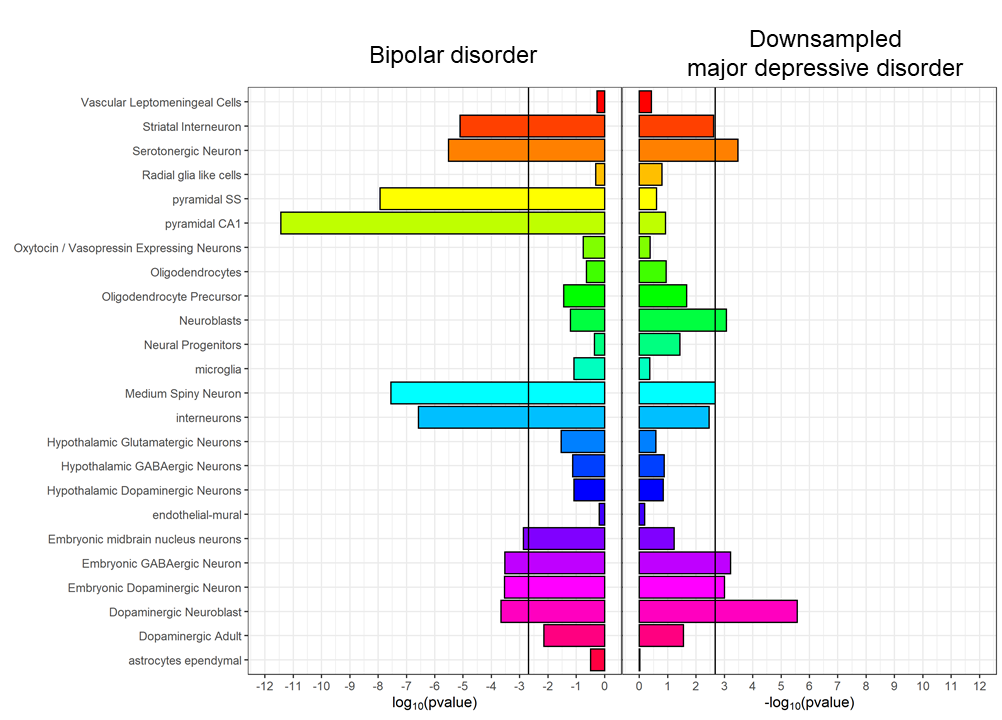

Supplementary Figure 11: Cell-type expression specificity of genes associated with bipolar disorder (PGC BD, left) and the down-sampled major depressive disorder GWAS (down-sampled MDD, right). Black vertical lines = significant enrichment (p < 2x10-3, Bonferroni correction for 24 cell types). See Supplementary Table 15 for full results.

#### Supplementary Figure 12

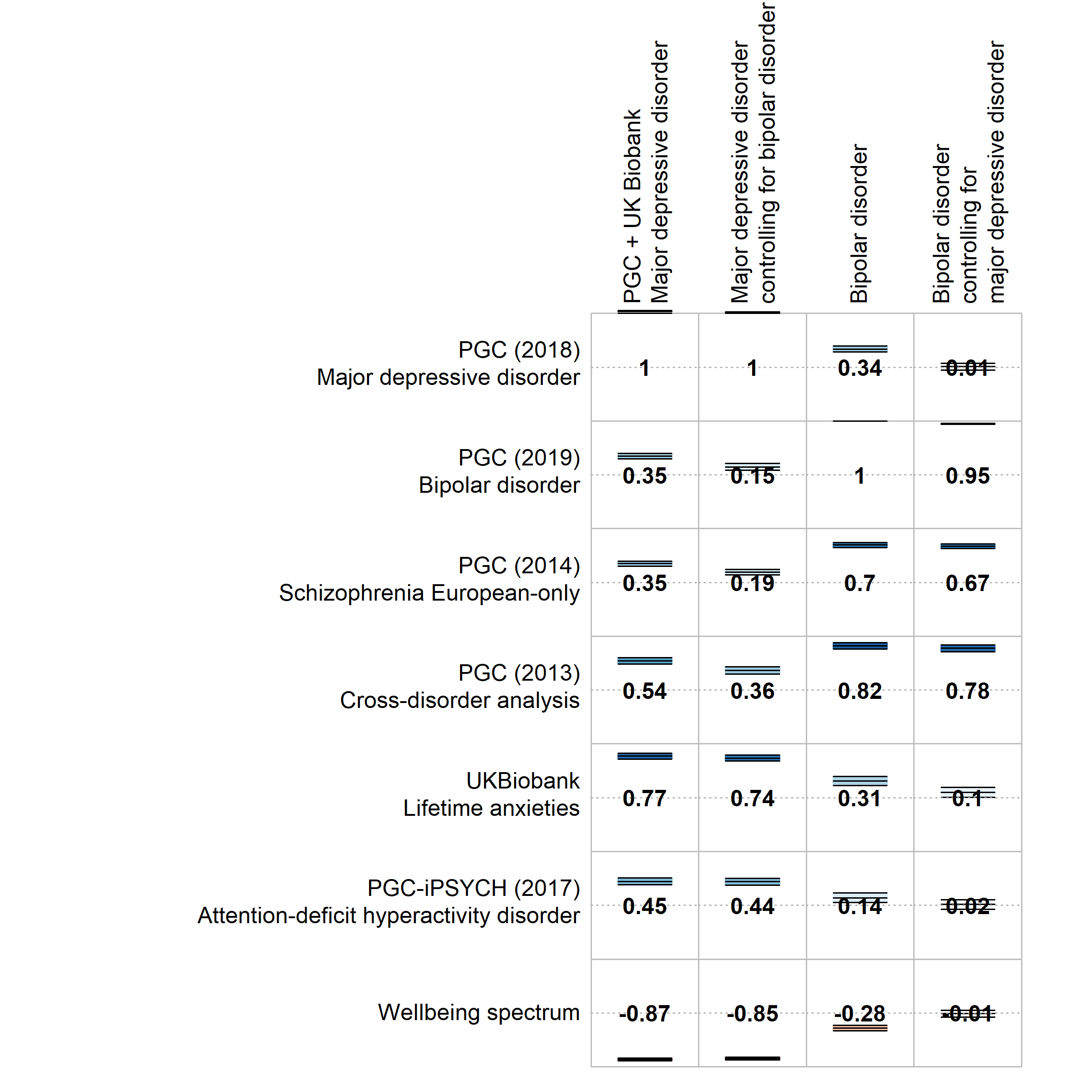

Supplementary Figure 12a: Selected genetic correlations of psychiatric traits with the main and conditional analyses of MDD (combined MDD, MDDcBD), and bipolar disorder (PGC BD and BDcMDD). Full genetic correlation results are provided in Supplementary Table 5.

*Supplementary Figure 12 (continued)*

**b)**

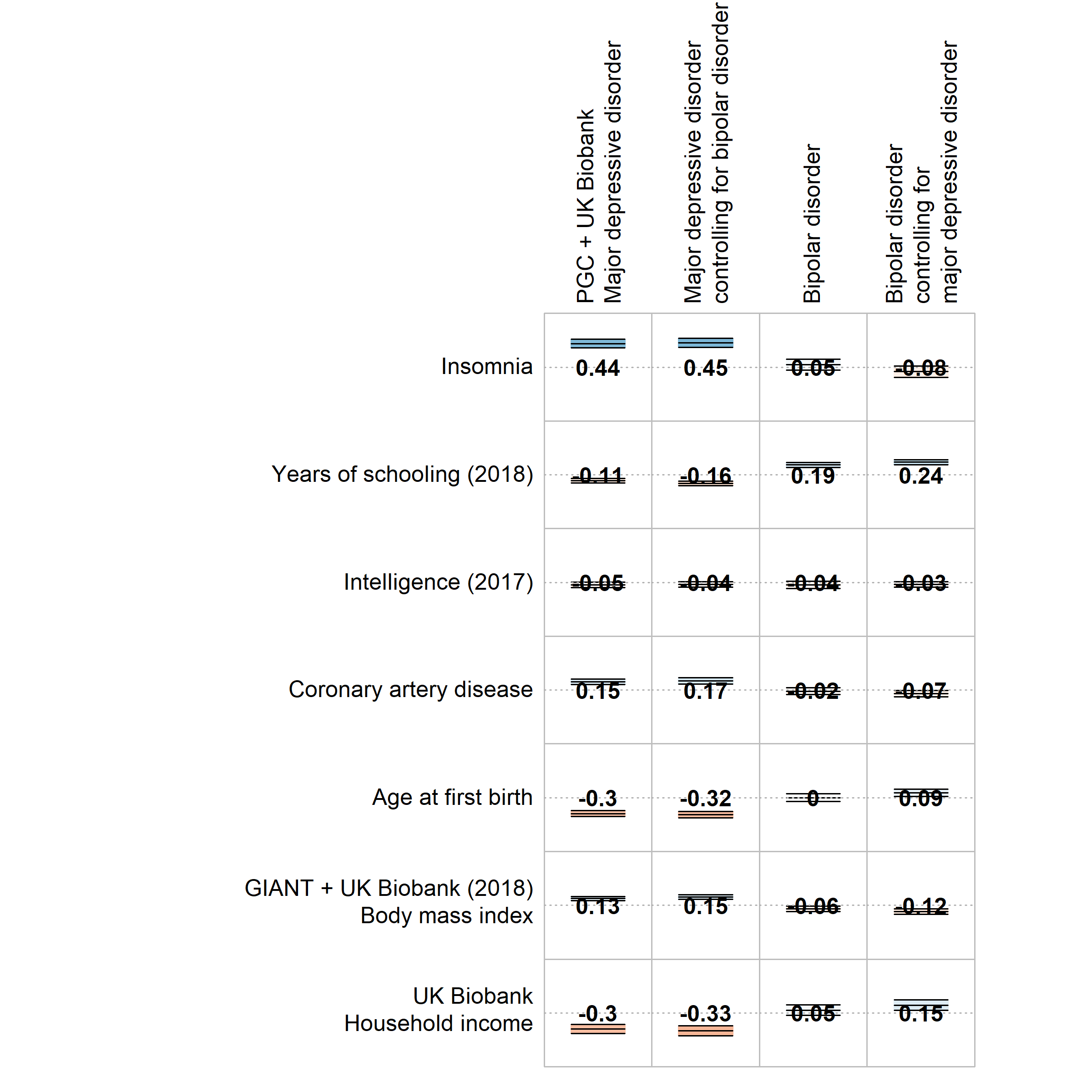

Supplementary Figure 12b: Selected genetic correlations of other traits with the main and conditional analyses of MDD (combined MDD, MDDcBD), and bipolar disorder (PGC BD and BDcMDD). Full genetic correlation results are provided in Supplementary Table 5.

#### Supplementary Figure 13

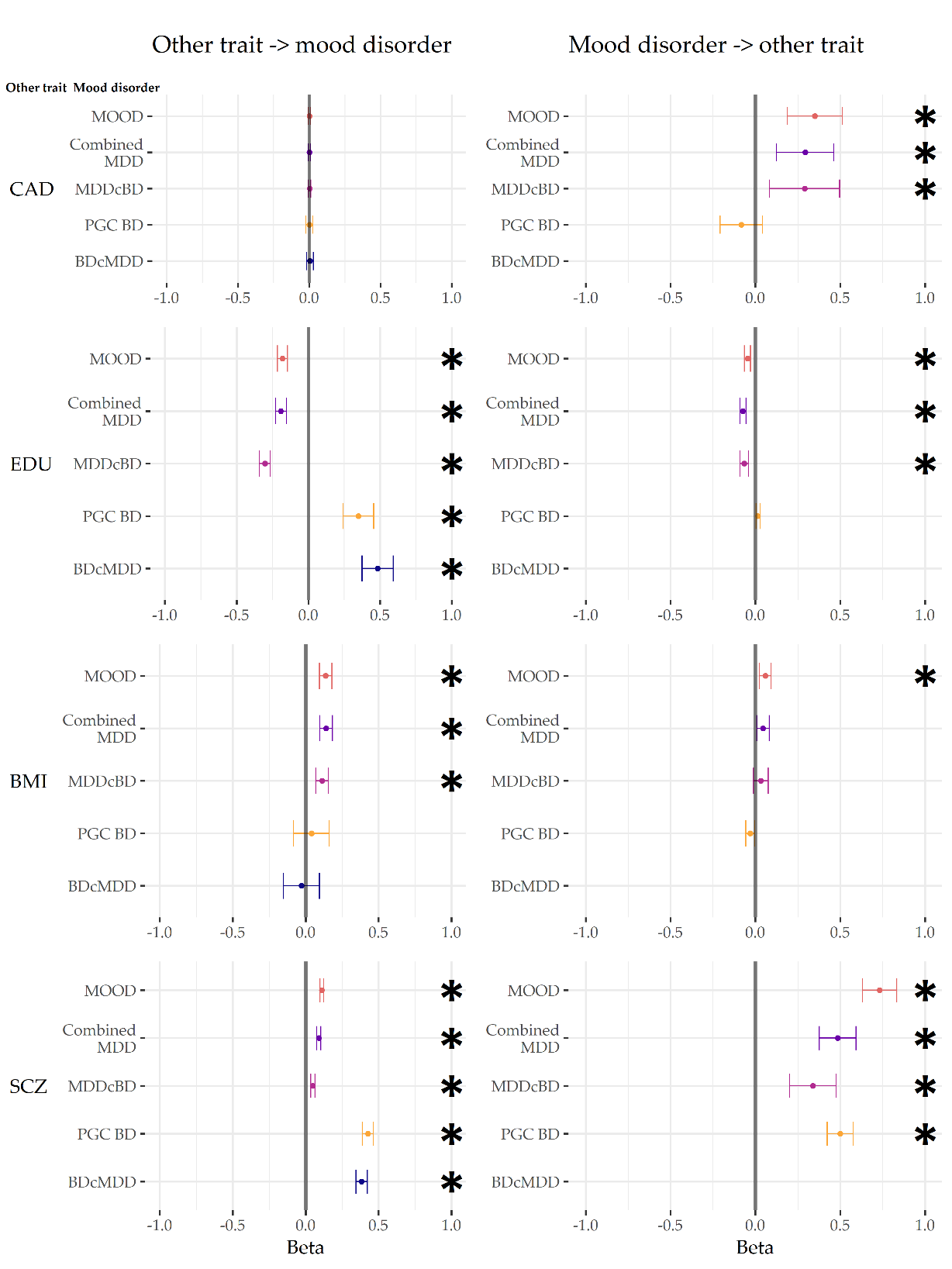

Supplementary Figure 13: GSMR results from analyses with the main meta-analysis (MOOD), the main and conditional MDD (combined MDD, MDDcBD) and bipolar disorder (PGC BD, BDcMDD) analyses. External traits are coronary artery disease (CAD), educational attainment (EDU), body mass index (BMI), and schizophrenia (SCZ).
* p < 0.004 (Bonferroni correction for two-way comparisons with six external traits). For figure data, including the number of non-pleiotropic SNPs included in each instrument, see Supplementary Table 12.

#### Supplementary Figure 14

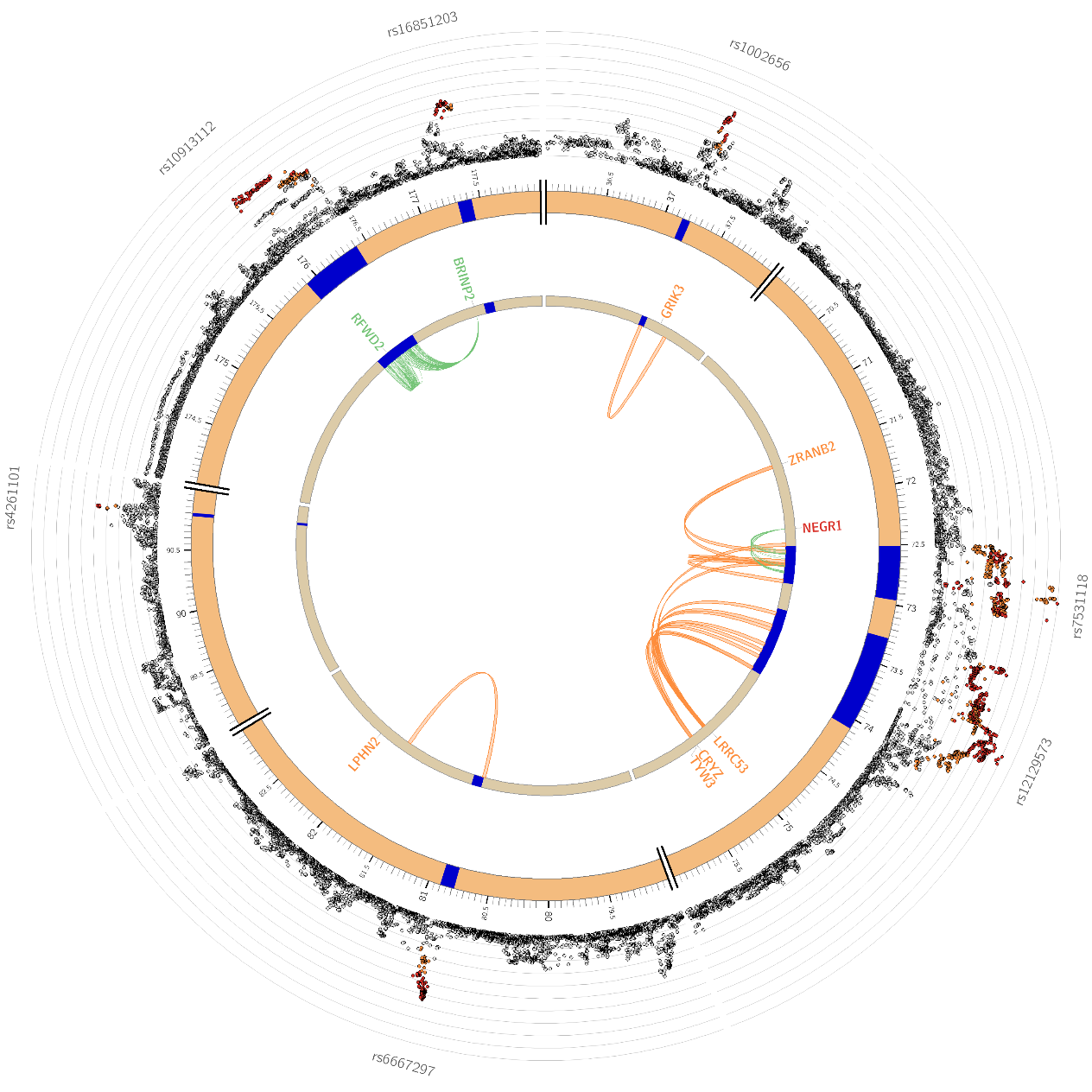

Supplementary Figure 14: Circos plot of significant loci from mood disorders (MOOD) on chromosome 1. Outer circle: Plot of individual variants in each locus, coloured by LD to labelled index variant (r2 0.8 -> 1, orange -> red). Middle ring: position of locus on chromosome, loci shown in blue. Inner ring: Links between loci and nearby genes, by eQTLs (green), chromatin contacts (orange) or both (red)

#### Supplementary Figure 15

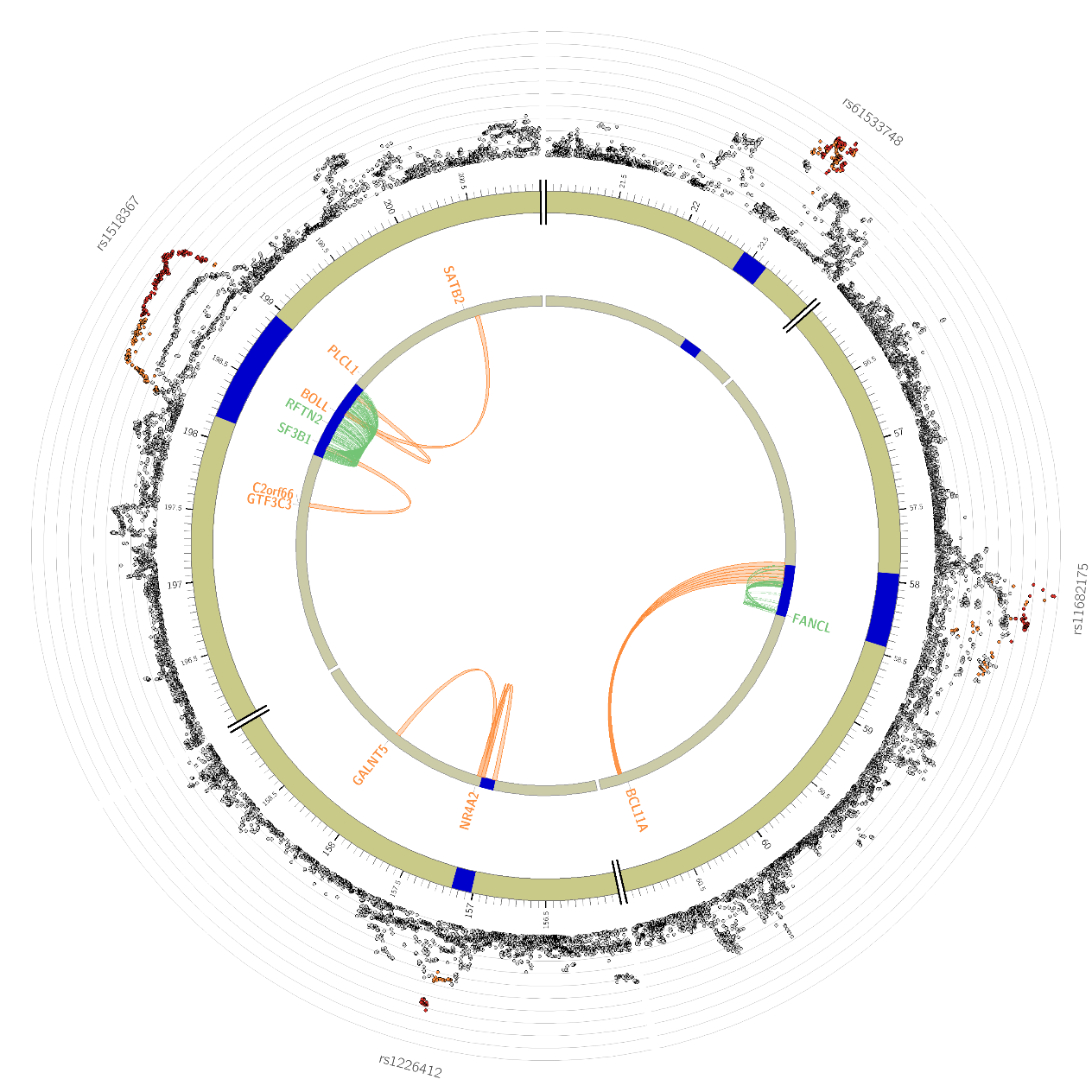

Supplementary Figure 15: Circos plot of significant loci from mood disorders (MOOD) on chromosome 2. Outer circle: Plot of individual variants in each locus, coloured by LD to labelled index variant (r2 0.8 -> 1, orange -> red). Middle ring: position of locus on chromosome, loci shown in blue. Inner ring: Links between loci and nearby genes, by eQTLs (green), chromatin contacts (orange) or both (red)

#### Supplementary Figure 16

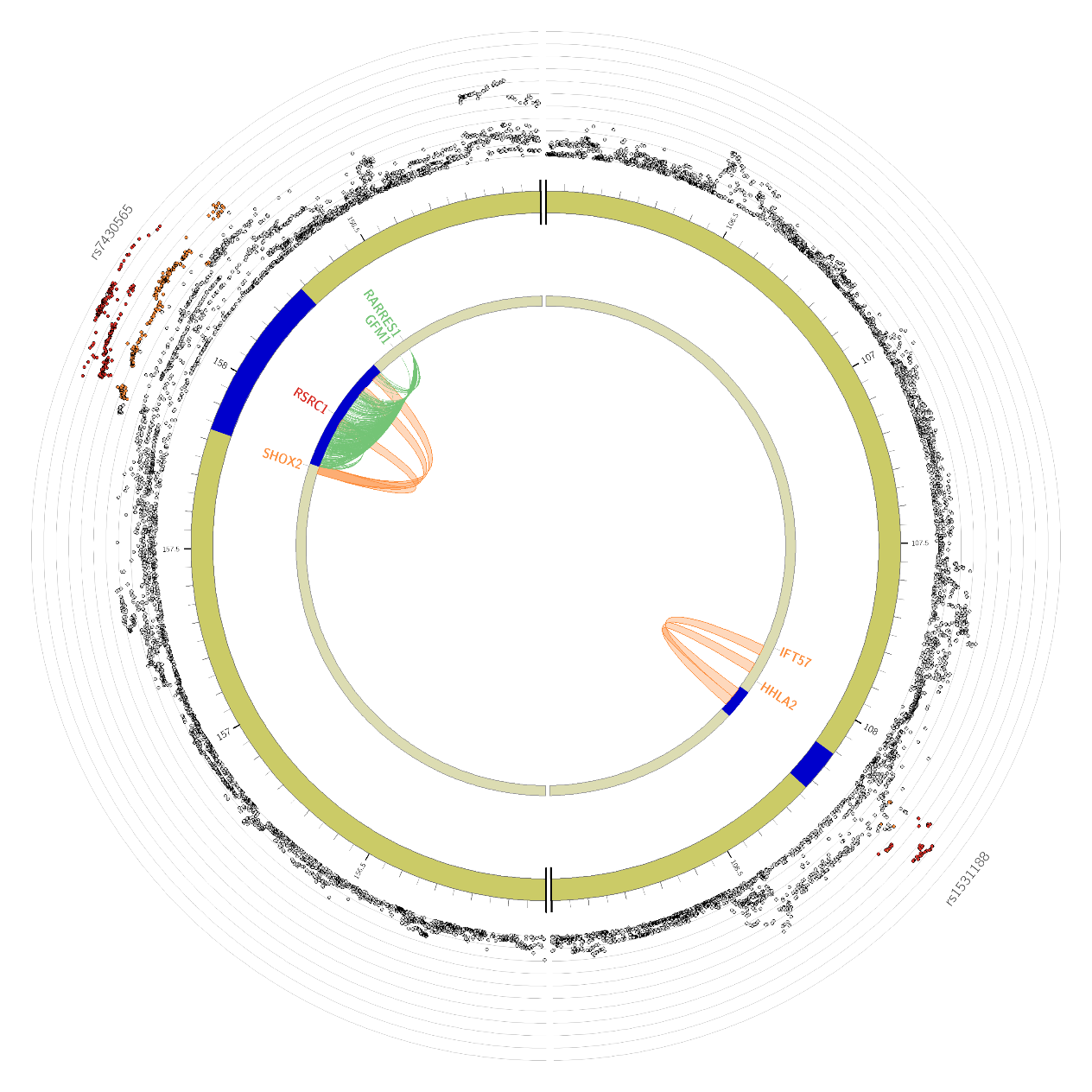

Supplementary Figure 16: Circos plot of significant loci from mood disorders (MOOD) on chromosome 3. Outer circle: Plot of individual variants in each locus, coloured by LD to labelled index variant (r2 0.8 -> 1, orange -> red). Middle ring: position of locus on chromosome, loci shown in blue. Inner ring: Links between loci and nearby genes, by eQTLs (green), chromatin contacts (orange) or both (red)

#### Supplementary Figure 17

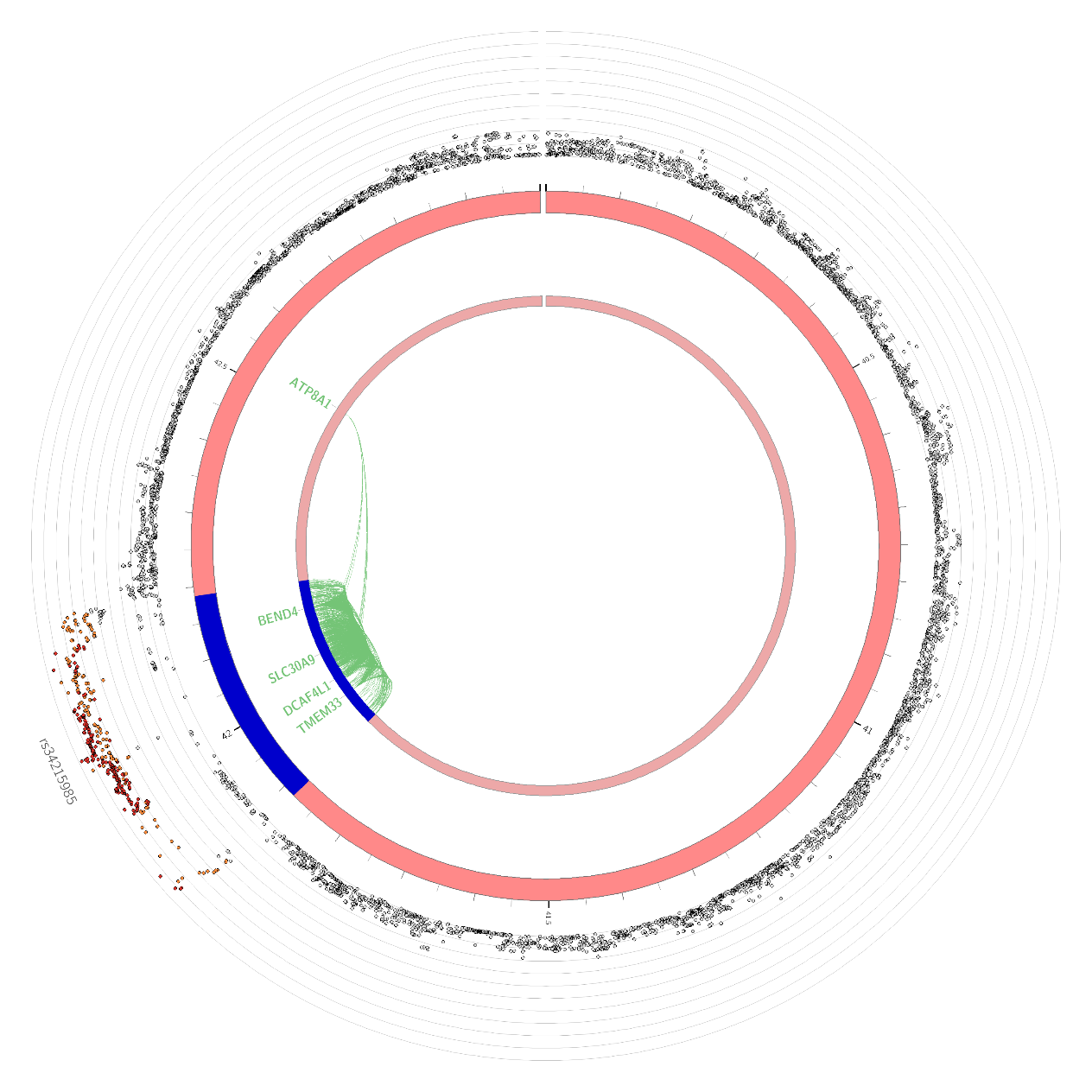

Supplementary Figure 17: Circos plot of significant loci from mood disorders (MOOD) on chromosome 4. Outer circle: Plot of individual variants in each locus, coloured by LD to labelled index variant (r2 0.8 -> 1, orange -> red). Middle ring: position of locus on chromosome, loci shown in blue. Inner ring: Links between loci and nearby genes, by eQTLs (green), chromatin contacts (orange) or both (red)

#### Supplementary Figure 18

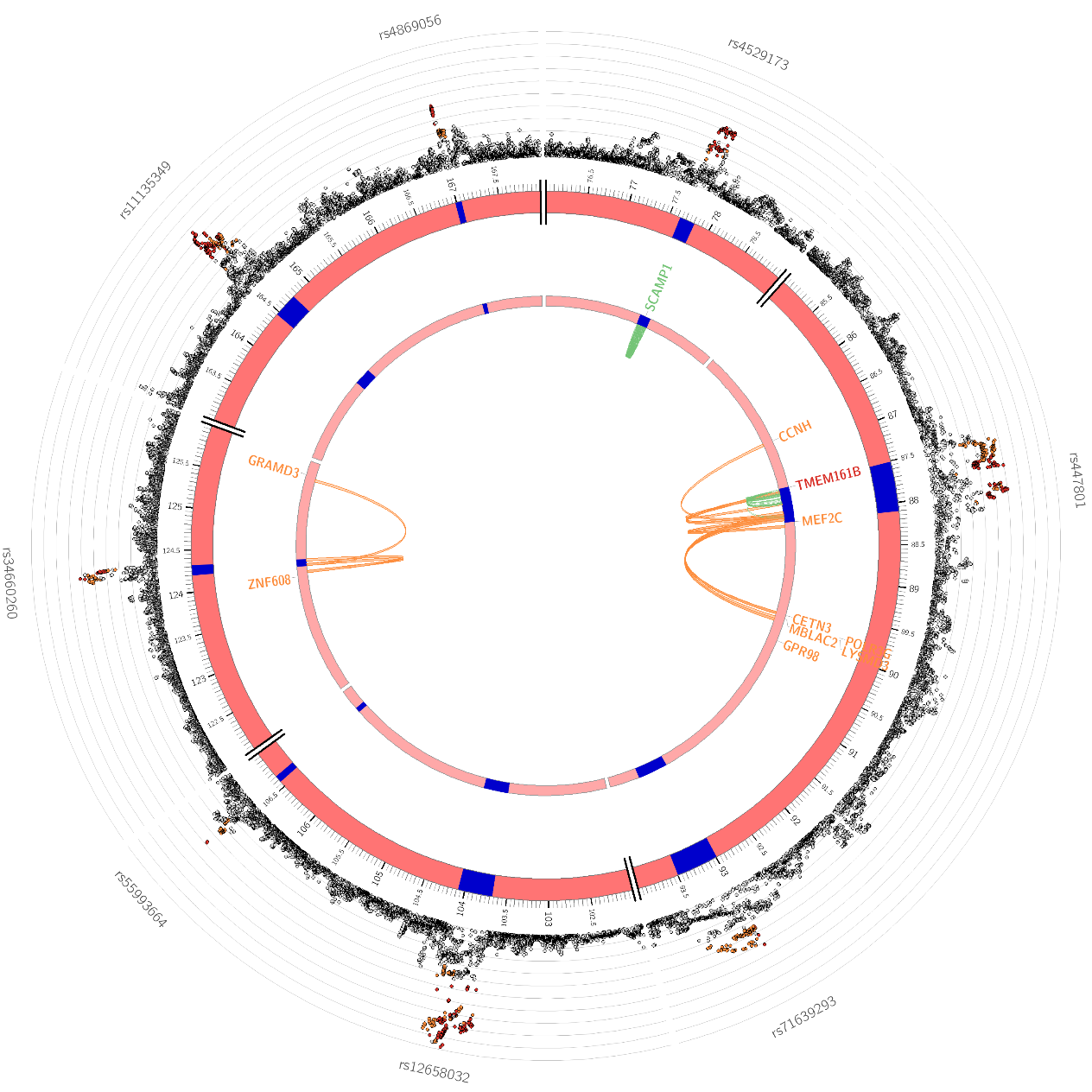

Supplementary Figure 18: Circos plot of significant loci from mood disorders (MOOD) on chromosome 5. Outer circle: Plot of individual variants in each locus, coloured by LD to labelled index variant (r2 0.8 -> 1, orange -> red). Middle ring: position of locus on chromosome, loci shown in blue. Inner ring: Links between loci and nearby genes, by eQTLs (green), chromatin contacts (orange) or both (red)

#### Supplementary Figure 19

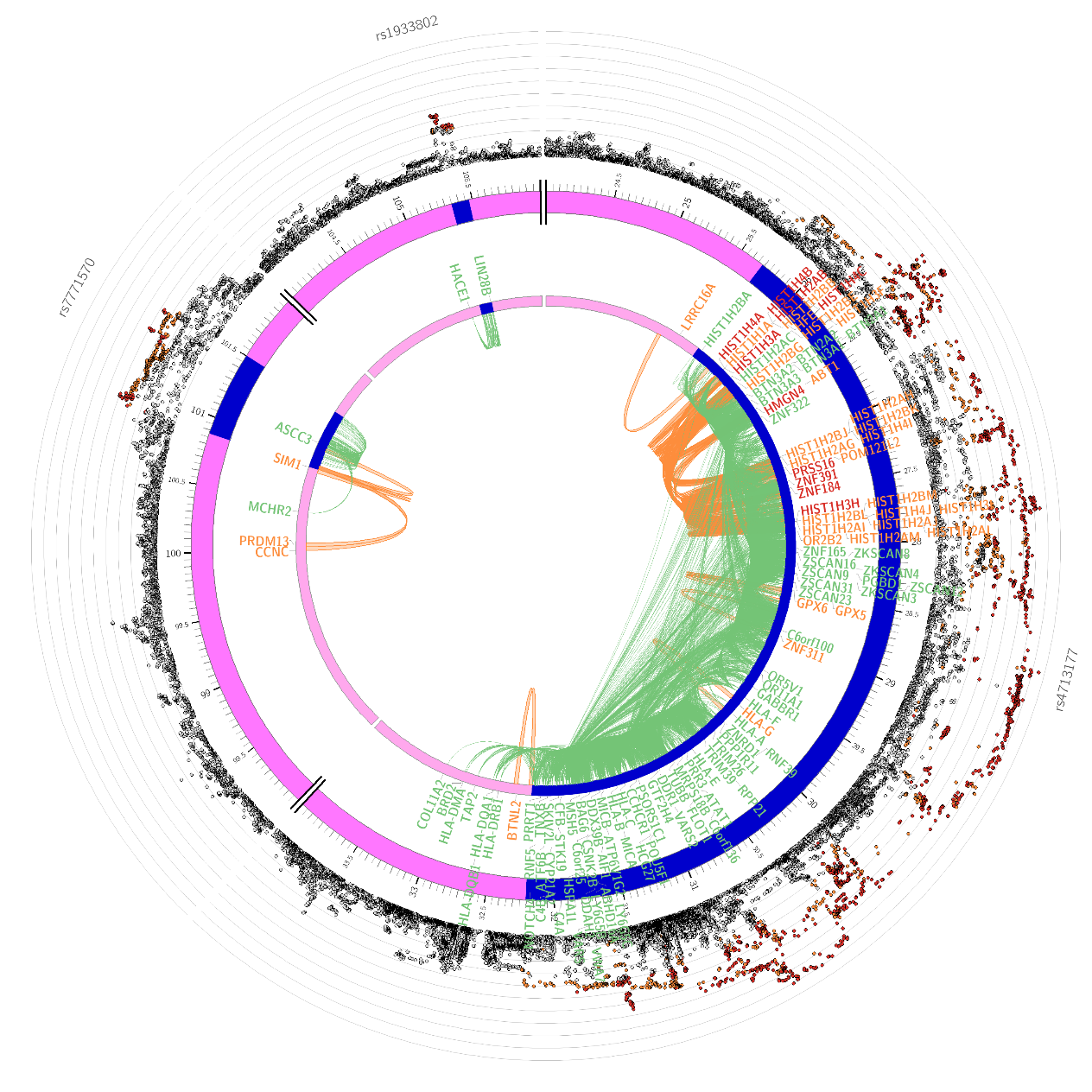

Supplementary Figure 19: Circos plot of significant loci from mood disorders (MOOD) on chromosome 6. Outer circle: Plot of individual variants in each locus, coloured by LD to labelled index variant (r2 0.8 -> 1, orange -> red). Middle ring: position of locus on chromosome, loci shown in blue. Inner ring: Links between loci and nearby genes, by eQTLs (green), chromatin contacts (orange) or both (red)

#### Supplementary Figure 20

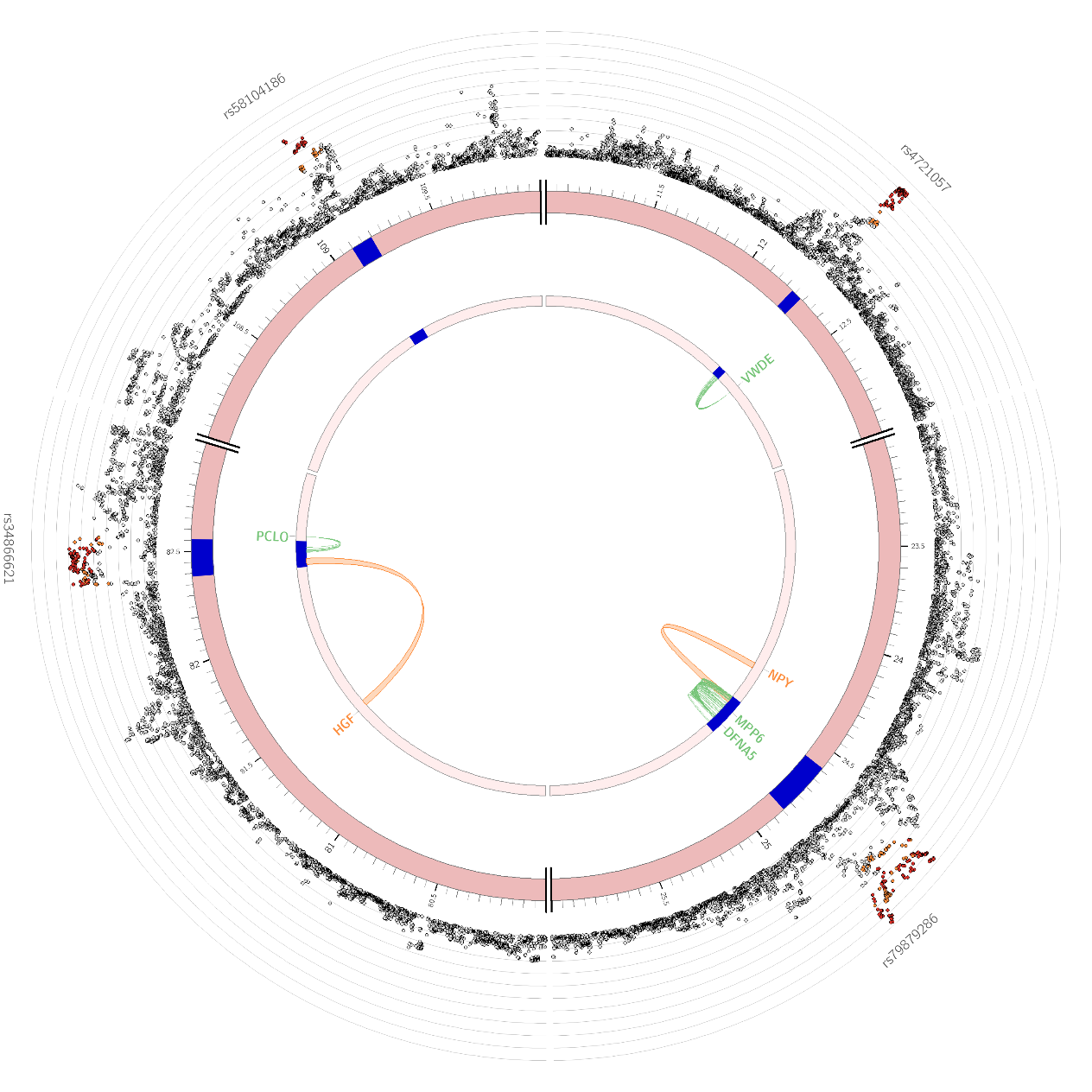

Supplementary Figure 20: Circos plot of significant loci from mood disorders (MOOD) on chromosome 7. Outer circle: Plot of individual variants in each locus, coloured by LD to labelled index variant (r2 0.8 -> 1, orange -> red). Middle ring: position of locus on chromosome, loci shown in blue. Inner ring: Links between loci and nearby genes, by eQTLs (green), chromatin contacts (orange) or both (red)

#### Supplementary Figure 21

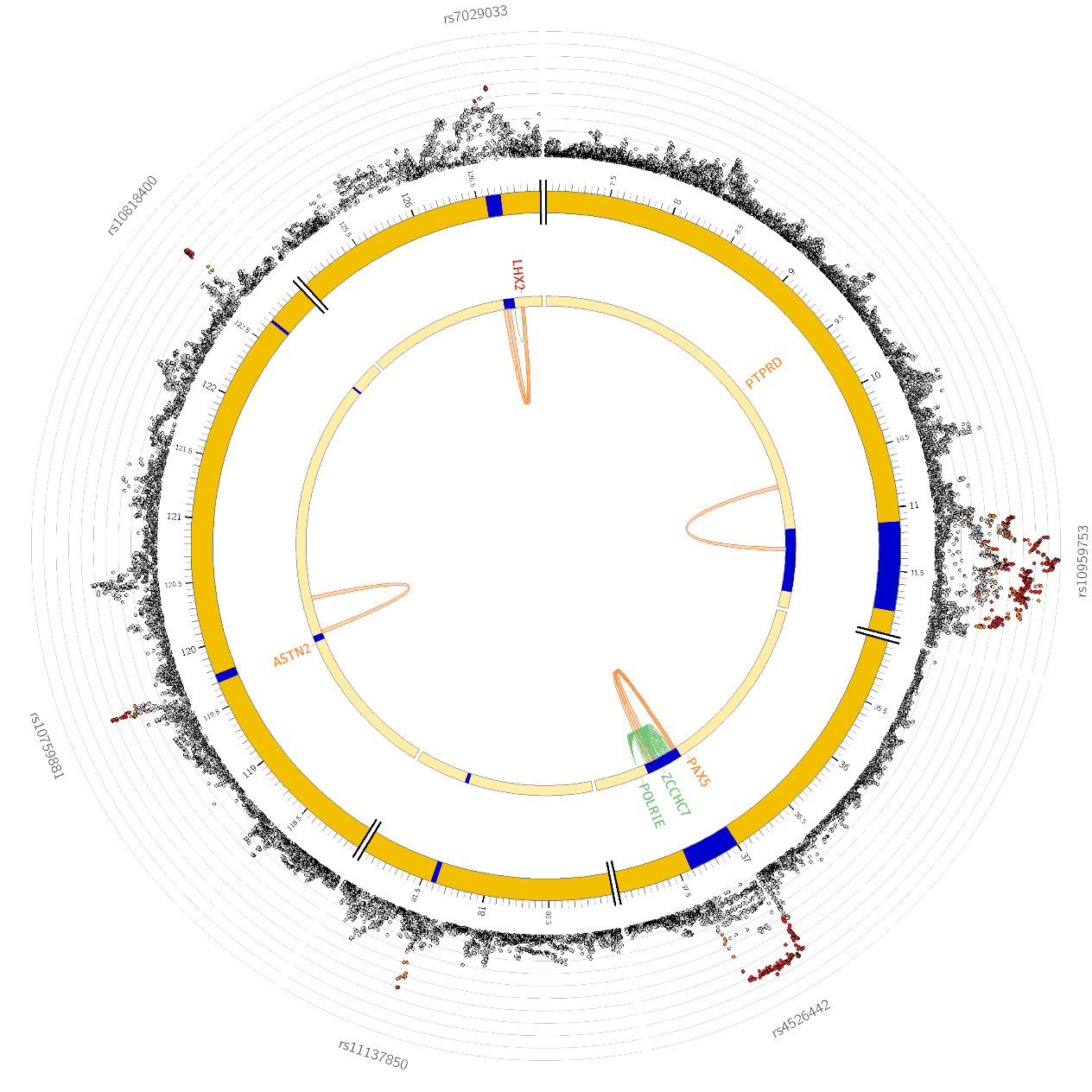

Supplementary Figure 21: Circos plot of significant loci from mood disorders (MOOD) on chromosome 9 (Note – there are no significant loci present on chromosome 8, so no circos plot is required). Outer circle: Plot of individual variants in each locus, coloured by LD to labelled index variant (r2 0.8 -> 1, orange -> red). Middle ring: position of locus on chromosome, loci shown in blue. Inner ring: Links between loci and nearby genes, by eQTLs (green), chromatin contacts (orange) or both (red)

#### Supplementary Figure 22

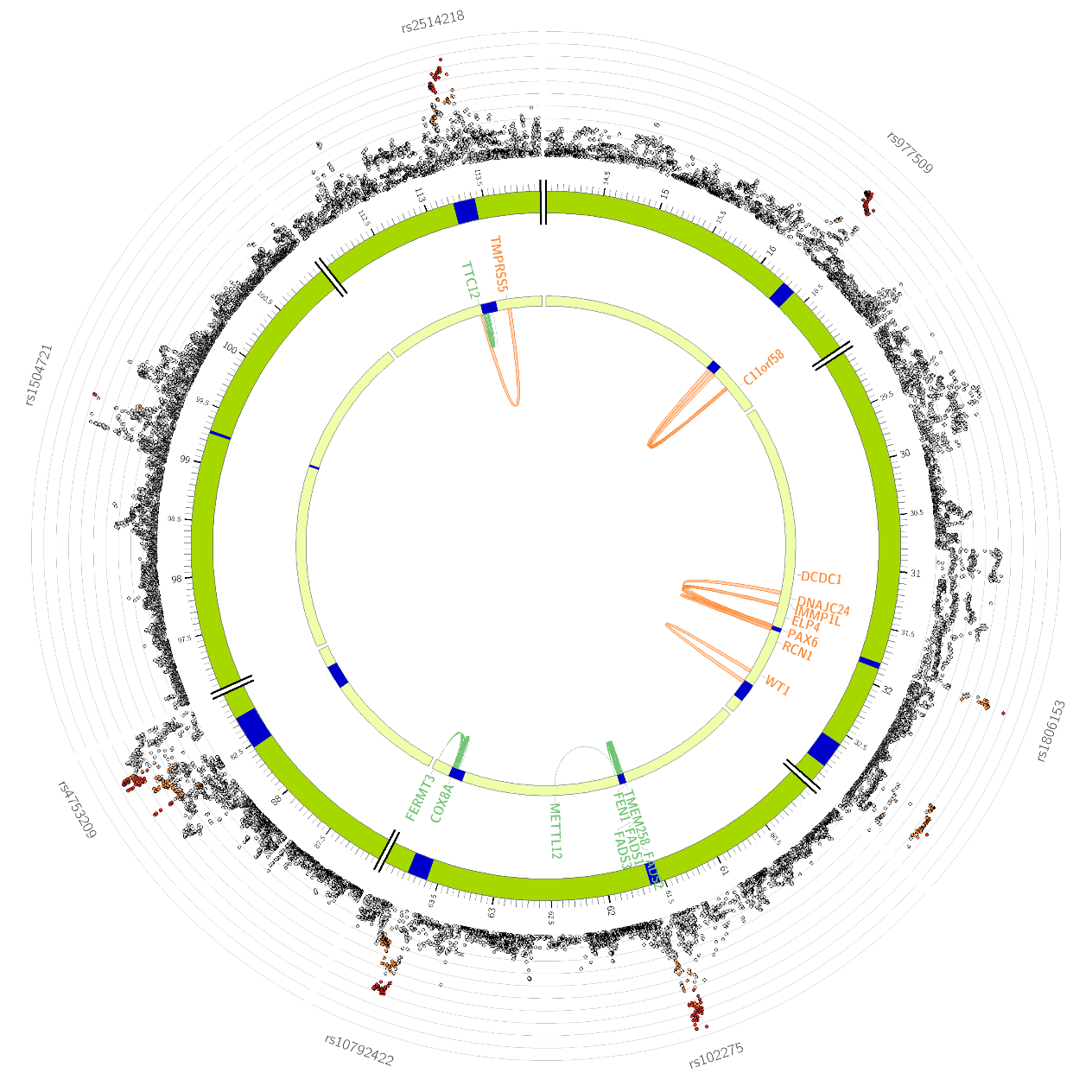

Supplementary Figure 22: Circos plot of significant loci from mood disorders (MOOD) on chromosome 10. Outer circle: Plot of individual variants in each locus, coloured by LD to labelled index variant (r2 0.8 -> 1, orange -> red). Middle ring: position of locus on chromosome, loci shown in blue. Inner ring: Links between loci and nearby genes, by eQTLs (green), chromatin contacts (orange) or both (red)

#### Supplementary Figure 23

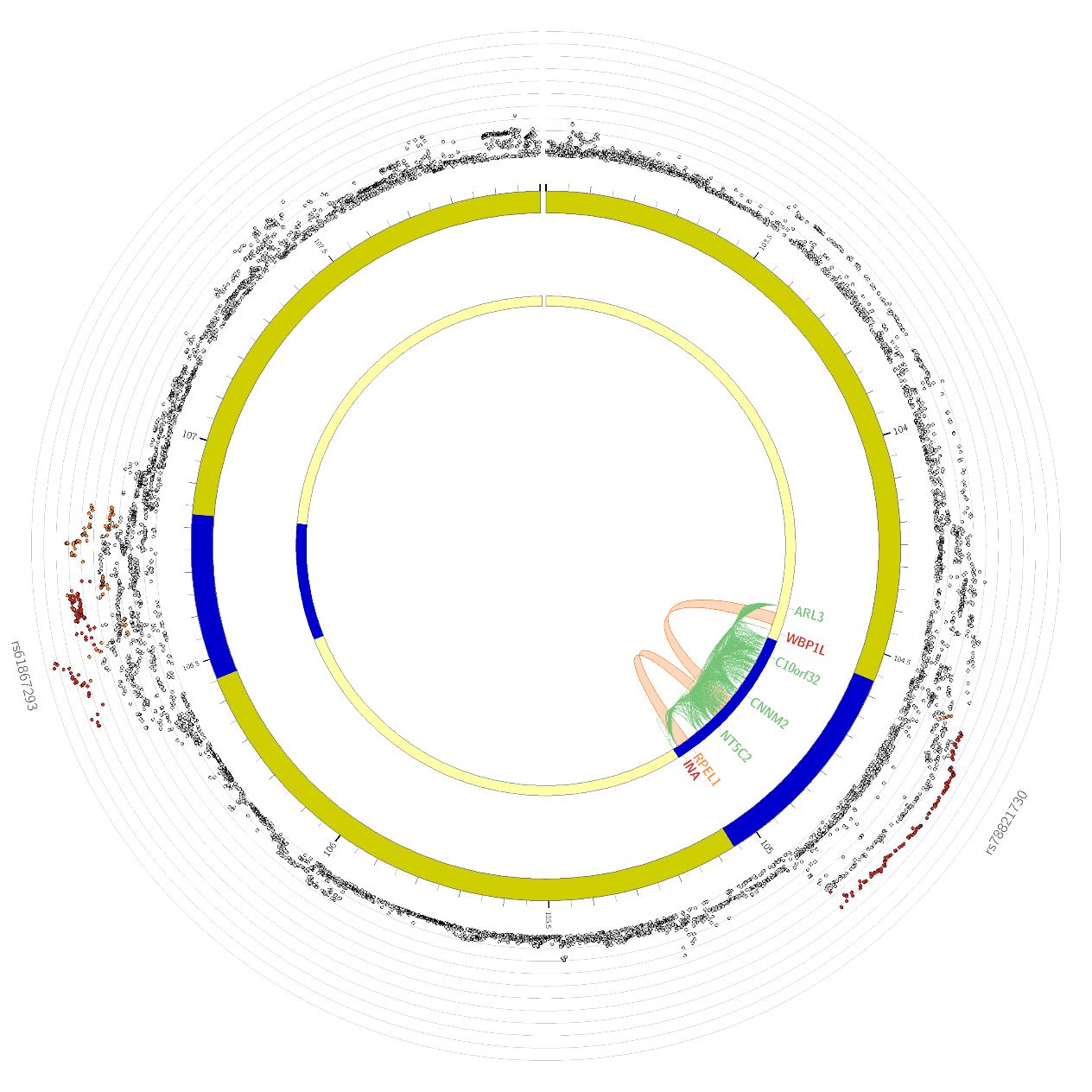

Supplementary Figure 23: Circos plot of significant loci from mood disorders (MOOD) on chromosome 11. Outer circle: Plot of individual variants in each locus, coloured by LD to labelled index variant (r2 0.8 -> 1, orange -> red). Middle ring: position of locus on chromosome, loci shown in blue. Inner ring: Links between loci and nearby genes, by eQTLs (green), chromatin contacts (orange) or both (red)

#### Supplementary Figure 24

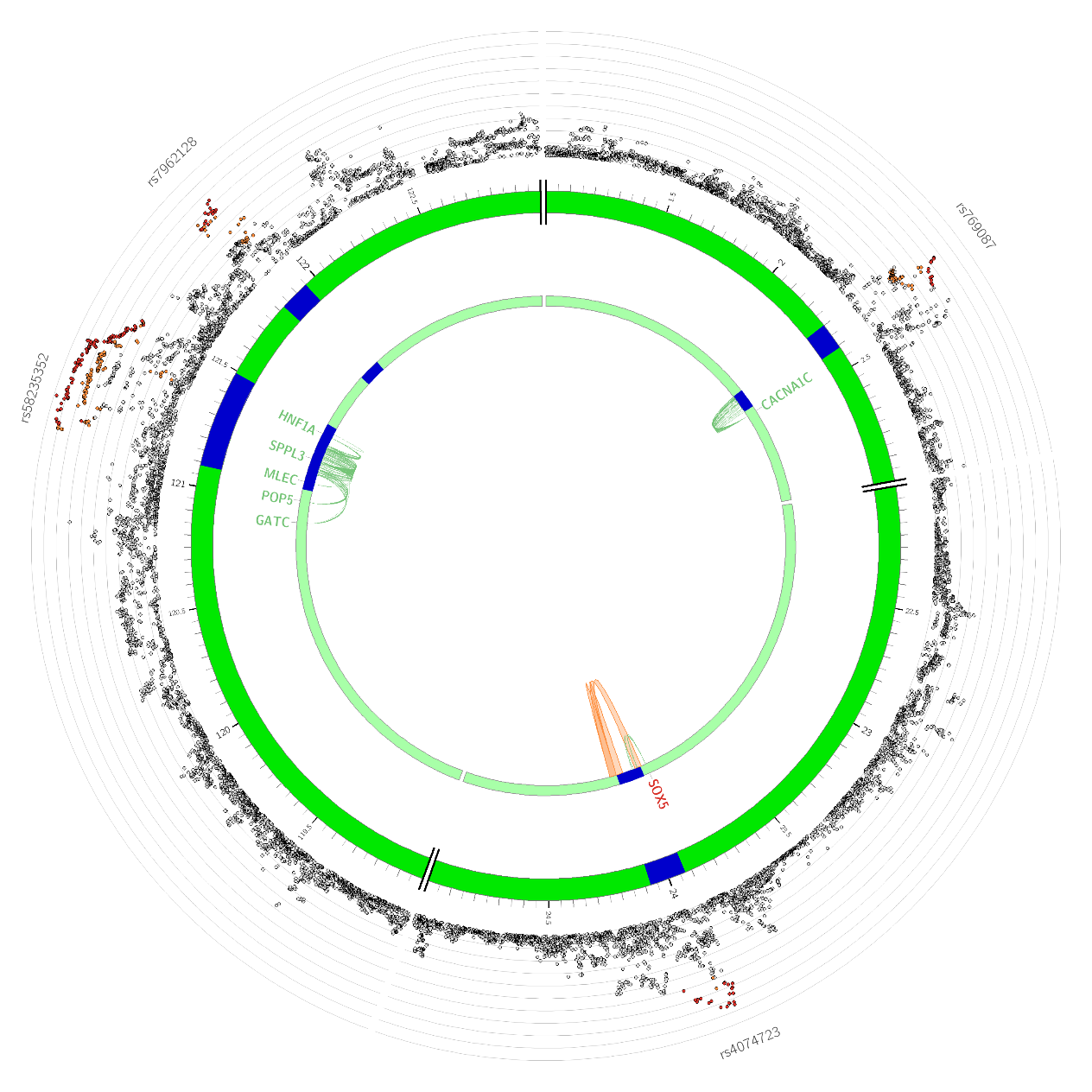

Supplementary Figure 24: Circos plot of significant loci from mood disorders (MOOD) on chromosome 12. Outer circle: Plot of individual variants in each locus, coloured by LD to labelled index variant (r2 0.8 -> 1, orange -> red). Middle ring: position of locus on chromosome, loci shown in blue. Inner ring: Links between loci and nearby genes, by eQTLs (green), chromatin contacts (orange) or both (red)

#### Supplementary Figure 25

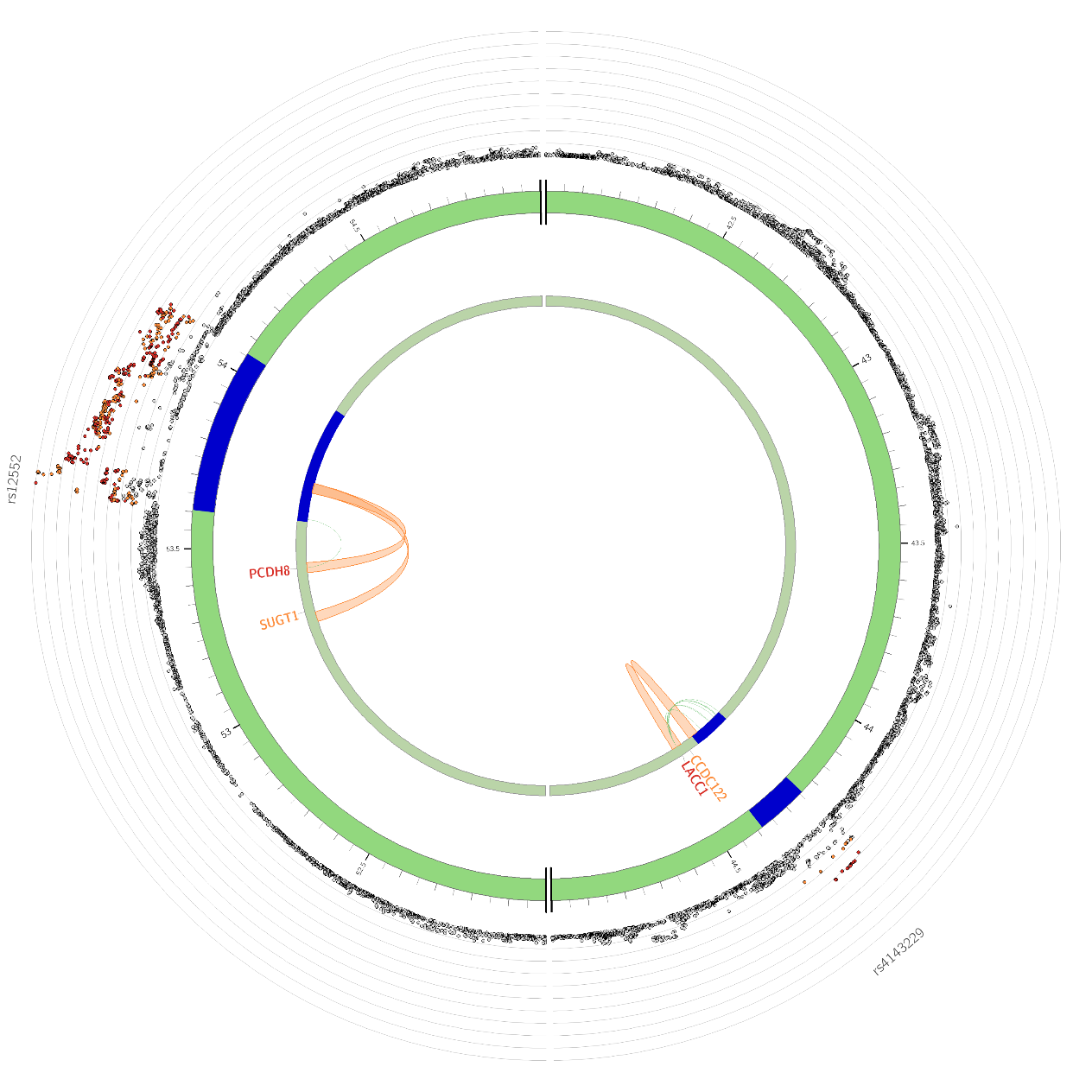

Supplementary Figure 25: Circos plot of significant loci from mood disorders (MOOD) on chromosome 13. Outer circle: Plot of individual variants in each locus, coloured by LD to labelled index variant (r2 0.8 -> 1, orange -> red). Middle ring: position of locus on chromosome, loci shown in blue. Inner ring: Links between loci and nearby genes, by eQTLs (green), chromatin contacts (orange) or both (red)

#### Supplementary Figure 26

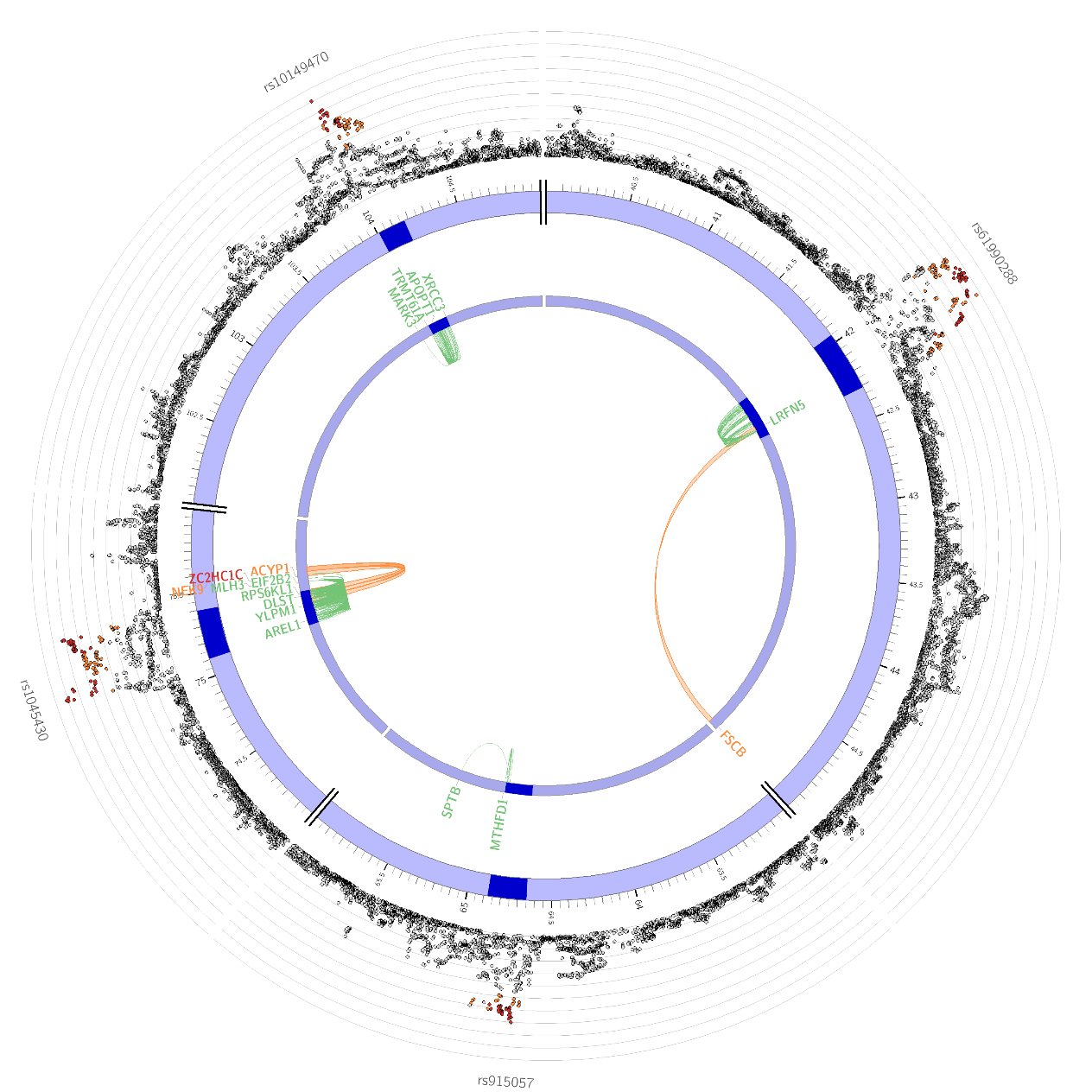

Supplementary Figure 26: Circos plot of significant loci from mood disorders (MOOD) on chromosome 14. Outer circle: Plot of individual variants in each locus, coloured by LD to labelled index variant (r2 0.8 -> 1, orange -> red). Middle ring: position of locus on chromosome, loci shown in blue. Inner ring: Links between loci and nearby genes, by eQTLs (green), chromatin contacts (orange) or both (red)

#### Supplementary Figure 27

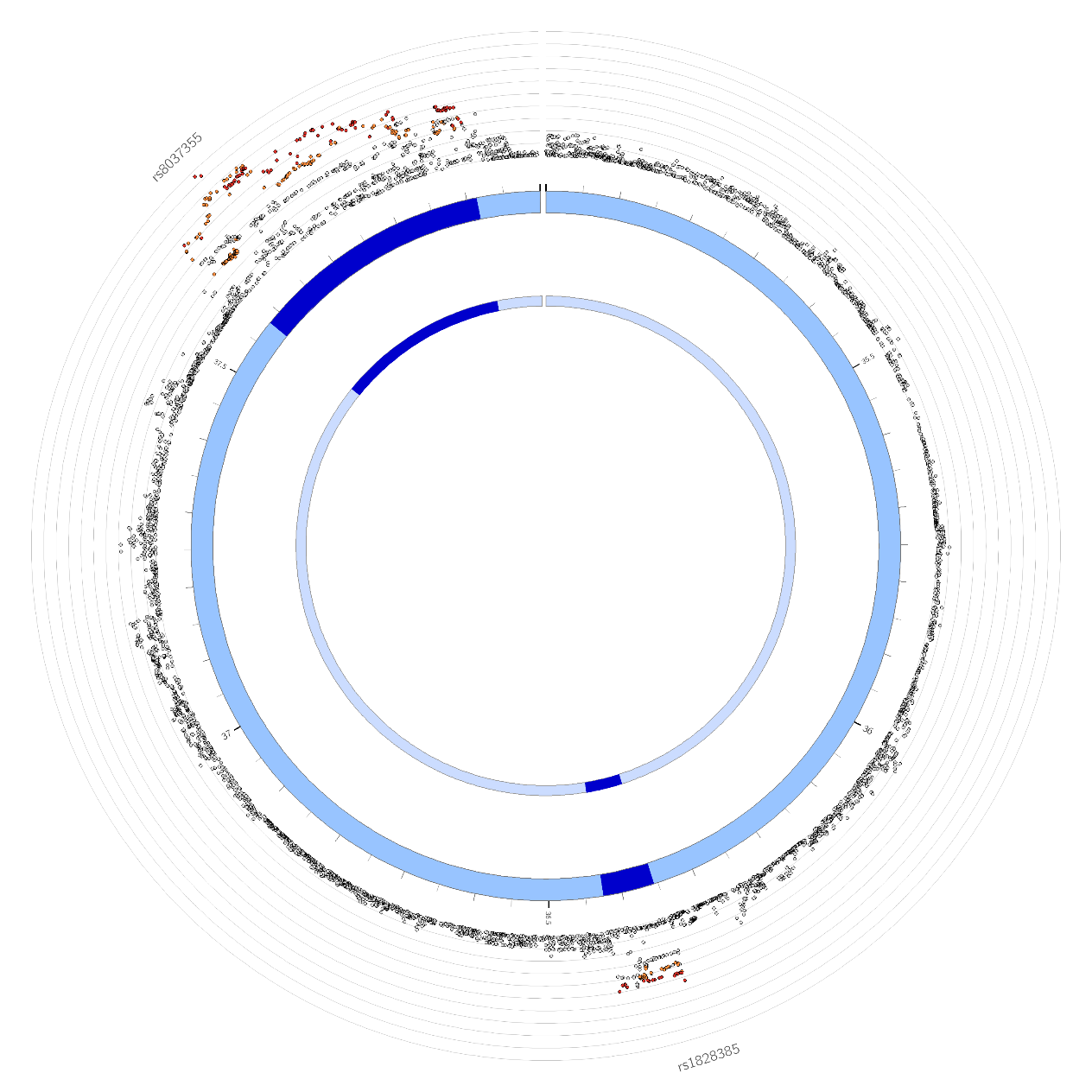

Supplementary Figure 27: Circos plot of significant loci from mood disorders (MOOD) on chromosome 15. Outer circle: Plot of individual variants in each locus, coloured by LD to labelled index variant (r2 0.8 -> 1, orange -> red). Middle ring: position of locus on chromosome, loci shown in blue. Inner ring: Links between loci and nearby genes, by eQTLs (green), chromatin contacts (orange) or both (red)

#### Supplementary Figure 28

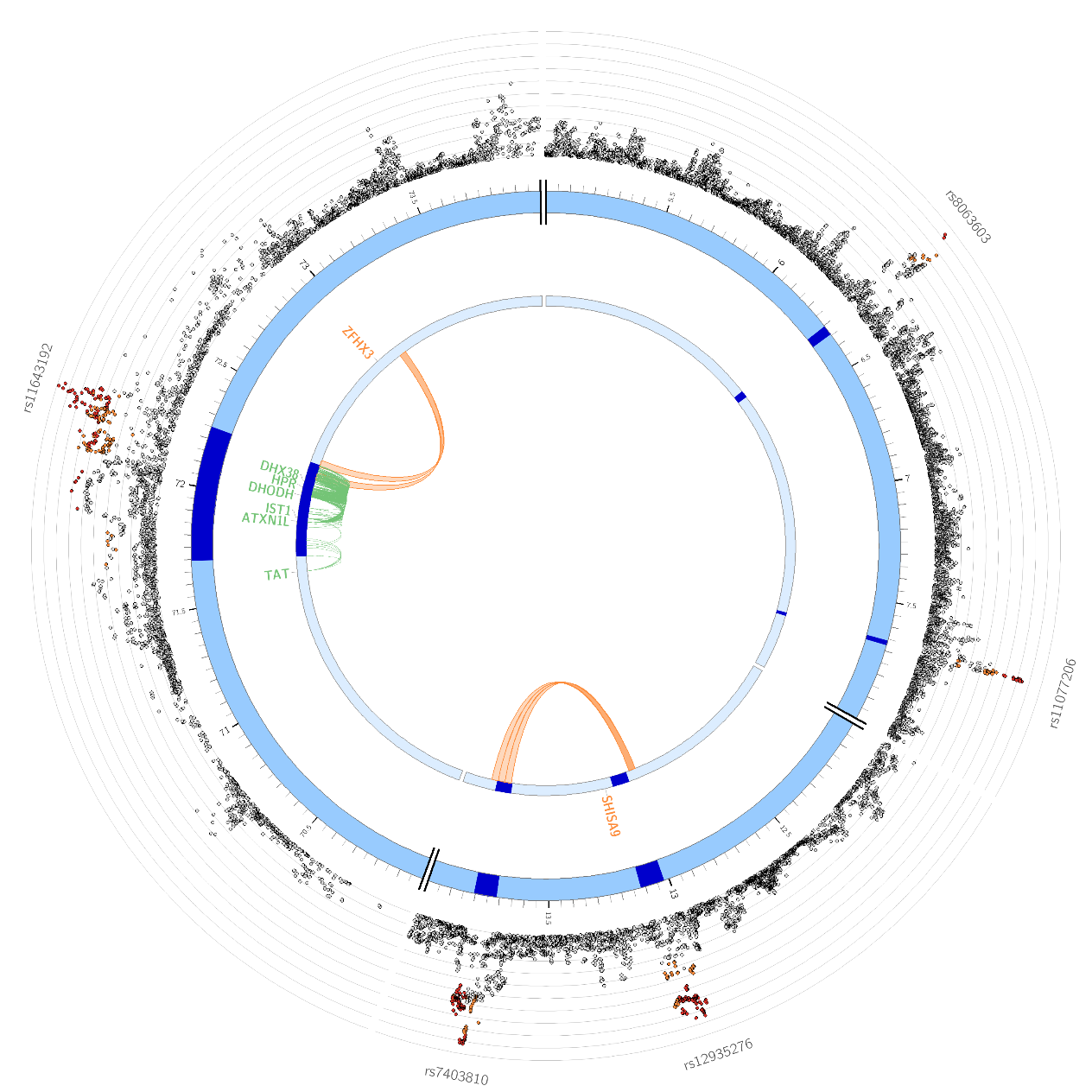

Supplementary Figure 28: Circos plot of significant loci from mood disorders (MOOD) on chromosome 16. Outer circle: Plot of individual variants in each locus, coloured by LD to labelled index variant (r2 0.8 -> 1, orange -> red). Middle ring: position of locus on chromosome, loci shown in blue. Inner ring: Links between loci and nearby genes, by eQTLs (green), chromatin contacts (orange) or both (red)

#### Supplementary Figure 29

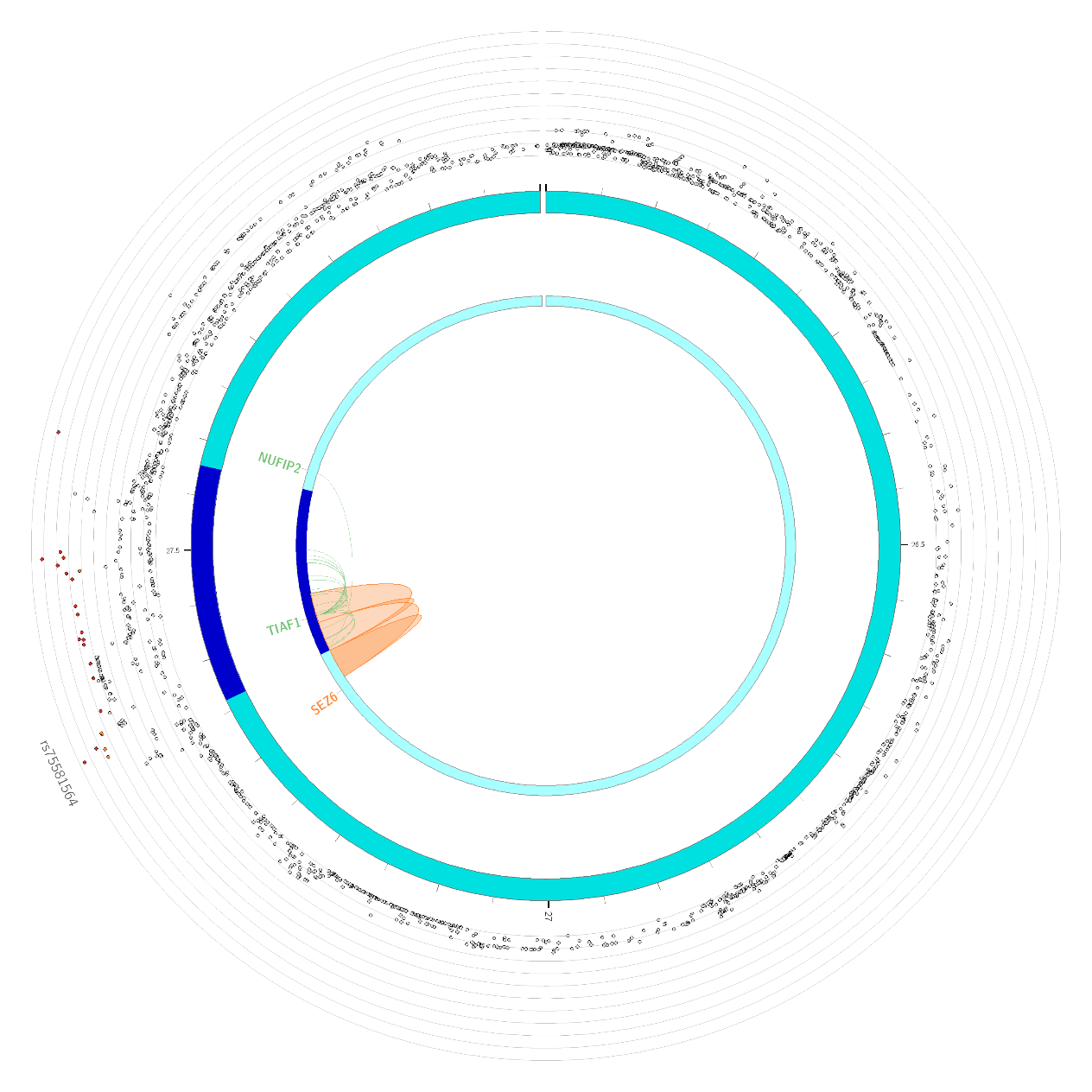

Supplementary Figure 29: Circos plot of significant loci from mood disorders (MOOD) on chromosome 17. Outer circle: Plot of individual variants in each locus, coloured by LD to labelled index variant (r2 0.8 -> 1, orange -> red). Middle ring: position of locus on chromosome, loci shown in blue. Inner ring: Links between loci and nearby genes, by eQTLs (green), chromatin contacts (orange) or both (red)

#### Supplementary Figure 30

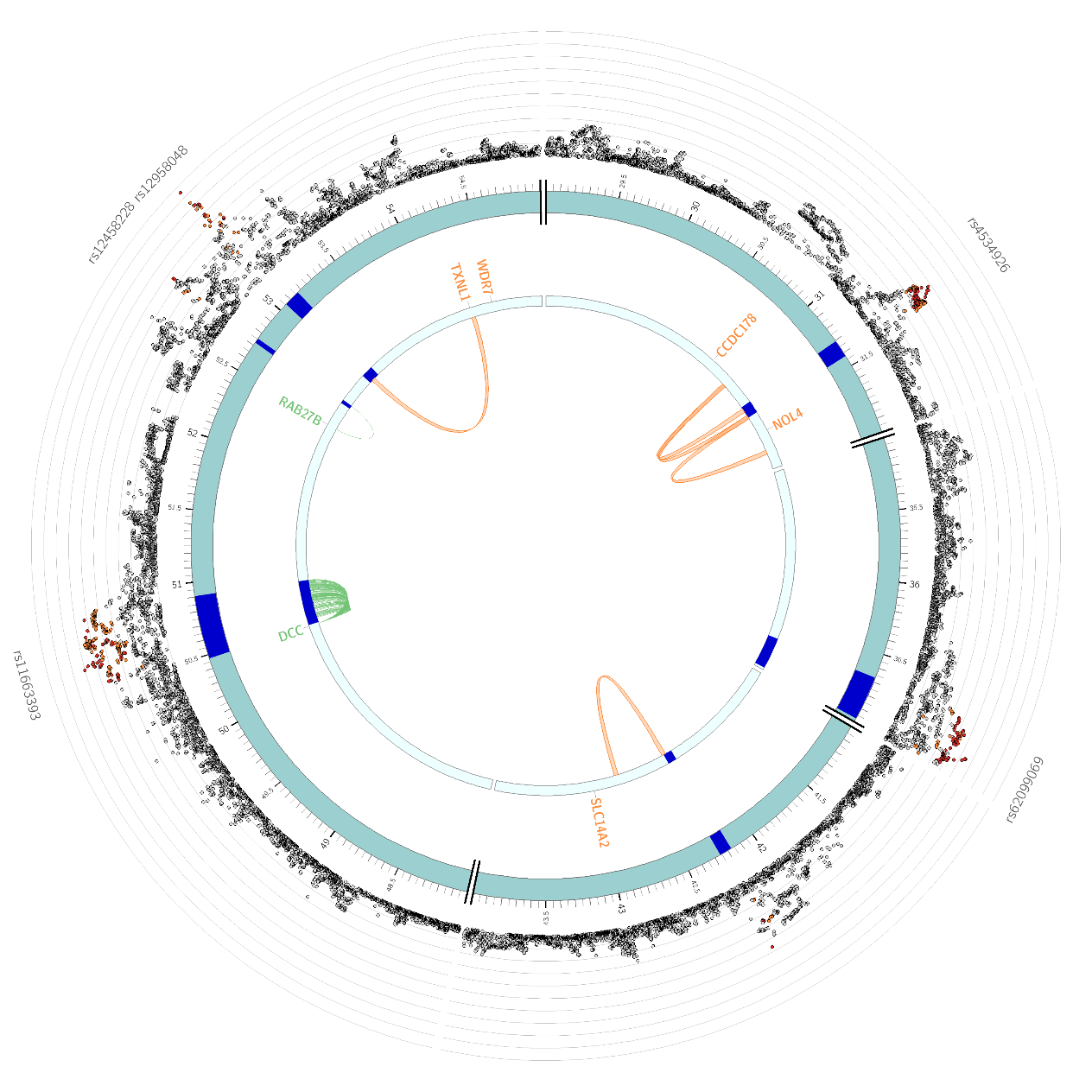

Supplementary Figure 30: Circos plot of significant loci from mood disorders (MOOD) on chromosome 18. Outer circle: Plot of individual variants in each locus, coloured by LD to labelled index variant (r2 0.8 -> 1, orange -> red). Middle ring: position of locus on chromosome, loci shown in blue. Inner ring: Links between loci and nearby genes, by eQTLs (green), chromatin contacts (orange) or both (red)

#### Supplementary Figure 31

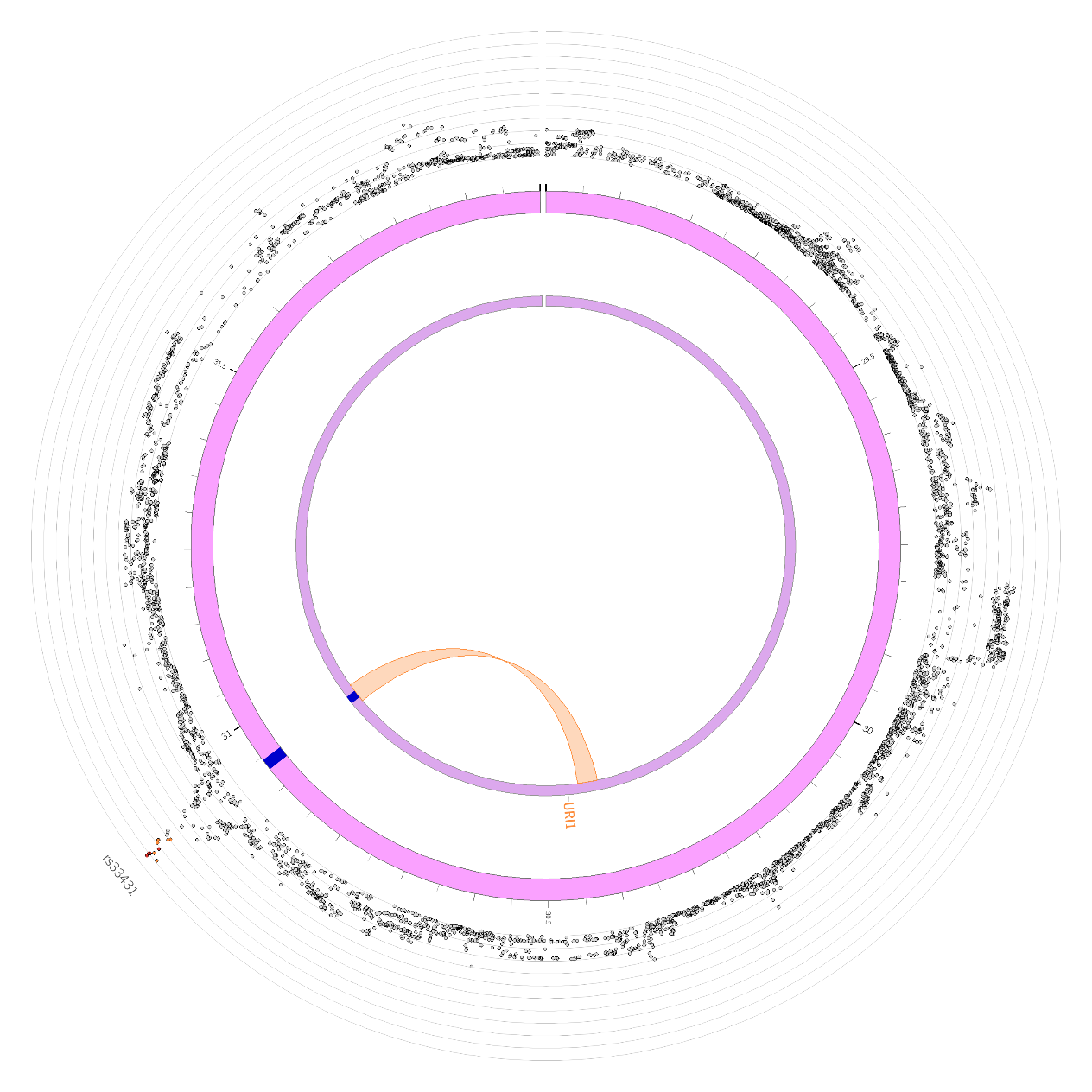

Supplementary Figure 31: Circos plot of significant loci from mood disorders (MOOD) on chromosome 19. Outer circle: Plot of individual variants in each locus, coloured by LD to labelled index variant (r2 0.8 -> 1, orange -> red). Middle ring: position of locus on chromosome, loci shown in blue. Inner ring: Links between loci and nearby genes, by eQTLs (green), chromatin contacts (orange) or both (red)

#### Supplementary Figure 32

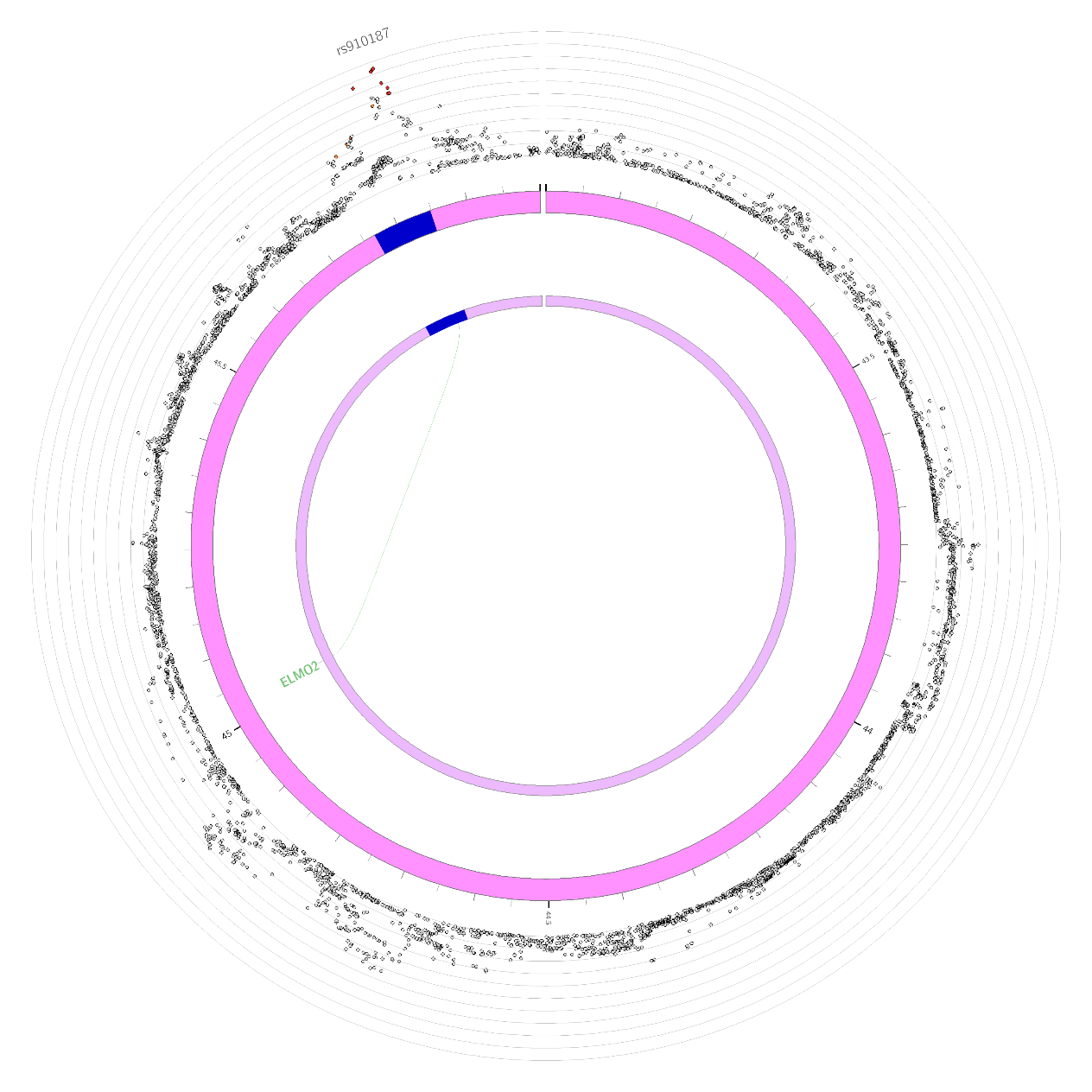

Supplementary Figure 32: Circos plot of significant loci from mood disorders (MOOD) on chromosome 20. Outer circle: Plot of individual variants in each locus, coloured by LD to labelled index variant (r2 0.8 -> 1, orange -> red). Middle ring: position of locus on chromosome, loci shown in blue. Inner ring: Links between loci and nearby genes, by eQTLs (green), chromatin contacts (orange) or both (red)

#### Supplementary Figure 33

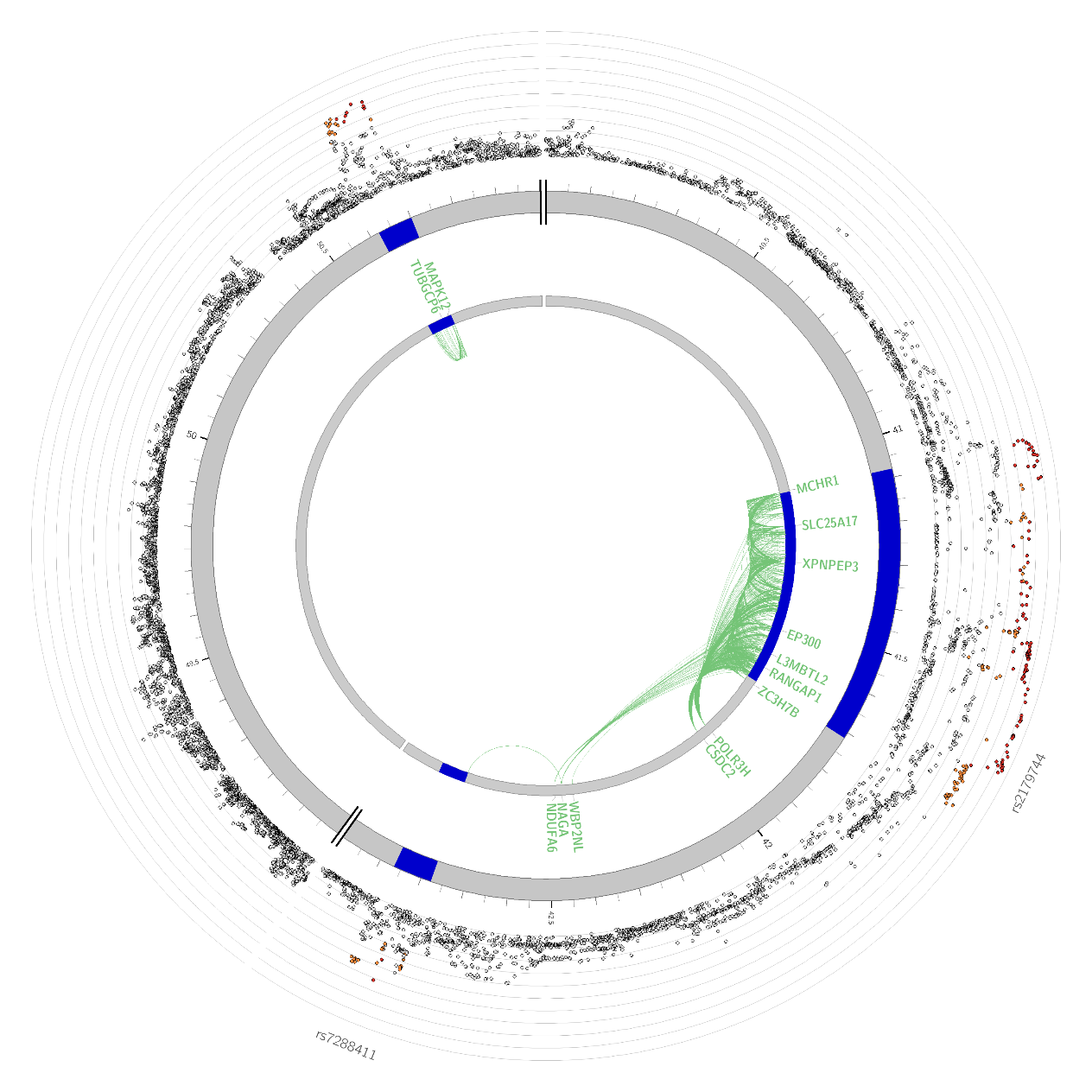

Supplementary Figure 33: Circos plot of significant loci from mood disorders (MOOD) on chromosome 22 (Note – there are no significant loci present on chromosome 21, so no circos plot is required). Outer circle: Plot of individual variants in each locus, coloured by LD to labelled index variant (r2 0.8 -> 1, orange -> red). Middle ring: position of locus on chromosome, loci shown in blue. Inner ring: Links between loci and nearby genes, by eQTLs (green), chromatin contacts (orange) or both (red)

#### Supplementary Figure 34

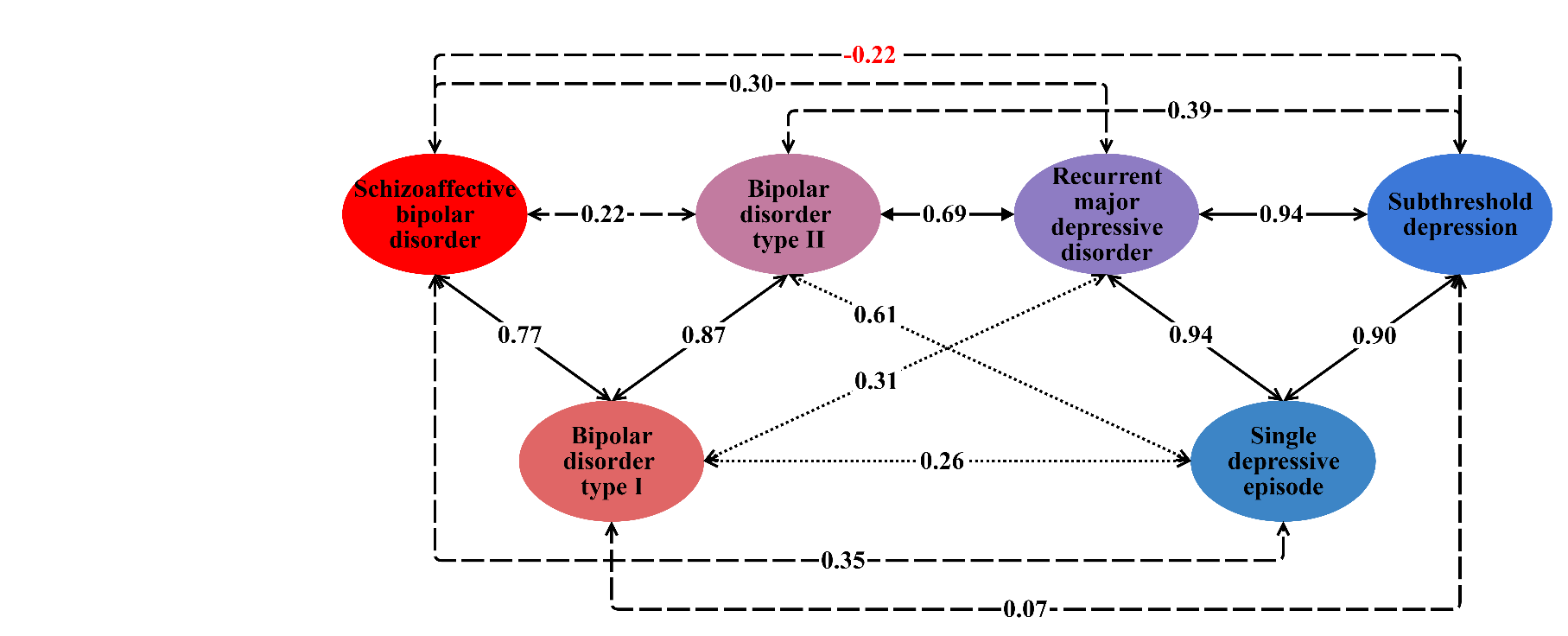

Supplementary Figure 34: Genetic correlations across the mood disorder spectrum, with all paths. Arrows labels show genetic correlations. Solid arrows represent genetic correlations significantly different from 0 and not significantly different from 1.
Dotted arrows represent genetic correlations significantly different from 0 and from 1.
Dashed arrows represent genetic correlations not significantly different from 0.
Significance in both cases means p < 0.00333 (Bonferroni correction for 15 tests).

#### Supplementary Figure 35

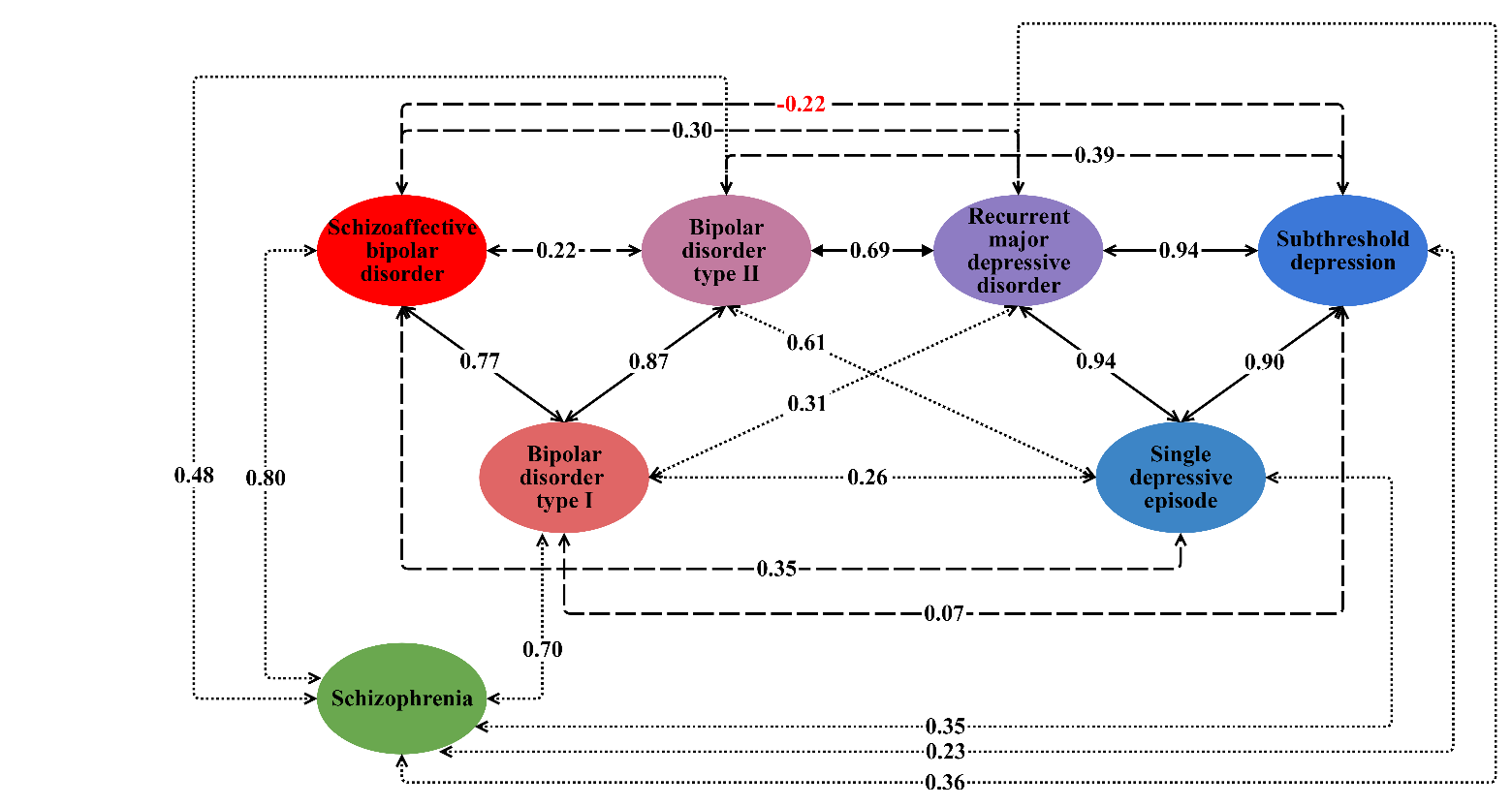

Supplementary Figure 35: Genetic correlations across the mood disorder spectrum with all paths (as Supplementary Figure 34), in the context of genetic correlations with schizophrenia. Arrows labels show genetic correlations. Solid arrows represent genetic correlations significantly different from 0 and not significantly different from 1.
Dotted arrows represent genetic correlations significantly different from 0 and from 1.
Dashed arrows represent genetic correlations not significantly different from 0.
For consistency with Supplementary Figure 31, significance in both cases means p < 0.00333 (Bonferroni correction for 15 tests).

#### Supplementary Figure 36

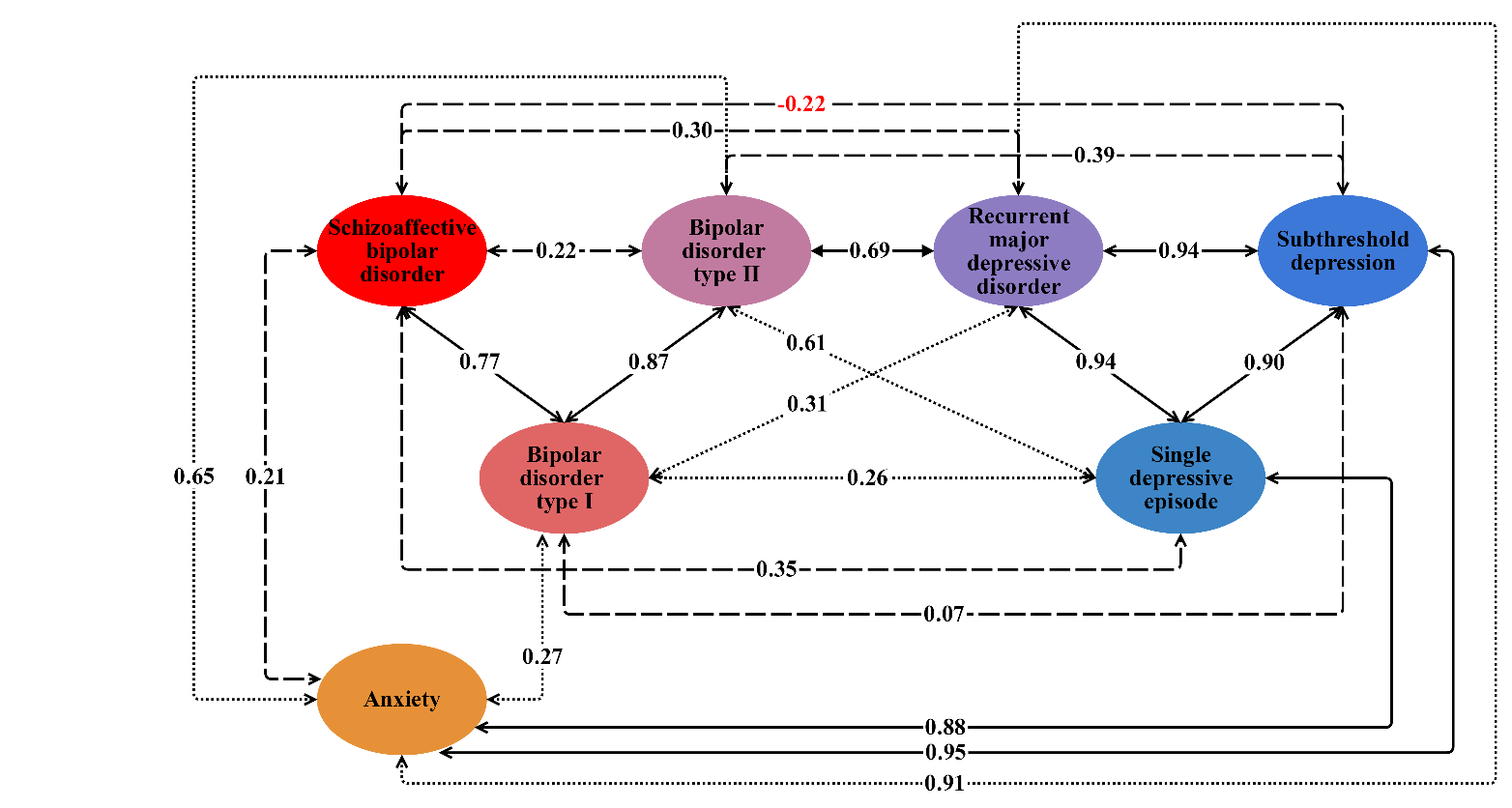

Supplementary Figure 36: Genetic correlations across the mood disorder spectrum with all paths (as Supplementary Figure 34), in the context of genetic correlations with anxiety. Arrows labels show genetic correlations. Solid arrows represent genetic correlations significantly different from 0 and not significantly different from 1.
Dotted arrows represent genetic correlations significantly different from 0 and from 1.
Dashed arrows represent genetic correlations not significantly different from 0.
For consistency with Supplementary Figure 31, significance in both cases means p < 0.00333 (Bonferroni correction for 15 tests).

#### Supplementary Figure 37

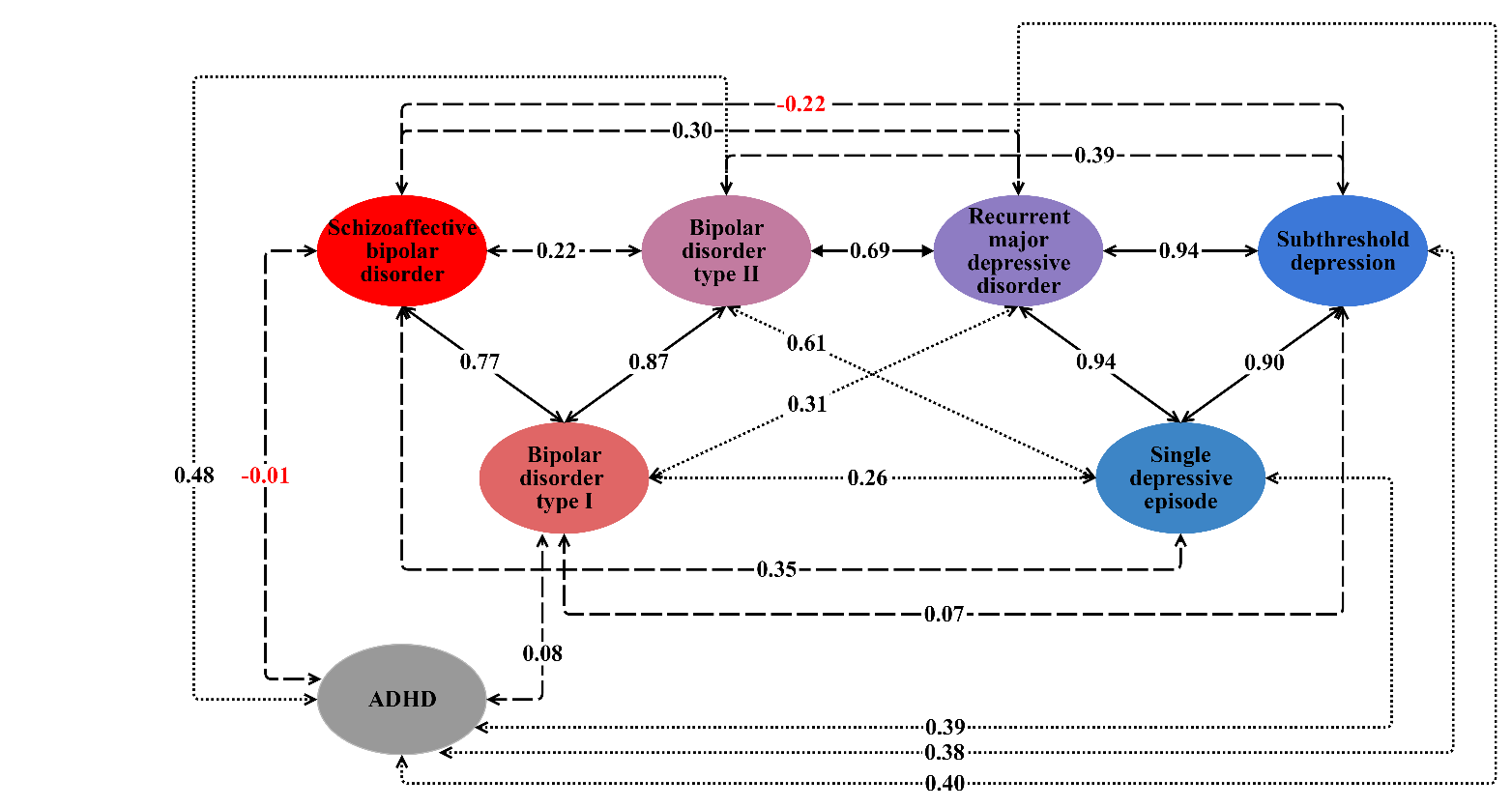

Supplementary Figure 37: Genetic correlations across the mood disorder spectrum with all paths (as Supplementary Figure 34), in the context of genetic correlations with ADHD. Arrows labels show genetic correlations. Solid arrows represent genetic correlations significantly different from 0 and not significantly different from 1.
Dotted arrows represent genetic correlations significantly different from 0 and from 1.
Dashed arrows represent genetic correlations not significantly different from 0.
For consistency with Supplementary Figure 31, significance in both cases means p < 0.00333 (Bonferroni correction for 15 tests).

#### Supplementary Figure 38

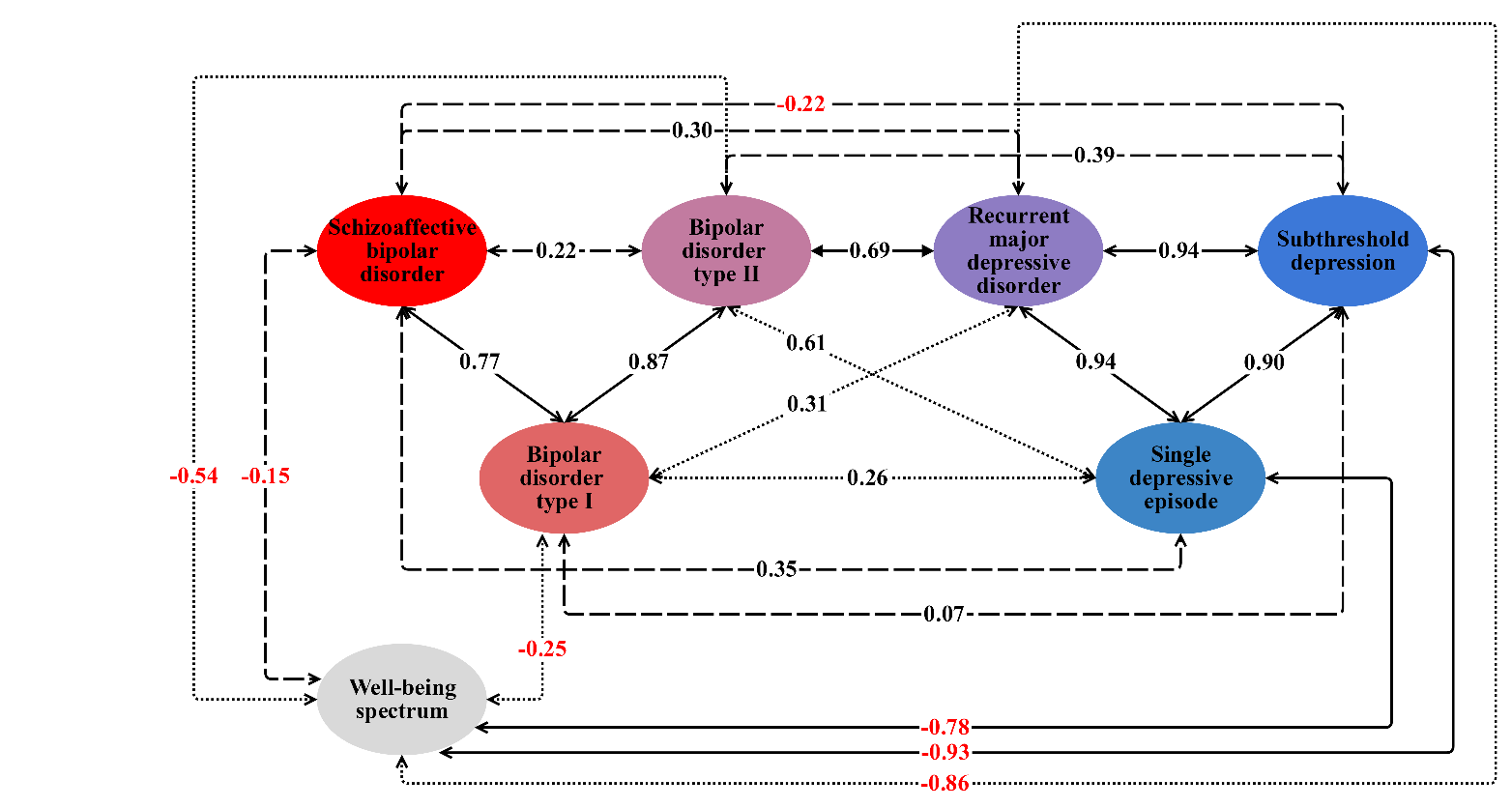

Supplementary Figure 38: Genetic correlations across the mood disorder spectrum with all paths (as Supplementary Figure 34), in the context of genetic correlations with the well-being spectrum. Arrows labels show genetic correlations. Solid arrows represent genetic correlations significantly different from 0 and not significantly different from 1.
Dotted arrows represent genetic correlations significantly different from 0 and from 1.
Dashed arrows represent genetic correlations not significantly different from 0.
For consistency with Supplementary Figure 31, significance in both cases means p < 0.00333 (Bonferroni correction for 15 tests).

#### Supplementary Figure 39

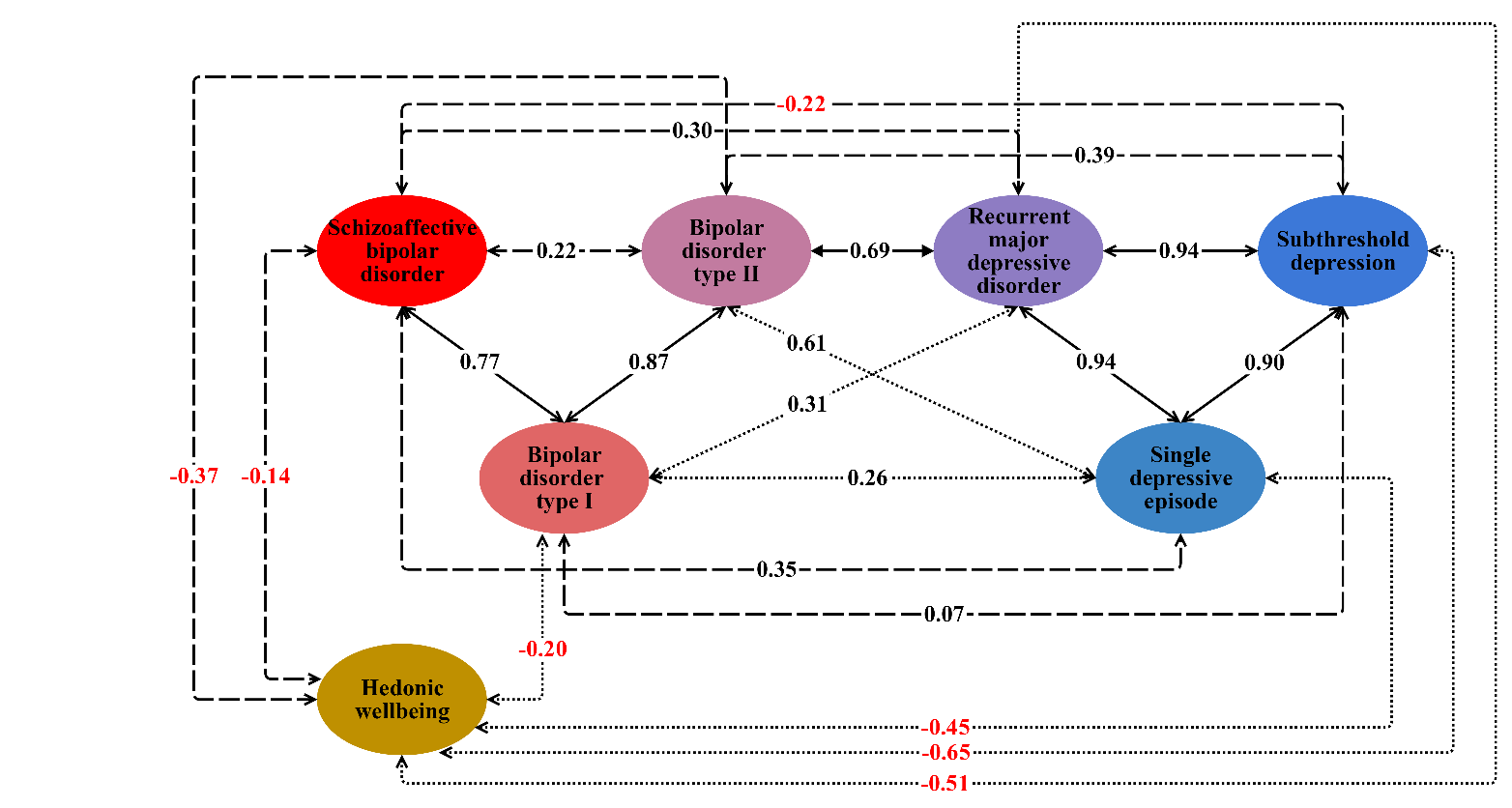

Supplementary Figure 39: Genetic correlations across the mood disorder spectrum with all paths (as Supplementary Figure 34), in the context of genetic correlations with hedonic well-being. Arrows labels show genetic correlations. Solid arrows represent genetic correlations significantly different from 0 and not significantly different from 1.
Dotted arrows represent genetic correlations significantly different from 0 and from 1.
Dashed arrows represent genetic correlations not significantly different from 0.
For consistency with Supplementary Figure 31, significance in both cases means p < 0.00333 (Bonferroni correction for 15 tests).

#### Supplementary Figure 40

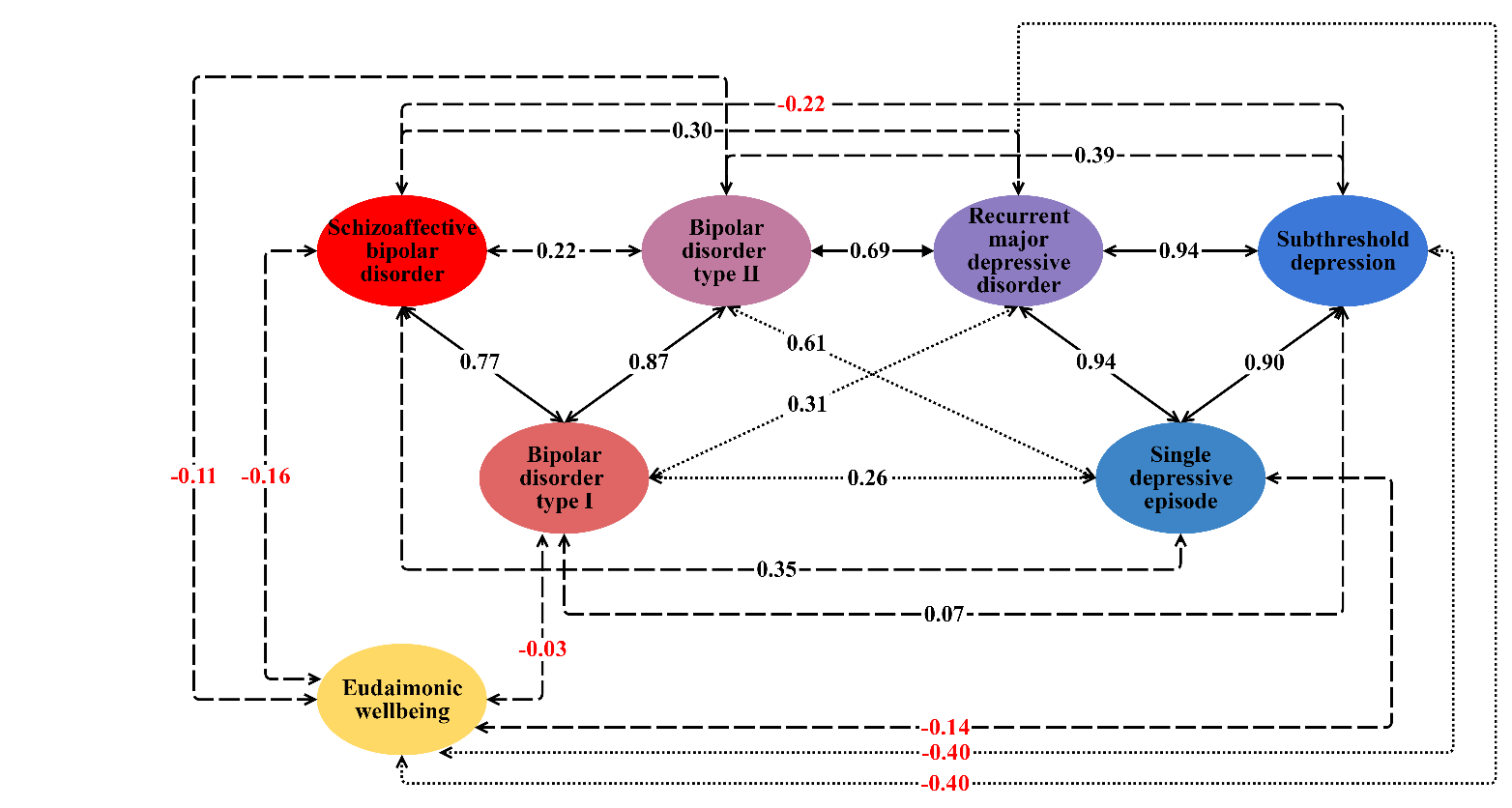

Supplementary Figure 40: Genetic correlations across the mood disorder spectrum with all paths (as Supplementary Figure 34), in the context of genetic correlations with eudaimonic well-being. Arrows labels show genetic correlations. Solid arrows represent genetic correlations significantly different from 0 and not significantly different from 1.
Dotted arrows represent genetic correlations significantly different from 0 and from 1.
Dashed arrows represent genetic correlations not significantly different from 0.
For consistency with Supplementary Figure 31, significance in both cases means p < 0.00333 (Bonferroni correction for 15 tests).

#### Supplementary Figure 41

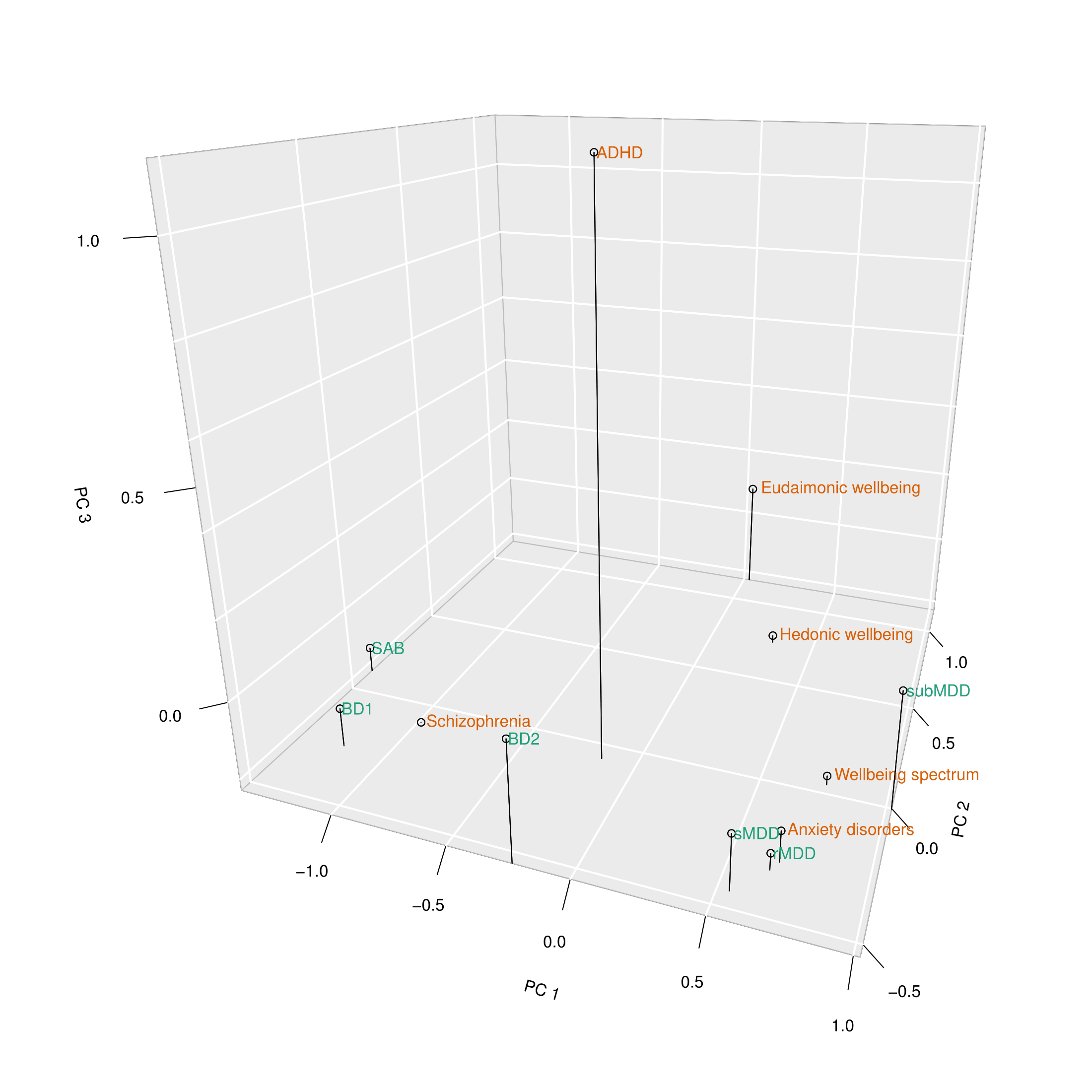

Supplementary Figure 41: Three-dimensional scatterplot of the first three principal components of the genetic correlation matrix of the mood disorder subtypes (green labels) with six external GWAS (orange labels). Principal component (PC) 1 separates BD1, SAB and schizophrenia from the depressive disorders, with BD2 and ADHD intermediate. PC2 separates eudaimonic and hedonic wellbeing from the other phenotypes. PC3 separates ADHD from the other phenotypes.

#### Supplementary Figure 42

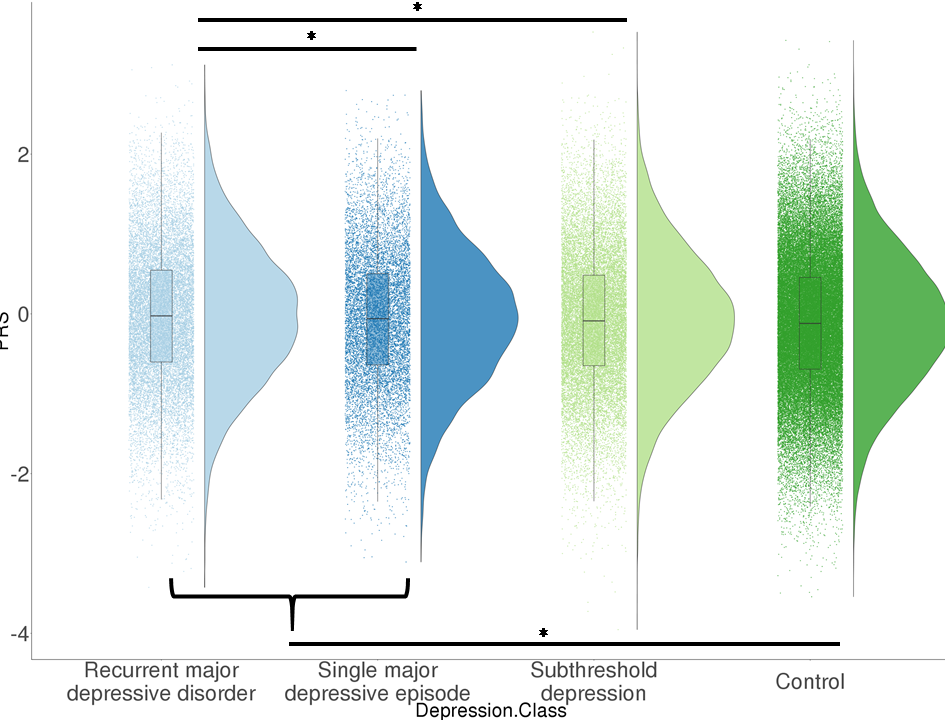

Supplementary Figure 42: Distribution of polygenic risk scores derived from PGC BD (pThresh = 0.2) in individuals with recurrent major depressive disorder (rMDD cases), single episode major depressive disorder (sMDD cases), subthreshold depression (subMDD pseudo-cases) and controls. Black bars - significant differences in means (empirical p < 0.05)

#### Supplementary Figure 43

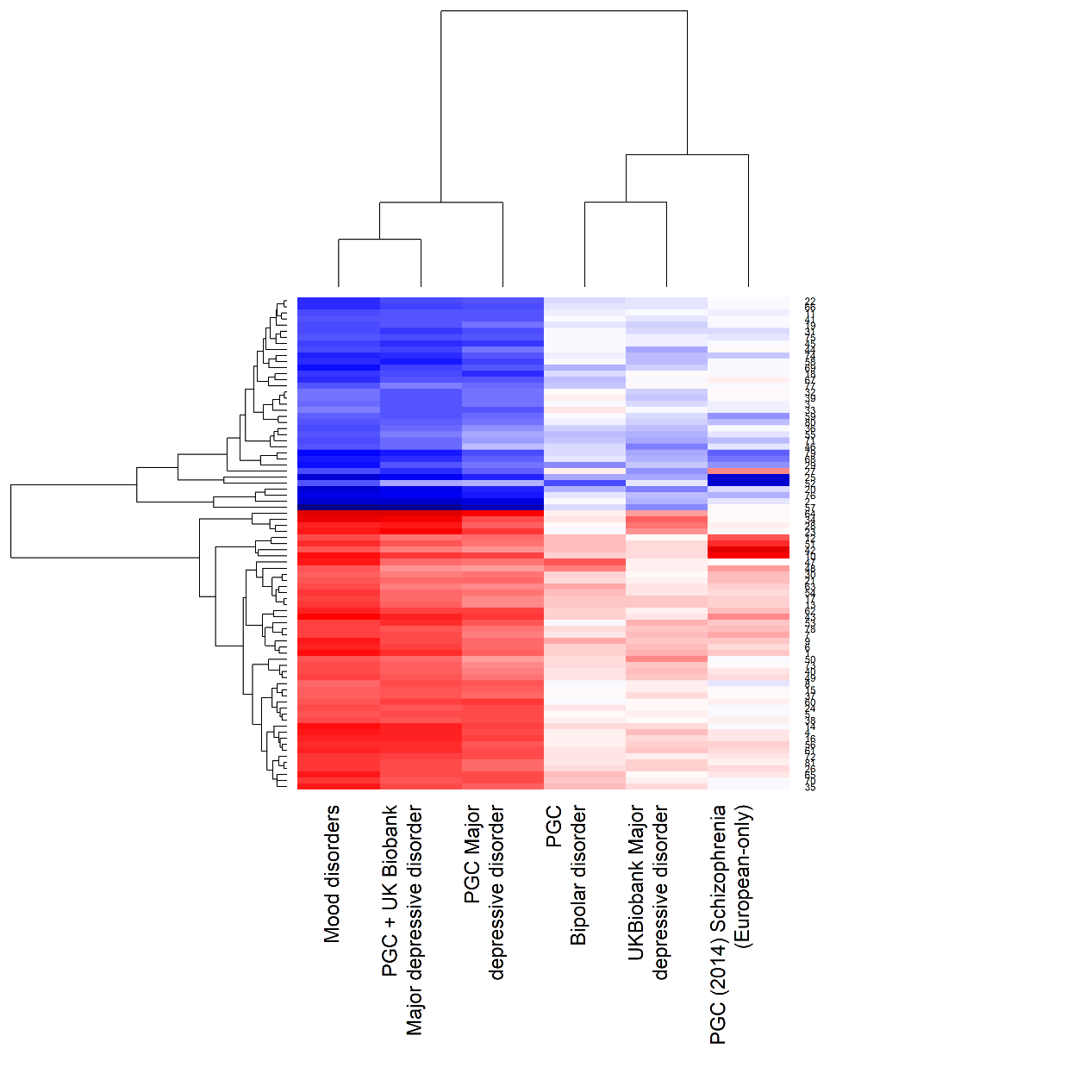

Supplementary Figure 43: Hierarchical clustering of the significant loci from the mood disorders meta-analysis (MOOD) with the same loci from PGC MDD, UKB MDD, PGC BD, and SCZ. Blue = positive direction of effect. Red = negative direction of effect.

#### Supplementary Figure 44

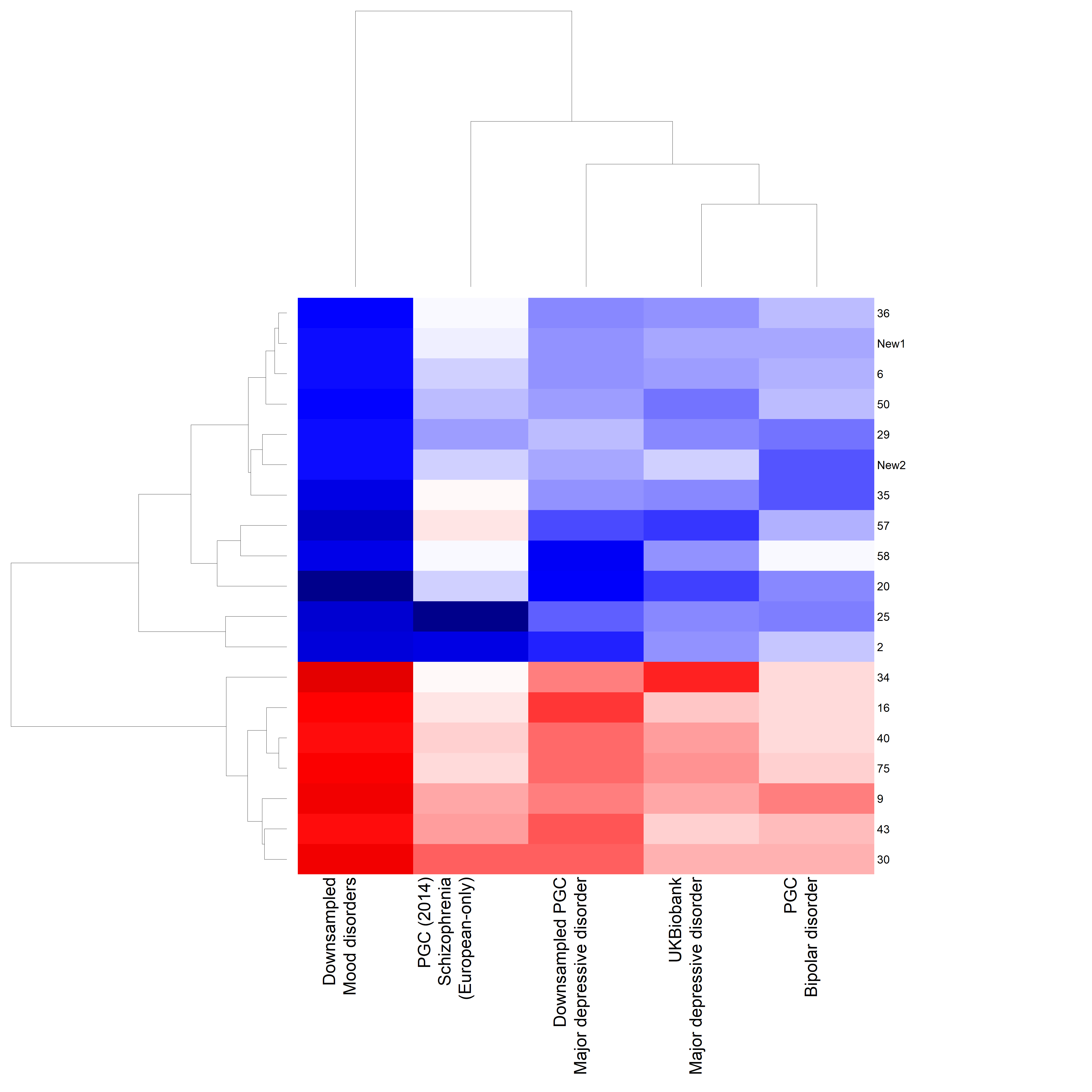

Supplementary Figure 44: Hierarchical clustering of the significant loci from the down-sampled mood disorders meta-analysis (downsampled MOOD) with the same loci from downsampled PGC MDD, UKB MDD, PGC BD, and SCZ. Blue = positive direction of effect. Red = negative direction of effect.
